# Chloroplast competition is controlled by lipid biosynthesis in evening primroses

**DOI:** 10.1101/330100

**Authors:** Johanna Sobanski, Patrick Giavalisco, Axel Fischer, Julia Kreiner, Dirk Walther, Mark Aurel Schöttler, Tommaso Pellizzer, Hieronim Golczyk, Toshihiro Obata, Ralph Bock, Barbara B. Sears, Stephan Greiner

## Abstract

In most eukaryotes, organellar genomes are transmitted preferentially by the mother, but molecular mechanisms and evolutionary forces underlying this fundamental biological principle are far from understood. It is believed that biparental inheritance promotes competition between the cytoplasmic organelles and allows the spread of so-called selfish cytoplasmic elements. Those can be, for example, fast replicating or aggressive chloroplasts (plastids) that are incompatible with the hybrid nuclear genome and therefore maladaptive. Here we show that the ability of plastids to compete against each other is a metabolic phenotype determined by extremely rapidly evolving genes in the plastid genome of the evening primrose *Oenothera*. Repeats in the regulatory region of *accD* (the plastid-encoded subunit of the acetyl-CoA carboxylase, which catalyzes the first and rate limiting step of lipid biosynthesis), as well as in *ycf2* (a giant reading frame of still unknown function), are responsible for the differences in competitive behavior of plastid genotypes. Polymorphisms in these genes influence lipid synthesis and most likely profiles of the plastid envelope membrane. These in turn determine plastid division and/or turn-over rates and hence competitiveness. This work uncovers cytoplasmic drive loci controlling the outcome of biparental chloroplast transmission. Here, they define the mode of chloroplast inheritance, since plastid competitiveness can result in uniparental inheritance (through elimination of the “weak” plastid) or biparental inheritance (when two similarly “strong” plastids are transmitted).

**Significance statement:** Plastids and mitochondria are usually uniparentally inherited, typically maternally. When the DNA-containing organelles are transmitted to the progeny by both parents, evolutionary theory predicts that the maternal and paternal organelles will compete in the hybrid. As their genomes do not undergo sexual recombination, one organelle will “try” to outcompete the other, thus favoring the evolution and spread of aggressive cytoplasms. The investigations described here in the evening primrose, a model species for biparental plastid transmission, have discovered that chloroplast competition is a metabolic phenotype. It is conferred by rapidly evolving genes that are encoded on the chloroplast genome and control lipid biosynthesis. Due to their high mutation rate these loci can evolve and become fixed in a population very quickly.

## Introduction

Most organelle genomes are inherited from the mother (1, 2), but biparental transmission of plastids has evolved independently multiple times. About 20% of all angiosperms contain chloroplasts in the pollen generative cell indicating at least a potential for biparental transmission (2-4). Whereas reasons for this are controversial (2, 5-9), a genetic consequence of the biparental inheritance patterns is a genomic conflict between the two organelles. When organelles are transmitted to the progeny by both parents, they compete for cellular resources. Since the plastids do not fuse and hence their genomes do not undergo sexual recombination, selection will favor the organelle genome of the competitively superior plastid (2, 10-13). In a population, the ensuing arms race can lead to evolution and spread of selfish or aggressive cytoplasmic elements that potentially could harm the host cell. The mechanisms and molecular factors underlying this phenomenon are elusive, yet there is solid evidence for a widespread presence of competing organelles (2). Heteroplasmic cells can be created from cell fusion events, mutation of organelle DNA or sexual crosses (14-17), however, only very few cases have been studied in some detail in model organisms. One them is *Drosophila*, where mitochondrial competition experiments can be set up via cytoplasmic microinjections (18, 19). Another is the evening primrose (genus *Oenothera*) (20, 21). This plant genus is certainly the model organism of choice to study chloroplast competition; the original theory of selfish cytoplasmic elements is based on evening primrose genetics (10): In the *Oenothera*, biparental plastid inheritance is the rule (22) and the system is a prime example of naturally occurring aggressive chloroplasts (2, 10). Based on extensive crossing studies, five genetically distinguishable chloroplast genome (plastome) types were shown to exist. Those were designated by Roman numerals (I to V) and grouped into three classes according to their inheritance strength or assertiveness rates in crosses (strong: plastomes I and III, intermediate: plastome II, weak: plastomes IV and V), reflecting their ability to outcompete a second chloroplast genome in the F1 generation upon biparental transmission (20, 22, 23). The plastome types were initially identified based on their (in)compatibility with certain nuclear genomes (24, 25). Strong plastomes provide hitchhiking opportunities for loci that result in incompatible or maladaptive chloroplasts, and these would be viewed as selfish cytoplasmic elements (2, 10). It was further shown that the loci, which determine the differences in competitive ability of the chloroplast, are encoded by the chloroplast genome itself (20, 22, 23).

## Results

To pinpoint the underlying genetic determinants, we developed a novel association mapping approach that correlates local sequence divergence to a phenotype. We analyzed 14 complete chloroplast genomes from *Oenothera* wild type lines, whose inheritance strength had been previously classified in exhaustive crossing analyses (20, 22, 26). This enabled us to correlate the experimentally determined inheritance strengths to sequence divergence in a given alignment window (Materials and Methods; SI Appendix, Text for details). Both the Pearson’s and Spearman’s applied correlation metrics associated the genetic determinants of inheritance strength with four major sites: the regulatory region of the fatty acid biosynthesis gene *accD* (promoter, 5’-UTR and protein N-terminus), the *origin of replication B (oriB), ycf1* and *ycf2* (including its promoter/5’-UTR; Fig. 1; SI Appendix, Fig S1, Dataset S1, and SI Text). The *ycf1* and *ycf2* genes are two open reading frames (ORFs) of unknown function; *ycf1* (or *tic214*) has tentatively been identified as an essential part of the chloroplast protein import machinery (27), but this function has been questioned (28). The *accD* gene encodes the beta-carboxyltransferase subunit of the plastid-localized plant heteromeric acetyl-CoA carboxylase (ACCase). The enzyme is responsible for catalyzing the initial tightly-regulated and rate-limiting step in fatty acid biosynthesis. The other three required ACCase subunits, alpha-carboxyltransferase, biotin-carboxyl carrier protein, and biotin carboxylase, are encoded by nuclear genes (29). The polymorphisms detected through our correlation mapping represent large insertions/deletions (indels), which are in-frame in all coding sequences (Fig. 1B; SI Appendix, Figs. S2-S6, Dataset S2, and SI Text). Several other polymorphisms in intergenic spacers of photosynthesis and/or chloroplast translation genes were correlated with plastome assertiveness rates (see Fig. 1A; SI Appendix, Dataset S1 and SI Text for details). However, both of those gene classes are unlikely to affect chloroplast inheritance, since other studies have shown that mutations in genes that result in chlorophyll-deficient chloroplasts do not alter their inheritance strengths in the evening primrose (20, 30, 31) (SI Appendix, SI Text). By contrast, the large ORFs of unknown function, the origins of replication and a central gene in lipid metabolism such as *accD* (29), are serious candidates to encode factors involved in chloroplast competition.

**Fig. 1.**
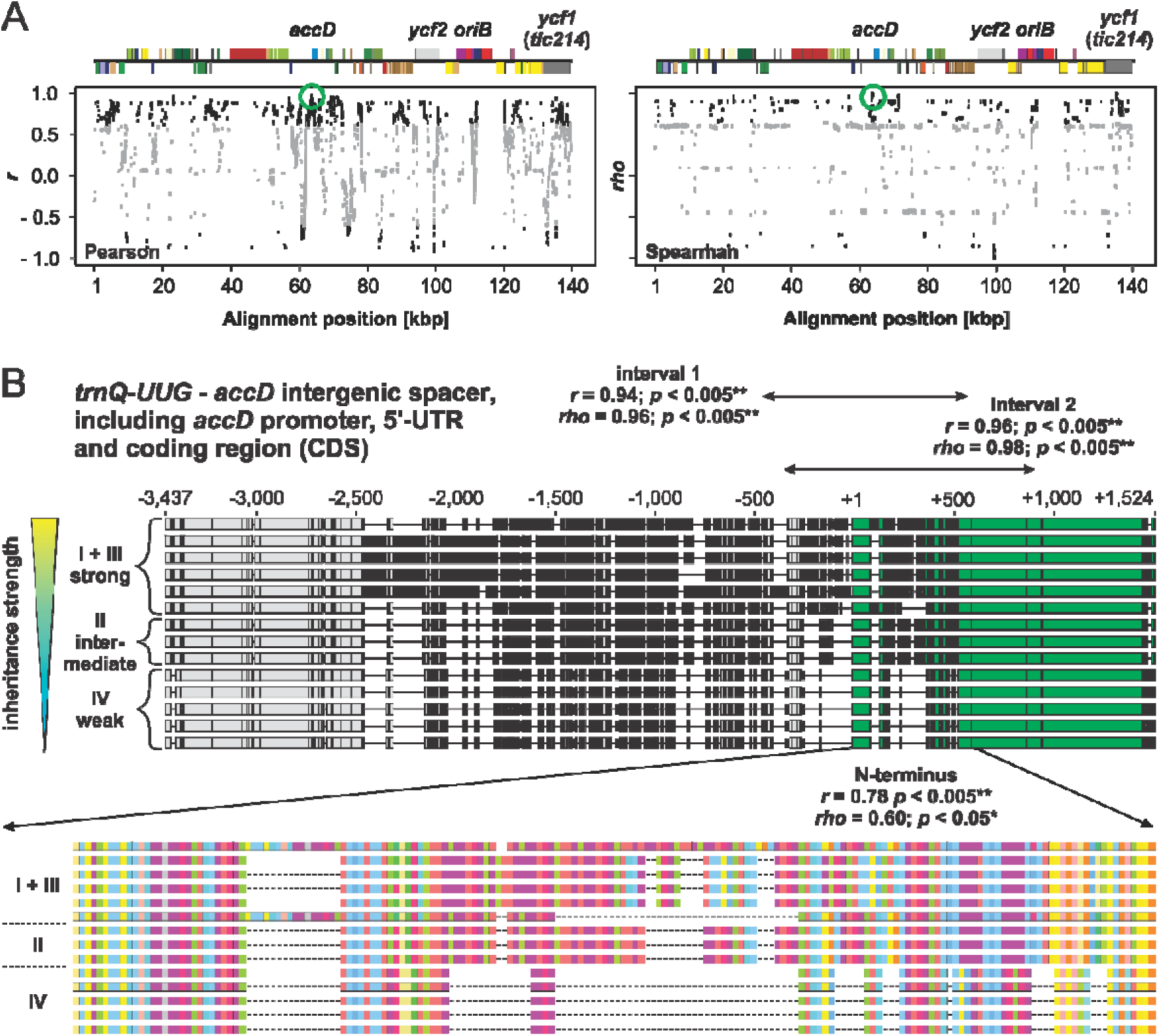
Correlation mapping to identify chloroplast loci for inheritance strength in the wild type chloroplast genomes. (*A*) Spearman and Pearson correlation to inheritance strength plotted against alignment windows of the wild type plastomes. Relevant genes or loci with significant correlation are indicated in the linear plastome maps above. The region displayed in panel (B) is highlighted by green circles. Significant correlations (*p* < 0.05) are shown in black. Correlations to *k*-means classes are shown. For details, see Materials and Methods, SI Appendix (SI Text) and Main Text. (B) Correlation to inheritance strength at the *accD* region in the wild type plastomes. Individual sequences are sorted according to their competitive ability. Polymorphic regions are indicated in black, thin lines represent gaps mostly resulting from deletions. Upper panel: Alignment of the *trnQ* - *UUG* - *accD* intergenic spacer (−3,437 to −1) and the *accD* gene, including promoter, 5’-UTR and coding sequence (CDS). The *accD* CDS, starting from +1, is highlighted in green. Regions marked by “interval 1” and “interval 2” display nearly absolute correlation to inheritance strength (see SI Appendix, SI Text and Dataset S1 for details). Note that these sequence intervals span the promotor region, the 5’-UTR and the 5’-end of *accD*. All three are considered to play a regulatory role (29, 75-78). Lower panel: Amino acid sequence of the *accD* N-terminus and correlation to inheritance strength. Colors indicated different amino acids. Most variation in the sequence is conferred by repeats encoding glutamic acid-rich domains marked in purple (also see SI Appendix, SI Text).

Due to the lack of measurable sexual recombination in seed plant chloroplast genomes (2), our correlation mapping method first established associations of polymorphic loci that are fixed in the slow plastome type IV with weak inheritance strength, regardless of their functional relevance (SI Appendix, SI Text). To test whether those loci affect inheritance strength, we conducted a genetic screen for weak chloroplast mutants derived from the strong chloroplast genome I by employing a *plastome mutator* (*pm*) allele (Materials and Methods). The *pm*-based mutagenesis approach yields indels in repetitive regions (32), similar to those identified by our association mapping (see SI Appendix, SI Text for details). This led to the isolation of 24 plastome I variants with altered inheritance strength (Fig. 2; SI Appendix, SI Text). Since we selected for green and photosynthetically competent chloroplasts in the mutagenesis (Materials and Methods for details), none of those variants differed from the wild type in their photosynthetic parameters, chloroplast size or chloroplast volume per cell. The plants with the variant plastomes did not display any growth phenotype (SI Appendix, Figs. S7-S9 and SI Text). Sequence analysis of 18 variants, spanning the range of observed variation in inheritance strength, revealed an average of seven mutation events per variant. The analysis included one additional variant (VC1) with more background mutations, which resulted from its isolation after being under the mutagenic action of the *plastome mutator* for several generations (Materials and Methods; SI Appendix, SI Text for details). Most of the *pm*-induced mutations are composed of single base pair indels at oligo(N) stretches in intergenic regions, i.e. not of functional relevance, or larger in-frame indels at the highly repetitive sites of *accD*, *ycf1*, *ycf2* or the *oriB* (SI Appendix, Figs. S2-S6, Table S1, and Dataset S2). Correlation analysis to inheritance strengths at these sites confirmed the relevance of *accD* and *ycf2* in chloroplast inheritance (*r* = 0.72, *p* = 0.05, for the 5’-end of AccD; *r* = 0.91, *p* < 0.0005 for the promotor/5’-UTR of ycf2 and *r* = 0.70, *p* < 0.005, for a mutated site in *Ycf2*; Fig. 3; SI Appendix, Figs. S2-S6 and S10, and SI Text). These findings were confirmed by two additional very weak mutants derived from the strong plastome III (Materials and Methods; SI Appendix, SI Text). Based on the full chloroplast genome sequences of these lines (SI Appendix, Dataset S2), it appeared that the promotor/5’-UTR of *accD* is affected in one of the lines. In addition, in both lines the *ycf2* gene is most heavily mutated when compared to all other (weak) materials sequenced so far. This strongly advocates for *ycf2* being involved in chloroplast competition (SI Appendix, Table S2). The very weak variants of plastome III, as well as correlation analysis of oriB and *ycf1* in the plastome I variants did not support an involvement of these regions in the inheritance phenotype (Fig. 3; SI Appendix Figs. S6 and S10, Tables S1 and S2, and SI Text). Furthermore, the second replication origin (*oriA*) was found to be nearly identical within all sequenced wild type or mutant plastomes.

**Fig. 2.**
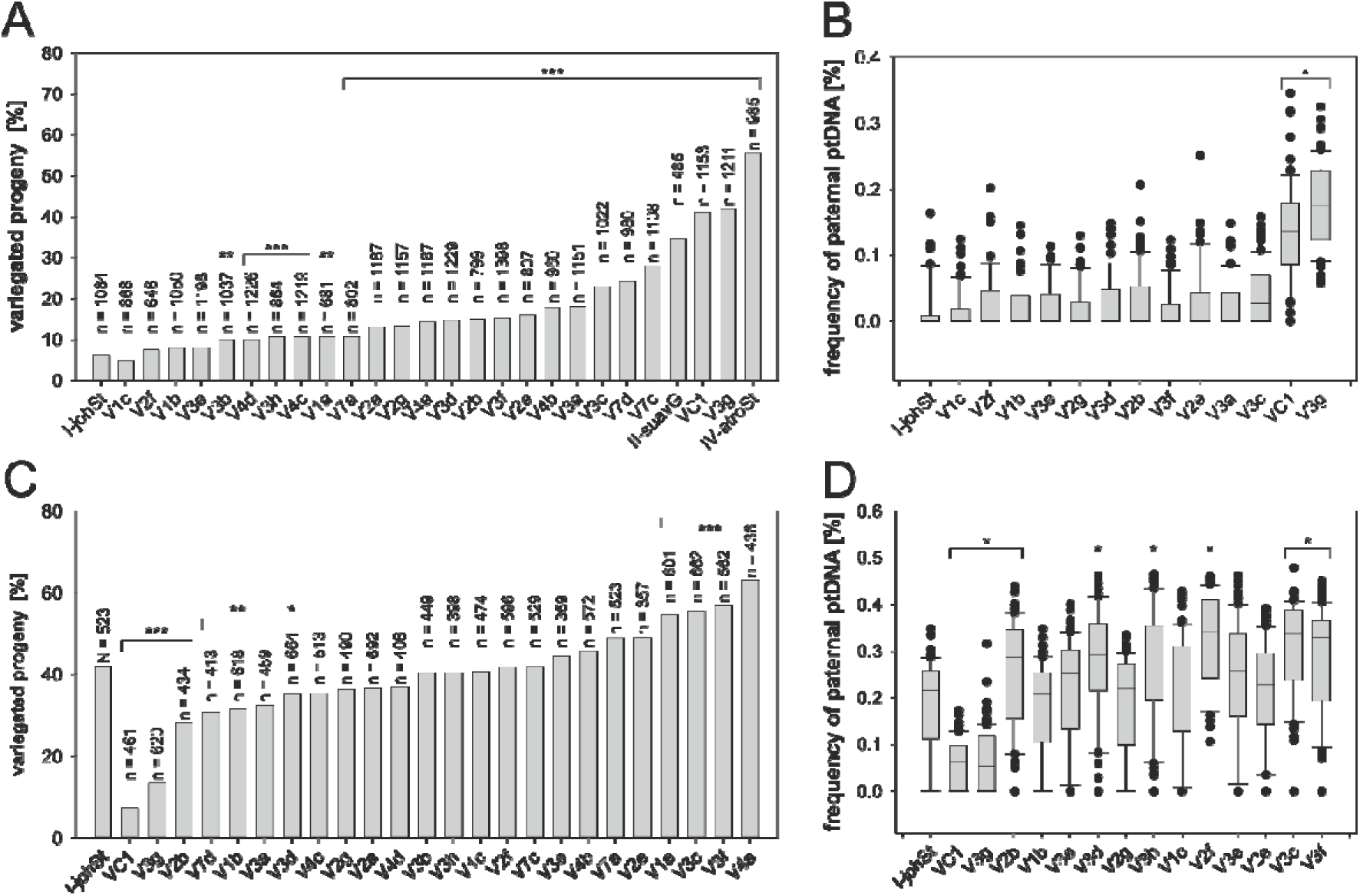
Transmission efficiencies of plastome I variants and the wild type plastomes I-johSt (strong), II-suavG (intermediate) and IV-atroSt (weak) as determined by crosses with I-chi/I-hookdV (strong) and IV-delta/IV-atroSt (weak). (*A,C*) Classical approach based on counting of variegated seedlings. (*B,D*) MassARRAY^®^ assay quantifying maternal and paternal ptDNA. Wild types and green variants were crossed (A) as the seed parent to I-chi as male parent and (*C*) crossed as the pollen parent to IV-delta as female parent. The X-axis depicts the average percentage of variegated progeny (= percentage biparental inheritance) obtained from three seasons (2013, 2014 and 2015). Fisher’s exact test determined significance of differences between variants and their progenitor I-johSt (*** p < 0.0001, ** *p* < 0.001, * *p* < 0.01). (*B*) Crosses of I-johSt and variants with I-hookdV as male and, (*D*) with IV-atroSt as female parent. Box-plots represent the transmission frequencies of the paternal plastomes measured by MassARRAY^®^. To account for significant differences to I-johSt Kruskal-Wallis, one-way ANOVA on ranks was performed (* *p* < 0.05). Due to the detection threshold of the MassARRAY^®^ (5-10%) most variants show the same or slightly decreased transmission efficiency as their wild type I-johSt (*B,D*). Only for the weak variants VC1 and V3g is the difference of the ratio of paternal and maternal ptDNA in the pool large enough to result in the detection of a significantly lower assertiveness rate in both crossing directions. Altogether, the classical approach using bleached chloroplast mutants gives more reliable results and allows a much finer discrimination of transmission efficiencies (cf. *A* vs. *B* and *C* vs. *D*). For details see SI Appendix, SI Text.

**Fig. 3.**
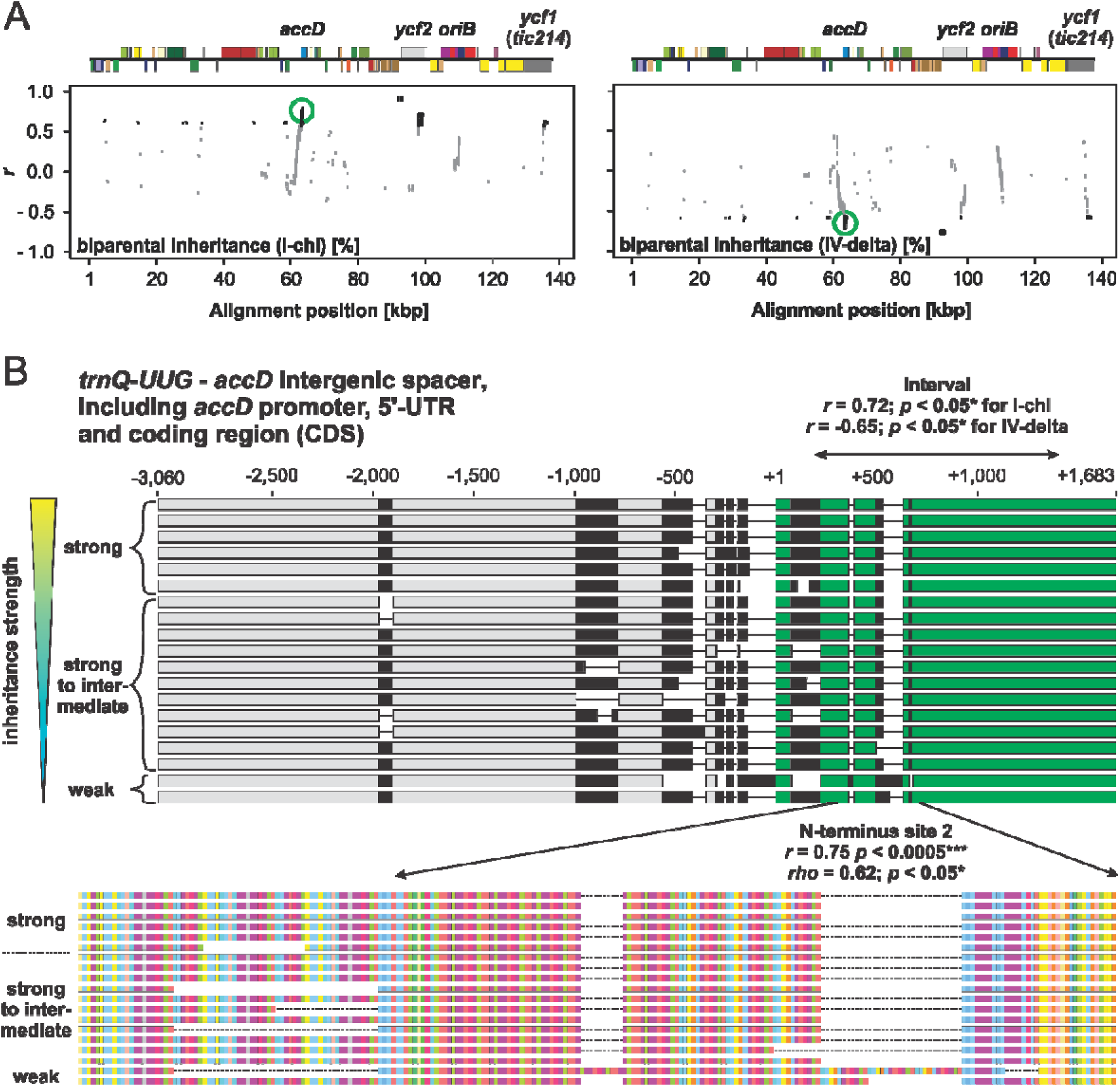
Correlation mapping to identify chloroplast loci for inheritance strength in the plastome I variants. (*A*) Pearson correlation to inheritance strength plotted against alignment windows of the variants’ plastomes. Relevant genes or loci with significant correlation are indicated in the linear plastome maps above. The region displayed in panel (*B*) is highlighted by green circles. Significant correlations (*p* < 0.05) are shown in black. Correlations to I-chi and IV-delta crosses (Fig. 2A,C) are shown. For details, see Materials and Methods, SI Appendix (SI Text) and Main Text. (B) Correlation to inheritance strength at the *accD* region in the variants’ plastomes. Individual sequences are sorted according to their competitive ability. Polymorphic regions are indicated in black, thin lines represent gaps mostly resulting from deletions. Upper panel: Alignment of the *trnQ-UUG* - *accD* intergenic spacer (−3,060 to −1) and the *accD* gene, including promoter, 5’-UTR and coding region (CDS). The *accD* CDS, starting from +1, is highlighted in green. The region marked by “interval” within the *accD* coding sequence displays high correlation to inheritance strength (see SI Appendix, SI Text and Dataset S1 for details). Lower panel: Amino acid sequence of the *accD* N-terminus and correlation to inheritance strength. Colors indicated different amino acids. Most variation in the sequence is conferred by repeats encoding glutamic acid-rich domains marked in purple (also see SI Appendix, SI Text).

These data argue against plastid DNA (ptDNA) replication *per se* being responsible for differences in chloroplast competitiveness. This conclusion is in line with previous analyses of the Oenothera replication origins, which had suggested that their variability does not correlate with the competitive strength of the plastids (33, 34) (also see SI Appendix, SI Text). We further confirmed this by determining the relative ptDNA amounts of chloroplasts with different inheritance strengths in a constant nuclear background. No significant variation of ptDNA amounts was observed over a developmental time course in these lines, thus excluding ptDNA stability and/or turnover as a potential mechanism (SI Appendix, Fig. S11 and SI Text). Moreover, no significant differences in nucleoid numbers per chloroplast nor in nucleoid morphology was observed, as judged by DAPI staining (SI Appendix, Figs. S12 and S13, SI Text).

Next, we conducted a more detailed analysis of *accD* and *ycf2*. In a constant nuclear background, the weak wild type plastome IV appeared to be an *accD* overexpressor when compared to the strong wild type plastome I, as judged from northern blot analyses. However, this overexpression, could not be detected in the plastome I variants that have a weakened competitive ability. Similar results were obtained for *ycf2*, which has an RNA of about 7 kb, reflecting the predicted size of the full-length transcript (SI Appendix, Fig. S14 and SI Text). Interestingly, lower bands, probably reflecting transcript processing and/or degradation intermediates, differ between the strong wild-type plastome I and the weak wild type plastome IV, with the latter being similar to the weak plastome I variants.

Since these analyses did not allow conclusions about the functionality of AccD or Ycf2 in our lines, we decided to determine the ACCase activity in chloroplasts isolated from a constant nuclear background. As shown in Fig. 4A, the presence of large mutations/polymorphisms in the N-terminus of the *accD* reading frame co-occurs with higher levels of ACCase enzymatic activity. Surprisingly, mutations/polymorphisms in *ycf2* also have an influence on ACCase activity, as revealed by lines that are not affected by mutations in *accD*. The molecular nature of this functional connection between *Ycf2* and ACCase activity is currently unclear, although *Ycf2* shares weak homologies to the FtsH protease (35, 36) which has a regulatory role in lipopolysaccharide synthesis in *Escherichia coli* (37). In any case, a simple relationship of ACCase activity and competitive ability of plastids is not present, but alterations in the earliest step of fatty acid biosynthesis can conceivably result in various changes in lipid metabolism (see also SI Appendix, SI Text).

**Fig. 4.**
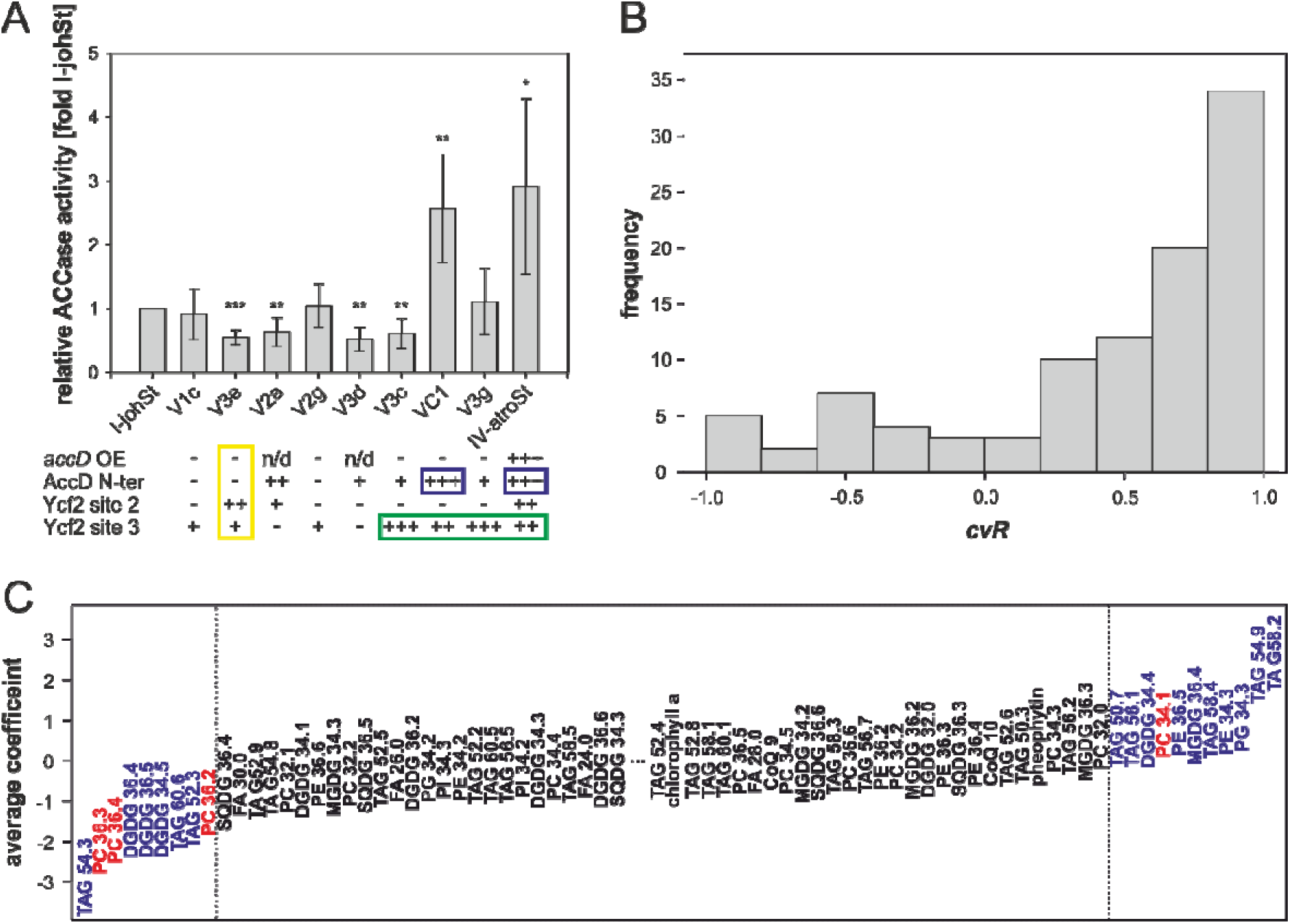
ACCase activity and prediction of inheritance strengths based on lipid level. (*A*) ACCase activity of chloroplasts with difference inheritance strengths, sorted according to competitive abilities (percentage biparental inheritance I-chi; cf. Fig. 2); *accD* OE = *accD* overexpressor; AccD N-ter = AccD N-terminus; Ycf2 site 2 and Ycf2 site 3; see SI Appendix, SI Text. Compared to I-johSt: -= not affected, + = mildly affected, ++ = intermediately affected, and +++ = strongly affected, n/d = not determined (cf. SI Appendix, Figs. S1-S5, and S14). Note the influence of mutations in *ycf2* on *accD* activity in a non-mutated *accD* background (yellow box), the striking co-occurrence of mutations in the *accD* N-terminus with ACCase activity (blue boxes), and co-occurrence of large mutations/polymorphisms in site 3 of Ycf2 with inheritance strengths (green box). Significance of difference compared to I-johSt was calculated using paired two-tailed t-test (*** *p* < 0.0005, ** *p* < 0.005, * p < 0.05). (*B*) Histogram of Pearson correlation coefficient cvR between actual and predicted inheritance strengths obtained from 100 cross-validation runs. Note shift towards 1.0, pointing to the predictive power of lipid data in the LASSO regression model. (*C*) Average linear LASSO model coefficients of the 102 lipids/molecules available for analysis (cf. SI Appendix, Table S4). The 20 identified predictive lipids are marked in color. PCs, the dominant phospholipids of the chloroplast outer envelope, are indicated in red. In general, large absolute values show predictive power either negatively or positively correlating with inheritance strength. Predictive lipids are designated when their absolute average weight was larger than one standard deviation of all weigh values obtained for the 102 lipids. For better presentability lipids/molecules with an absolute average weight ~0.00 were removed from the figure.

To examine the alterations in lipid biosynthesis, we determined the lipid composition of seedlings harboring strong and weak wild type chloroplasts as well as the variants that differ in chloroplast inheritance strength (SI Appendix, Table S3 and SI Text). Then, we employed a LASSO regression model to predict competitive ability of a given chloroplast (Fig. 4B). Since chloroplast inheritance strength is independent of photosynthetic competence (SI Appendix, SI Text), we included pale lines. The aim was to enrich the lipid signal responsible for inheritance strength, i.e. to deplete for structural lipids of the photosynthetic thylakoid membrane, which is a major source of chloroplast lipids (38). Indeed, out of 102 lipids analyzed, 20 predictive ones for inheritance strengths were identified (Fig. 4C; SI Appendix, Table S4 and SI Text). Strikingly, the signal is independent of greening (i.e. an intact thylakoid membrane system and the photosynthetic capacity), which is in line with the genetic data (SI Appendix, SI Text). This result hints at the chloroplast envelope determining assertiveness rates, a view that is supported by the fact that half of the predictive lipids come from lipid classes present in plastidial membranes and abundant in the chloroplast envelope such as MGDG, DGDG, PG, and PC (39). The remaining predictive lipids mostly represent storage lipids (TAG). This might be a result of an altered fatty acid pool (SI Appendix, SI Text). Statistical significance of enrichment of a given class could not be established due to low numbers (SI Appendix, Table S4 and S5, and SI Text), although, especially in the lipid class PC, which is the dominant phospholipid class in the chloroplast outer envelope (40), 4 out of 13 detected lipids were found to be predictive. In the chloroplast, PCs are specific to the envelope membrane and essentially absent from thylakoids. This makes it very likely that the lipid composition of the envelope membrane affects chloroplast competition. A possible explanation could be that strong and weak chloroplasts differ in division rates, for example, due to differential effectiveness in recruiting chloroplast division rings, which are anchored by membrane proteins (41). Alternatively, chloroplast stability might depend on envelope membrane composition. Strictly speaking, we cannot exclude a (further) involvement of extraplastidial membranes to the inheritance phenotype (most of the total cellular PCs are found in the ER and the plasma membrane) (38), but this would require a much more complicated mechanistic model.

## Discussion

The present work explains plastid competition following biparental inheritance from a mechanistic perspective and points to genetic loci that appear to be responsible for these differences. Moreover, since chloroplast competition can result in uniparental inheritance through the elimination of weak chloroplasts (9, 20, 30), at least for the evening primrose, the mechanistic explanation can be extended to uniparental transmission. Since about 20% of all angiosperms contain ptDNA in the sperm cell, it is likely that this mechanism is present in other systems (3, 5, 42). However, it should be emphasized that uniparental inheritance can be achieved by multiple mechanisms (2) and nuclear loci controlling the mode of organelle inheritance still need to be identified.

Arguably, the most surprising finding from our work is the discovery that chloroplast competition in evening primroses is essentially a metabolic phenotype and not directly connected to ptDNA replication or copy number (43). The underlying molecular loci are rapidly evolving genes throughout the plant kingdom. In general, the Ycf1 and Ycf2 proteins as well as the N-terminus of *accD* are highly variable in angiosperms (44-46). Interactions between them were repeatedly suggested (46, 47); e.g. the loss of *ycf1*, *ycf2* and *accD* genes from the plastome of grasses (48), as well as their common retention in many plastomes of non-photosynthetic parasites (49) provides room for speculation about functional interaction (46). In addition, *Silene*, a genus in which biparental chloroplast transmission is described (50), exhibits accelerated evolution of *accD* (51). In contrast, *Campanulastrum*, again displaying biparental chloroplast inheritance (9) has lost *accD* from the chloroplast genome, and its *ycf2* is under accelerated evolution (52). In pea, which usually shows uniparental chloroplast inheritance, one strain with biparental chloroplast inheritance exists and biparental transmission is accompanied by chloroplast/nuclear genome incompatibility in the resulting hybrid (53), with repeats in *accD* implicated as being responsible (54). Analysis of a very similar chloroplast/nuclear genome incompatibility in Oenothera also identifies repeats in *accD* as causative (55). Hence, correlation of the highly divergent *accD* gene (and/or *ycf2*) with the presence or absence of biparental inheritance and/or chloroplast incompatibility is certainly worth investigating on a broader phylogenetic scale. Connected with that, the *accD* plastome variants represent a superb material to study ACCase regulation and those of its nuclear counterparts. Interaction effects in different nuclear genomic background should be observable.

In the plastome of the evening primrose, the loci involved in chloroplast competition are very sensitive to replication slippage due to the presence of highly repetitive sequences, and that process appears to be the major mechanism of spontaneous chloroplast mutation (31, 44) (also see SI Appendix, SI Text). This result is somewhat reminiscent of recent findings in *Drosophila* where sequence variation in the non-coding regulatory region of the mitochondrial genome, containing the origins of replication, was associated with different competitive abilities. Similar to *Oenothera*, these sequences are highly repetitive, hypervariable and contribute to cytoplasmic drive. In *Drosophila*, they are among the most divergent ones in Metazoa pointing to their positive selection (19) (also see SI Appendix, SI Text).

Our analyses show that, due to their high mutation rates, cytoplasmic drive loci can evolve and become fixed in a population very quickly: In *Oenothera* the current view on the evolutionary history of the plastome is IV -> II -> I/III (56), with an estimated divergence time of approximately 830,000 years (44). This is largely based on chloroplast/nuclear incompatibility and can explain why plastomes IV and II (although compatible with the nuclear genomes of many *Oenothera* species), were outcompeted by the newly evolved aggressive plastomes I and III in many species. Interestingly, the extant plastomes I and III do not form a phylogenetic clade. The recently evolved strong plastome I clusters with the intermediately strong plastome II, with recently evolved plastome III as outgroup (SI Appendix, Fig. S1B). The divergence time of II and III, however, is only about 420,000 years (44). Hence, evolution and fixation of an aggressive cytoplasm happened twice independently within a very short time frame.

## Materials and Methods

### Plant material

Throughout this work, the terms *Oenothera* or evening primrose refer to subsection *Oenothera* (genus *Oenothera* section *Oenothera*) (56). Plant material used here is derived from the *Oenothera* germplasm collection harbored at the Max Planck Institute of Molecular Plant Physiology, Potsdam-Golm, Germany (57). Part of this collection is the so-called Renner Assortment, a collection of lines thoroughly characterized by the genetic school of Otto Renner (22, 58). Therefore, the original material of Franz Schötz in which he determined the classes of chloroplast replication speeds (21, 26) was available for correlation mapping (see below). For all other genetic or physiological work presented here, the nuclear genetic background of O. *elata* ssp. *hookeri* strain johansen Standard (59) was used. The employed chloroplast (genomes) are either native or were introgressed into that race by Wilfried Stubbe or Stephan Greiner. The wild type chloroplast genomes (I-johSt, I-hookdV, II-suavG, and IV-atroSt) are compatible with and hence green when combined with the johansen Standard nucleus. The chloroplast genome III-lamS confers a reversible bleaching, so-called *virescent*, phenotype in this genetic background (56, 60) (SI Appendix, Fig. S15). The white chloroplast mutants I-chi and IV-delta (SI Appendix, Fig. S15) are part of the huge collection of spontaneous plastome mutants compiled by Wilfried Stubbe and co-workers (31, 61, 62). Both mutants harbor a similar single locus mutation in the psaA gene (encoding a core subunit of photosystem I) and derive from the strong and weak wild type plastomes I-hookdV and IV-atroSt, respectively (31). Tables S6-S11 in the SI Appendix provide a summary of all strains and origins of the chloroplast genomes, including the *plastome mutator* induced variants that are subsequently described.

### Plastome mutator mutagenesis

The *plastome mutator* line is a descendant of the original isolate E-15-7 of Melvin D. Epp. The nuclear *pm* allele was identified after an ethyl methanesulfonate mutagenesis (63) in johansen Standard. When homozygous the *plastome mutator* causes a 200-1000x higher rate of chloroplast mutants compared to the spontaneous frequency. The underlying mutations mostly represent indels resulting from replication slippage events (32, 63-66).

Johansen Standard plants newly restored to homozygosity for the nuclear *plastome mutator* allele (*pm*/*pm*) were employed to mutagenize the chloroplast genome (I-johSt) as described (35). Homozygous *pm* plants were identified when new mutant chlorotic sectors were observed on them. On those plants, flowers on green shoots were backcrossed to the wild type *PM* allele as pollen donor. In the resulting *pm*/*PM* populations the chloroplast mutations were stabilized. This led (after repeated backcrosses with the *PM* allele and selection with appropriate markers against the paternal chloroplast) to homoplasmic green variants derived from the strong plastome I-johSt. The variants differ by certain indels or combination of indels, and the material was designated V1a, V1b, V2a, etc., where “V” stands for variant, the Arabic number for the number of the backcrossed plant in the experiment, and the small Latin letter for the shoot of a given plant. An additional line, named VC1, was derived from a similar *plastome mutator* mutagenesis of I-johSt, but the mutagenesis was conducted over several generations. Due to this fact, VC1, which is also a green variant, carries a much larger number of background mutations than do variants V1a, V1b, V2a, etc. (SI Appendix, Table S10). The two variant chloroplast genomes III-V1 and III-V2 (SI Appendix, Table S11) have a derivation similar to VC1. They are derived from the strong wild type chloroplast genome III-lamS, which displays a reversible bleaching (*virescent* phenotype) in the johansen Standard nuclear genetic background. To mutagenize this chloroplast genome, it was introgressed into the *pm*/*pm* background of johansen Standard by Wilfried Stubbe and self pollinated for a number of generations. When stabilized with the *PM* allele, it still displayed a *virescent* phenotype that is comparable to the original wild type plastome III-lamS (SI Appendix, Fig. S15). Both lines quite likely go back to the same ancestral *pm*/*pm*-johansen Standard III-lamS plant, i.e. experienced a common mutagenesis before separating them, although the number of (independent) mutagenizing generations is unclear.

### Determination of plastid inheritance strength

In the evening primroses biparental transmission of plastids shows maternal dominance, i.e. F1 plants are either homoplasmic for the maternal chloroplast or heteroplasmic for the paternal and maternal chloroplasts. If in such crosses one of the chloroplasts is marked by a mutation, resulting in a white phenotype, the proportion of variegated (green/white; i.e. heteroplasmic) seedlings can be used to determine chloroplast inheritance strength (as percentage of biparental inheritance). Moreover, if in such crosses one of the crossing partners is kept constant, the inheritance strength of all tested chloroplasts in respect to the constant one can be determined (20, 21, 26, 30). For example, in the I-chi crosses (where the strong white plastid is donated by the father; see below), more variegated seedlings are found in the F1, indicating that more paternal (white) chloroplasts were able to out-compete the dominating maternal green chloroplasts. Hence, in this crossing direction, small biparental percentage values indicate strong (assertive) plastomes from the maternal parent and high biparental values indicate weak variants were contributed by the maternal parent. The situation is reversed in the reciprocal cross where the white chloroplast is donated by the mother, as is the case in the IV-delta crosses. Here, the weak white chloroplast is maternal and strong green variants contributed by the pollen give high fractions of variegated plants in the F1, whereas low percentages of biparental progeny result when weak green variants are carried by the pollen donor.

### Crossing studies

All crossing studies between chloroplast genomes were performed in the constant nuclear background of the highly homozygous johansen Standard strain (see above). Germination efficiency in all populations was 100% (see SI Materials and Methods). Transmission efficiencies of the green plastome I variants (V1a, V1b, V2a, etc.) were determined using the white chloroplast I-chi (strong inheritance strength) and IV-delta (weak inheritance strength) as crossing partners, respectively. This allows the determination of the inheritance strength of a given green chloroplast relative to a white one based on quantification of the biparental (variegated) progeny among the F1, since progeny that inherit only chloroplasts from the paternal parent are extremely rare. The fraction of variegated (green/white) seedlings was assessed in the I-chi crosses where the white mutant was contributed by the paternal parent. Similarly, variegated (white/green) seedlings were quantified in the IV-delta crosses where the white mutant was donated by the maternal parent (21, 26, 30) (see above). In the I-chi crosses, the green plastome I variants, as well as the wild type chloroplast genomes I-johSt (strong inheritance strength; native in the genetic background of johansen Standard and the original wild type chloroplast genome used for mutagenesis), II-suavG (intermediate inheritance strength) and IV-atroSt (weak inheritance strength) were crossed as female parent to I-chi in three following seasons (2013, 2014, and 2015). In the IV-delta crosses, green variants and I-johSt were crossed as male parent to IV-delta, again in three independent seasons, 2013, 2014 and 2015. From each cross of each season randomized populations of 100-300 plants were grown twice independently, followed by visual assessment of the number of variegated seedlings/plantlets 14-21 days after sowing (DAS; I-chi crosses) or 7-14 DAS (IV-delta crosses). Based on these counts, the percentage of variegated progeny was calculated for each individual cross. To determine statistically significant differences between the transmission efficiencies of the plastome I variants and I-johSt, the numbers from all three seasons were summed for a particular cross and a Fisher’s exact test was employed.

A very similar experiment was performed to determine the inheritance strength of the two variants III-V1 and III-V2 (SI Appendix, SI Text), which derive from the strong chloroplast genomes III-lamS. Here in two independent seasons (2015 and 2016) the wild type I-johSt was used as pollen donor to induce variegation between the maternal plastome III (giving rise to a *virescent* phenotype) and the green plastome I (native in the background of johansen Standard; see above).

To determine transmission efficiencies independent of white chloroplast mutants or other bleached material, the plastome I variants (including their wild type I-johSt) were crossed to the green wild type plastomes IV-atroSt (weak inheritance strength) as female parent and to I-hookdV (strong inheritance strength) as male one in two independent seasons (2013 and 2014). F1 progeny was harvested 6 DAS by pooling 60-80 randomized seedlings and the ratios of the plastome types in the pool were analysed via MassARRAY^®^ (Agena Bioscience, Hamburg, Germany) as described below.

### MassARRAY^®^: multiplexed genotyping analysis using iPlex Gold

SNP genotyping to distinguish plastome I-johSt and I-hookdV/I-chi or I-johSt and IV-atroSt/IV-delta and subsequent quantification of their plastome ratios in appropriate F1s was carried out with the MassARRAY^®^ system (Agena Bioscience, Hamburg, Germany). The system was used to analyze chloroplast transmission efficiencies in different crosses. For this, total DNA was prepared from 60-80 randomized pooled plantlets 6 DAS. Then, 10 SNPs distinguishing the plastomes I-johSt and I-hookdV/I-chi (I/I assay) and 15 SNPs between I-johSt and IV-atroSt/IV-delta (I/IV assay) were selected. Two appropriate primers flanking the SNP and one unextended primer (UEP; binding an adjacent sequence to the SNP) were designed using MassARRAY^®^ Assay Design v4.0 (Agena Bioscience, Hamburg, Germany). Primer sequences, SNPs and their positions in I-johSt are listed in Table S12. Plastome regions were amplified in a 5 µl PCR reaction containing PCR buffer (2 mM MgCl_2_, 500 µM dNTP mix, 1 U HotStartTaq; Agena Bioscience, Hamburg, Germany), 10 ng DNA and 10 (I/I assay) or 15 (I/IV assay) PCR primer pairs, respectively, at concentrations ranging from 0.5-2.0 µM. The reaction mix was incubated for 2 min at 95°C in 96 well plates, followed by 45 cycles of 30 sec at 95°C, 30 sec at 56°C and 60 sec at 72°C, and a final elongation for 5 min at 72°C. Excess nucleotides were removed by adding 0.5 U Shrimp alkaline phosphatase (SAP enzyme) and SAP buffer (Agena Bioscience, Hamburg, Germany), followed by an incubation for 40 min at 37°C and 5 min at 85°C. For the primer extension reaction the iPLEX reaction mixture (containing Buffer Plus, Thermo Sequenase and termination mix 96; Agena Bioscience, Hamburg, Germany) and, depending on the primer, the extension primers at a concentration of 7-28 µM were added. Sequence-specific hybridization and sequence-dependent termination were carried out for 30 sec at 94°C, followed by 40 cycles of 5 sec at 94°C plus five internal cycles of 5 sec at 52°C and 5 sec at 80°C, and finally 3 min at 72°C. After desalting with CLEAN resin (Agena Bioscience, Hamburg, Germany) the samples were spotted on 96-pad silicon chips preloaded with proprietary matrix (SpectroCHIP; Agena Bioscience, Hamburg, Germany) by using the Nanodispenser RS1000 (Agena Bioscience, Hamburg, Germany). Subsequently, data were acquired with MALDI-TOF mass spectrometer MassARRAY^®^ Analyzer 4 (Agena Bioscience, Hamburg, Germany) and analyzed with the supplied software. To identify significant differences in the frequencies of paternal ptDNA Kruskal-Wallis one-way analysis of variance (ANOVA) on ranks was performed.

### *k*-means clustering to classify inheritance strength

For the wild type chloroplasts, inheritance strength was classified using the biparental transmission frequencies (percentage of variegated plants in F1, see above) of the chloroplasts “biennis white” and “blandina white” according to Schötz (26). For details see SI Appendix, SI Text. Both crossing series included the same 25 wild type chloroplasts, 14 of which had fully sequenced genomes and were employed for correlation mapping (SI Appendix, Table S7 and below). For original data, see Schötz (1968) (26), summaries in Cleland (1972, p. 180) (22), or SI Appendix, Table S13. Based on the two transmission frequencies, the wild type plastomes were clustered using the *k*-means algorithm with Euclidean distance as distance dimension. The optimal number of centers was calculated with the pamk function of the fpc package, as implemented in R v.3.2.1 (67). Strikingly, essentially the same three classes (strong, intermediate, and weak) were obtained that had been previously determined by Schötz (20, 22) (SI Appendix, Fig. S16 and SI Text).

For the variants, we used the transmission frequencies from I-chi and IV-delta crosses obtained from this work (Fig. 2; SI Appendix, Table S10 and SI Text). Since the data-driven determination of the optimal number of clusters (k = 2; see above) does not reflect the biological situation, upon repeated *k*-means runs, we chose the number of centers with the best trade-off between lowest swapping rate of the samples between the clusters and the biological interpretability. This approach resulted in four classes (see SI Text and Fig. S16 for details).

### Correlation mapping

For correlation mapping in the wild types, 14 completely sequenced plastomes (GenBank accession numbers EU262890.2, EU262891.2, KT881170.1, KT881171.1, KT881172.1, KT881176.1, KU521375.1, KX687910.1, KX687913.1, KX687914.1, KX687915.1, KX687916.1, KX687917.1, KX687918.1) with known inheritance strength (20, 22, 26) (SI Appendix, Table S7) were employed. The eight chloroplast genomes assigned to accession numbers starting with KU or KX were newly determined in the course of this work. Mapping of genetic determinants in the green variants was done in 18 fully sequenced mutagenized plastomes (V1a, V1b, V1c, V2a, V2b, V2g, V3a, V3b, V3c, V3d, V3e, V3f, V3g, V3h, V4b, V4c, V7a, and VC1), as well as their wild type reference (I-johSt; GenBank accession number AJ271079.4).

In both sequence sets, divergence at a given alignment window was correlated to the experimentally determined inheritance strengths of a chloroplast genome. For the wild types, inheritance strength was measured using the paternal transmission frequencies (percentage of variegated plants in the F1) of the chloroplasts “biennis white” or “blandina white” according to Schötz (26), see SI Text for details, or *k*-means classes combining the two datasets by clustering (SI Appendix, Table S13, SI Text and see above). For the variants, we used the transmission frequencies from the I-chi and IV-delta crosses determined in this work, as well as the *k*-means classes that were obtained from them (Fig. 2; SI Appendix, Table S10, SI Text and above).

For correlation of these transmission frequencies to loci on the chloroplast genome, the redundant inverted repeat A (IR_A_) was removed from all sequences. Then, plastomes were aligned with ClustalW (68) and the alignments were curated manually (SI Appendix, Dataset S2). Subsequently, using a script in R v3.2.1 (67) (SI Appendix, Dataset S3), nucleotide changes (SNPs, insertion and deletions) relative to a chosen reference sequence plastome [I-hookdV (KT881170.1) for the wild type set and I-johSt (AJ271079.4) for the variant set; see SI Text for details] were counted window-wise by two approaches: (i) segmenting the reference sequence in overlapping windows using a sliding window approach with a window size of 1 kb and a step size of 10 bp, yielding a matrix of 13,912 × 13 (wild type set) and 13,668 × 18 (variants), respectively, or (ii) defining regions of interest with correspondingly chosen window sizes. Then, Pearson’s and Spearman’s correlation coefficients were calculated between (i) the total count of nucleotide changes for every plastome in the aligned sequence window compared to the reference (total sequence divergence) and (ii) the determined inheritance strength of the plastomes (SI Appendix, Fig. S17 and Dataset S1). For the sliding window approach, *p*-values were adjusted for multiple testing using Benjamini-Hochberg correction. To reduce the number of *p*-value adjustments, adjacent alignment windows with identical count vectors were collapsed into one. To annotate the correlation mapping script output file, gene annotation of the consensus sequence (SI Appendix, Dataset S2) was converted into bed format (SI Appendix, Dataset S3) and combined with the correlation bins using intersectBed and groupBy of the bedtools package (69). For visualization (Fig. 1A and 3A; SI Appendix, Fig. S1 and S10), correlation coefficients obtained for every alignment window were plotted as a function of the alignment position. Correlation values above the *p*-value threshold (> 0.05) are greyed out. Linear chloroplast genome maps were derived from the annotation of the consensus of both sequence sets (SI Appendix, Dataset S2) and drawn by OrganellarGenomeDRAW v1.2 (70) in a linear mode using a user-defined configuration XML file. Alignments of selected plastome regions were visualized in Geneious v10.2.3 (71) and subsequently, as the output of OrganellarGenomeDRAW, edited in CorelDraw X8 (Corel Corporation, Ottawa, ON, Canada).

### ACCase activity assay

ACCase activity was measured in isolated chloroplast suspensions (72, 73) diluted to 400 µg chlorophyll/ml (SI Appendix, SI Materials and Methods). To validate equilibration to chlorophyll, protein concentration using a Bradford assay (Quick StartTM Bradford 1x Dye Reagent; Bio-Rad, Hercules, CA, USA; with BSA solutions of known concentrations as standards) and chloroplast counts per ml suspension were determined for the same samples. For chloroplast counting, the suspension was further diluted 1:10, with 15 µl subsequently loaded on a CELLOMETER™ Disposable Cell Counting Chamber (Electron Microscopy Sciences, Hatfield, PA, USA) and analyzed under a Zeiss Axioskop 2 (Zeiss, Oberkochen, Germany). For each sample six “B squares” were counted and chloroplast concentration was calculated as chloroplasts/ml = 10 × average count per “B square” / 4 × 10. All three equilibration methods gave comparable results.

ACCase activity was measured as the acetyl-CoA-dependent fixation of H^14^CO_3_^-^ into acid-stable products. For each plant line (I-johSt, V1c, V3e, V2a, V2g, V3d, V3c, VC1, V3g and IV-atroSt) three independent chloroplast isolations (= three biological replicates) were analyzed in triplicates, including individual negative controls (minus acetyl-CoA) for each measurement. 10 µl of chloroplast suspensions were incubated with 40 µl of reagent solution having a final concentration of 100 mM Tricine-KOH pH 8.2, 100 mM potassium chloride, 2 mM magnesium chloride, 1 mM ATP, 0.1 mM Triton X-100, 10 mM sodium bicarbonate, 0.5 mM acetyl-CoA and 40 mM radioactively labelled sodium bicarbonate (NaH^14^CO_3_ ca. 4000 d*pm*/nmol; Amresham, Little Chalfont, UK) at room temperature for 20 min. For the negative control, acetyl-CoA in the reaction mixture was replaced by water. Reactions were stopped by adding 50 µl 2 M hydrochloric acid. The sample was transferred to a scintillation vial and acid labile radioactivity (i.e. remaining H^14^CO_3_^-^) was evaporated by heating for 20 min at 85°C. After addition of 3 ml scintillation cocktail (Rotiszint^®^ eco plus, Carl Roth, Karlsruhe, Germany), the acid stable radioactivity from incorporation of H^14^CO_3_^-^ (^14^C d*pm*) were detected by liquid scintillation counter (LS6500, Beckman Coulter, Brea, CA). ACCase activity is represented as the ^14^C incorporation rate into acid stable fraction (d*pm* min^-1^) calculated by dividing the total fixed radioactivity by 20 min. The rates in three replicated reactions were averaged and corresponding values from negative control samples were subtracted and normalized by the number of chloroplast to gain ACCase activity in individual samples. The average rates were calculated for each line. To combine all measurements, relative ACCase activities were calculated for each experiment as relative to the I-johSt line, and significant differences between each line and the wild type were identified using two-tailed paired t-test, followed by *p*-value adjustment using the Benjamini-Hochberg procedure.

### Predictability of inheritance strength based on lipid-level data as explanatory variables

Lipidomics data from *Oenothera* seedlings of the strain johansen Standard, harboring chloroplast genomes with different assertiveness rates (see SI Appendix, SI Materials and Methods), were analyzed jointly to test for predictability of inheritance strength based on lipid levels. For this, 33 probes representing 16 genotypes whose chloroplast genomes ranged from inheritance strength class 1 to 5 (see SI Appendix, SI Text) were measured in five replicates in three independent experimental series (Table S3; SI Text). In this dataset, a total of 184 different lipids/molecules could be annotated (SI Appendix, Dataset S4; see above). To normalize across experiments, the data from each series were log-transformed and median-centered based on genotypes with inheritance strengths = 1, i.e. for every lipid/molecule, its median level across all “inheritance strengths = 1 genotypes” was determined and subtracted from all genotypes tested in the respective experimental series. “Inheritance strength 1” was then selected to serve as a common reference across all three experimental series. Subsequently, the three experimental series were combined into a single set. Only those lipids/molecules were considered further, for which level-data were available across all three datasets, leaving 102 lipids/molecules for analysis (SI Appendix, Dataset S4)

#### LASSO regression model

Inheritance strength was predicted based on the median-centered lipid level data using LASSO, a regularized linear regression approach (74) as implemented in the “glmnet” R-software package (R v3.2.1) (67). Glmnet was invoked with parameter α set to 1 to perform LASSO regression (SI Appendix, Dataset S4). The penalty parameter λ was determined from the built-in cross-validation applied to training set data (i.e. all but two randomly selected genotypes) and set to the one-standard-error estimate deviation from the optimal (minimal error) value and assuming Gaussian response type. All other parameters were taken as their default values.

#### Predictive lipids

As a regularized regression method, LASSO aims to use few predictor variables, which allows better identification of truly predictive lipids. Summarized from all 100 performed cross-validation runs, lipids/molecules were ranked by their mean linear model coefficients assigned to them in the LASSO regression with their absolute value indicating influence strength, and their sign indicating positive or negative correlation of their abundance to inheritance strength.

#### Test for enrichment of predictive lipids/molecules in lipid classes

Across all 100 cross-validation runs, the importance of each of the 102 molecules was assessed based on their average absolute weight factor (avgW) by which they entered the 100 LASSO models. Molecules with avaW of greater than one standard deviation obtained across all 102 molecules were considered important. Then, all lipid/molecule were assigned to their respective class (MGDG, DGDG, SQDG, PG, PC, PI, PE, FA, PE, TAG, CoQ, chlorophyll, and pheophytin) and every class was tested for enrichment in the lipid/molecule set considered to be important. This was done by employing Fisher’s exact test yielding both *p*-values and odds-ratios. The *p*-values express enrichment the odds-ratio express the relative enrichment or depletion of a particular class among the set of important lipids.

## Acknowledgments

We thank Werner Dietrich and Wilfried Stubbe who compiled the comprehensive *Oenothera* collection that allowed us to use the original material of the Renner school. John B. Ohlrogge is acknowledged for advice with the ACCase activity assay, Michael Tillich for help with the MassArray^®^ system and Stephen I. Wright for fruitful discussion on PGLS. We further thank the MPI-MP GreenTeam for their support, the Ornamental Plant Germplasm Centre for providing the line chicaginensis de Vries and Liliya Yaneva-Roder for technical assistance. This research was supported by the Max Planck Society to S.G., P.G., D.W., M.A.S., and R.B. DAPI fluorescent patterns were analysed with microscopy equi*pm*ent funded by the Polish National Science Centre (NCN) to H.G. (2015/19/B/N22/01692).

**Author Contribution**
J.S. performed the main experimental work. P.G., A.F., J.K., D.W., M.A.S., H.G., T.O., T.P., B.B.S, and S.G. provided supportive data. A.F. and S.G. developed the correlation mapping approach, J.K. implemented PGLS. All authors analyzed and discussed the data. B.B.S. and S.G. designed the study. S.G. and J.S. wrote the manuscript. P.G., A.F., D.W., M.A.S., H.G., R.B. and B.B.S. participated in writing.

