## Supplementary material for "Chloroplast competition is controlled by lipid biosynthesis in evening primroses": SI Appendix

**This PDF file includes:**

Supplementary Materials and Methods
Supplementary Information Text
Figs. S1 to S19
Tables S1 to S17
References for SI reference citations

**Other supplementary materials for this manuscript include the following:**

Datasets S1 to S4

### Datasets S1 to S4 for this manuscript include the following files:

***Dataset S1***

Correlation mapping full variant plastomes.xlsx
Correlation mapping full WT plastomes.xlsx
Correlation mapping selected loci.xlsx

These files comprise all correlation mapping data of this work including PGLS analyses, if applicable.

***Dataset S2***

WT_set_alignment.fas
I_variant_set_alignment.fas
III_variant_set_alignment.fas

These files contain the alignments used for correlation mapping, PGLS analysis and detection of polymorphisms in the variant plastomes in the FASTA format.

WT_consensus_annotation.gb
I_variants_consensus_annotation.gb
III_variants_consensus_annotation.gb

These files contain the annotated consensi from the alignments from above in the GenBank format, necessary to localize polymorphism in the alignments in coding or non-coding regions.

III-V1_annotation.gb
III-V2_annotation.gb

Annotation files in the GenBank format of the very weak variants of plastome III.

***Dataset S3***

correlation_mapping_script_and_data.zip

Zip-archive to reproduce the correlation mapping analyses present in this work. It contains the R-script, all necessary input files, as well as a README file.

lipids_analysis_script_and_data.tar.gz

R-script and lipid input data of the LASSO analysis. Inheritance strength classes of the samples is provided in Table S3.

***Dataset S4***

Lipids Experiment 1.xlsx
Lipids Experiment 2.xlsx
Lipids Experiment 3.xlsx

Lipid data as measured in experiments 1-3 of this work.

### Supplementary Materials and Methods

#### *Germination, plant cultivation and cross pollination*

Fresh seeds were germinated on wet filter paper at 27°C at 100-150 µE $m^{-2}$ $s^{-1}$. With this method essentially 100% germination is achieved after 1-3 days. If desired, seedlings were then cultivated to the appropriate developmental stage in a glasshouse. Crosses with flowering plants were performed as published earlier (1, 2).

#### *DNA isolation*

Total DNA for Illumina sequencing was isolated described before (2). For PCR analyses, quantitative real-time PCR and MassARRAY® experiments total DNA was isolated with the DNeasy Plant Mini Kit (Qiagen, Hilden, Germany) (3).

#### *DNA sequencing*

Illumina sequencing was performed at the Max Planck-Genome-Centre in Cologne (Germany), as described previously (2, 4). Exclusively "PCR-free" paired-end libraries (375 bp insert size) generated from total DNA isolations were employed. 100 bp Illumina paired-end reads were generated to determine the sequences of the plastome I and III variants, whereas 150 bp Illumina paired-end reads were used to assemble the wild type plastomes. Sanger sequence data were obtained from Eurofins MWG Operon (Ebersberg, Germany).

#### *Plastome assembly, sequence annotation and repeat analysis*

Complete chloroplast genomes were assembled with SeqMan NGen v12.1.0 or v13.0.0 (DNASTAR, Madison, WI, USA). Wild type and plastome III variants were done *de novo*. The sequences of the green plastome I variants derive from reference-guided assemblies. For this, their original wild type chloroplast genome I-johSt of *O. elata* ssp. *hookeri* strain johansen Standard (AJ271079.4) was used as reference (2). The notoriously repetitive regions in the *Oenothera* plastome, where individual repetitive elements can span more than an Illumina read length and/or the insert size of the employed paired-end libraries, namely upstream and/or within *accD*, *ycf1* (*tic214*), *ycf2*, and the *rrn16* - *trnI-GAU* spacer (*oriB*) were determined and/or confirmed by Sanger sequencing in all chloroplast genomes discussed in this work. Finalized sequences were annotated by GeSeq v0.9 (5) and the wild type chloroplast genomes were prepared for NCBI submission using GB2sequin v1.3 (6). Repeat structures were analyzed based on an inspection by eye and/or employing the EMBOSS suite (7) as previously described (8).

***Controlling for phylogenetic relatedness in the correlation mapping approach of the wild type plastomes***

Using the 13,912 1 kb windows of the wild type correlation mapping analyses (Materials and Methods), we determined whether the total count of nucleotide changes for every plastome in the aligned sequence window compared to the reference (total sequence divergence) significantly explained the inheritance strength of the plastomes, while controlling for shared patterns of nucleotide divergence between related lineages. To do this, we ran a phylogenetic generalized linear squares (PGLS) analysis using the *nlme* package in R v3.2.1 (9) in each genomic window. As underlying phylogeny, we used the maximum likelihood tree estimated from our manually curated plastome-wide ClustalW (10) multiple alignment (Dataset S2 and Materials and Methods; Fig. S1), obtained by employing Mega v7.02.6 (11) with its standard parameters and 300 bootstrap replications. Because our tree only encompasses 14 distinct lineages, we used the simplest model of phylogenetic correlation structure (12) to prevent overfitting by additional parameter estimation. In order to conform with the PGLS model assumptions, we repeated this analysis for our two continuous response variables - % of biparental inheritance from our biennis and blandina crosses (Table S7). After extracting *p*-values from the slope of each window-wise phylogenetically informed correlation, we controlled for an inflated false discovery rate by applying a Benjamini and Hochberg correction. Uncorrected *p*-values were plotted as a function of alignment position and *p*-values below the significance threshold of 0.05 were greyed out. Alignment windows whose corrected *p*-values still remained significant after correction were marked in red in the original Person/Spearman correlation mapping plot (Fig. S1).

***Chloroplast isolation***

For isolation of chloroplasts, *Oenothera* leaf tissue was harvested 7-8 weeks after sowing and processed as described previously (2). However, minor modifications were applied to allow a rapid isolation of chloroplasts from six plant lines in parallel: 35 g of leaf material was homogenized in 500 ml BoutHomX buffer. The pellet from the first centrifugation step was re-suspended in 100 ml ChloroWash and, after one filtration, the volume was adjusted to 150 ml before the second filtration. After the second centrifugation step re-suspended chloroplasts were loaded on two Percoll step gradients (each: 7 ml 85% Percoll, 14 ml 45% Percoll). Subsequent to gradient centrifugation the recovered chloroplasts were washed with 30 ml ChloroWash, followed by three additional washing steps with smaller volumes and a final re-suspension of the chloroplasts in 300-500 µl ChloroWash for the following ACCase activity measurements.

***Lipid extraction, mass spectrometry sample preparation and measurements***

Metabolites were extracted according to published protocols (13) from 50 mg *Oenothera* seedlings harvested 6 DAS. In brief, frozen tissue was homogenized by a ball mixer mill and transferred to cooled 2.0

ml round bottom microcentrifuge tubes. Subsequently, each sample was resuspended in 1.0 ml of a -20°C methanol:methyl-*tert* butyl-ether [1:3 (v/v)] mixture, containing 0.5 µg of 1,2-diheptadecanoyl-*sn*-glycero-3-phosphocholine (Avanti Polar Lipids, Alabaster, AL, USA) as an internal standard. Samples were then immediately vortexed before incubation for 10 min at 4°C on an orbital shaker. This step was followed by ultra-sonication in an ice-cooled bath-type sonicator for an additional 10 min. To separate the organic from the aqueous phase, 650 µl of a $H_2O$:methanol mix [3:1 (v/v)] was added to the homogenate, which was briefly vortexed before being centrifuged for 5 min at 14,000 *g*. Finally, 500 µl of the upper methyl-*tert* butyl-ether phase, containing the hydrophobic (lipid) compounds, was placed in a fresh 1.5 ml microcentifuge tube. This aliquot was either stored at -20°C for up to several weeks or immediately concentrated to complete dryness in a speed vacuum concentrator at room temperature. Prior to analysis the dried pellets were re-suspended in 400 µL acetonitrile:isopropanol [7:3 (v:v)], ultra-sonicated and centrifuged for 5 min at 14,000 *g*. The cleared supernatant was transferred to fresh glass vials and 2 µl of each sample was injected onto a $C_8$ reverse phase column (100 mm x 2.1 mm x 1.7 µm particles) using a Acquity UPLC system (Waters, Manchester, UK). In addition to the individual samples, we prepared pooled samples, in which 10 µl aliquots of each sample from the whole sample collection were combined. These pooled samples were measured after every 20$^{th}$ sample, to provide information on system performance including sensitivity, retention time consistency, sample reproducibility and compound stability.

The mobile phase for our chromatographic separation consisted of Buffer A (1% 1 M $NH_4$-acetate and 0.1% acetic acid in UPLC MS grade water), while Buffer B contained 1% 1 M $NH_4$-acetate and 0.1% acetic acid in acetonitrile/isopropanol [7:3 (v:v)] (BioSolve, Valkenswaard, Netherlands). The flow rate of the UPLC system was set to 400 µl/min with a Buffer A/Buffer B gradient of 1 min isocratic flow at 45% Buffer A (55% Buffer B), 3 min linear gradient from 45% to 25% Buffer A (55% to 75% Buffer B), 8 min linear gradient from 25% to 11% Buffer A (75% to 89% Buffer B), and 3 min linear gradient from 11% to 1% Buffer A (89% to 99% Buffer B). After cleaning the column for 4.5 min at 1% Buffer A/99% Buffer (B) the solution was set back to 45% Buffer A/55% Buffer (B) and the column was re-equilibrated for 4.5 min, resulting in a final run time of 24 min per sample.

Mass spectra were acquired with an Orbitrap-type mass spectrometer (Exactive; Thermo-Fisher, Bremen, Germany) and recorded in the full scan mode, covering a mass range from 100-1,500 m/z. The resolution was set to 60,000 with 2 scans per second, restricting the maximum loading time to 100 ms. Samples were injected using the heated electrospray ionization source (HESI) at a capillary voltage of 3.5 kV in positive and negative ionization mode. A sheath gas flow value of 40 was used, with an auxiliary gas flow value at 20 and a capillary temperature of 200°C, while drying gas temperature in the heated electro spray source was 350°C. The skimmer voltage was set to 20 V with tube lens value at 140 V. The spectra were recorded from 0 to 20 min of the UPLC gradients.

***Data processing and normalization of lipid data***

Data analysis of the raw files (*.raw) was performed using QI for metabolomics v2.3 (Nonlinear Dynamics, Newcastle upon Tyne, UK) according to the vendor description. Data were normalized to the internal standard (1,2-diheptadecanoyl-*sn*-glycero-3-phosphocholine) and the exact fresh weight of each sample. Lipid annotation was performed manually as described (14). Statistical data analysis was performed using Excel 2013 (Microsoft, Redmond, WA, USA), R v3.2.1 (15) and SIMCA-P v13.0 (Umetrics, Umea, Sweden).

***Determination of photosynthetic parameters***

Gas exchange measurements were performed with a GFS-3000 open gas exchange system equipped with the LED array unit 3055-FL as actinic light source for simultaneous chlorophyll a fluorescence measurements (Heinz Walz GmbH, Effeltrich, Germany). Light response curves of $CO_2$ assimilation were measured at 22°C cuvette temperature with 17,500 ppm humidity and a saturating $CO_2$ concentration of 2,000 ppm, to fully repress photorespiration. Plants were dark-adapted for a minimum of 30 min. Then, the maximum quantum efficiency of photosystem II in the dark-adapted state ($F_V/F_M$) and leaf respiration were determined. Afterwards, the actinic light intensity was first set to the growth light intensity of 200 µE $m^{-2}$ $s^{-1}$, followed by measurements at 500, 1,000, and finally 1,500 µE $m^{-2}$ $s^{-1}$. At each light intensity, gas exchange was recorded until a steady state of transpiration and leaf assimilation was reached. Maximum leaf assimilation was corrected for the respiration measured in darkness. After the end of the gas exchange measurements, the chlorophyll content and chlorophyll a/b ratio of the measured leaf section were determined in 80% (v/v) acetone according to (16). Leaf absorptance was calculated from leaf transmittance and reflectance spectra as 100% minus transmittance (%) minus reflectance (%). Spectra were measured between 400 and 700 nm wavelength using an integrating sphere attached to a photometer (V650, Jasco Inc., Groß-Umstadt, Germany). The spectral bandwidth was set to 1 nm, and the scanning speed was 200 nm $min^{-1}$.

***Fluorescence microscopy and differential interference contrast to analyze ptDNA nucleoids, chloroplast volume and number per cell***

We investigated leaf material from the central laminal region of the first true leaf 25 DAS. For this, four pieces of 5 $mm^2$ were excised from five individual plants per line and fixed. DAPI (4',6-diamidino-2-phenylindole) stains of ptDNA nucleoids and fluorescence microscopy were conducted as previously described (17, 18) with minor modifications: In brief, excised leaf fragments were fixed with 3% glutaraldehyde in 50 mM phosphate buffer (pH 7.2), washed in 1x PBS buffer (phosphate-buffered saline,

137 mM NaCl, 2.7 mM KCl, 10 mM $Na_2HPO_4$, 1.8 mM $KH_2PO_4$, pH 7.2) and macerated in 1% (w/v) cellulase and 1% (w/v) pectinase solution (both Sigma-Aldrich, St. Louis, MO, USA) in the 1x PBS for 30 min at 37°C. Subsequently, explants were washed in 1x PBS and stored at 4°C until use. For microscopy, small tissue sectors were gently squeezed into a drop of the 1x PBS between a microscope slide and a cover glass. Then preparations were frozen in liquid nitrogen and, after removing the cover glasses, air dried and mounted in a drop of DAPI solution [5 µg/ml DAPI (Sigma-Aldrich, St. Louis, MO, USA) in 1x PBS buffer in 70% glycerol ("for fluorescence microscopy"; Merck, Darmstadt, Germany)]. DAPI as fluorochrome is considered to be sensitive enough to detect DNA of a single plastid genome copy (19). The preparations were sealed with Fixogum rubber cement (Marabu, Tamm, Germany) and examined with a Nikon Eclipse Ni-U upright epifluorescence microscope equipped with a cooled monochrome camera (Nikon, Chiyoda, Japan) under a 100x UV objective. For each investigated cell, five to seven picture frames were digitally captured, each at a different focal plane. The frames were stacked and combined using standard macro commands of the Combine ZP software v1.0 developed by Alan Hadley (http://combinezp.software.informer.com). Nucleoids were counted manually in Adobe Photoshop CS3 (Adobe Systems San Jose, CA, USA). For this, chloroplasts in a cell were delimited and nucleoids identified (Fig. S13). Only chloroplasts with non-overlapping nucleoids were subjected to analysis. Overall we examined 22 cells per plant line and 6-29 chloroplasts per cell. In total 26,120 nucleoids were counted (Table S14). The significance of differences between all lines was tested by one-way ANOVA. In addition, to test differences between the strong plastome I-johSt and weak plastome V3g, VC1, and IV-atroSt, respectively, two-tailed homoscedastic *t*-test was calculated followed by *p*-value adjustment according to Benjamini-Hochberg.

For differential interference contrast (DIC) microscopy, explants excised as described above were transferred to 10% formalin in phosphate buffer (Tissue-Prep Buffered 10% Formalin; Electron Microscopy Sciences, Hatfield, PA, USA). Samples were then evaporated for at least 1 h and incubated at 4°C overnight. After washing with sterile water, leaf fragments were incubated under rotation in 0.1 M EDTA for 2 h at room temperature, followed by incubation at 4°C overnight. Directly before analysis, samples were incubated for 3 h at 60°C while shaking (500 rpm). Leaf pieces were mounted in water on a slide and cells were released by softly tapping on the top of the cover slide. Analysis was performed on a motorized epifluorescence microscope Olympus BX61 under a 40x objective (Olympus, Shinjuku, Japan) using DIC optics. This allows capturing of a focal plane in which all chloroplasts within a cell are visible. Subsequently, counting of chloroplasts per cell and measurement of chloroplast length and width was carried out with the Olympus cellSens Dimension software v1.7 (Olympus, Shinjuku, Japan). 4-5 individuals per plant line, and at least two leaf pieces per individual plant and a minimum of 5 spongy mesophyll cells per piece were investigated. To determine chloroplast numbers per cell, 55 cells per plant line were analyzed and overall

11,645 chloroplasts counted. For measuring chloroplast length and width 20 cells per plant line were chosen and three plastids per cell measured resulting in 480 individual measurements. Chloroplast volume index was calculated according length x width$^2$ (20). For statistical analysis of each experiment, for comparison of all lines, one-way ANOVA was performed. In addition, to test differences between the weak plastome I variants or IV-atroSt and the strong wild type I-johSt, a two-tailed homoscedastic *t*-test followed by adjustment of *p*-values according to Benjamini-Hochberg was performed. (Tables S15 and S16).

***Quantitative real-time PCR to determine plastid DNA amounts***

Plastid DNA content of wild type and variants with different inheritance strength were analyzed in a developmental series including 5, 21, and 32 DAS. For each line, three DNA preparations from independent pools of 20 individuals (5 DAS), or two DNA preparations from independent pools of three plantlets (21 DAS) or two DNA preparations from independent pools of three second leaves (32 DAS) were isolated, and subsequently analyzed two times in triplicate by qPCR: Reactions containing LightCycler® 480 SYBR Green I Master (Roche Diagnostics GmbH, Mannheim, Germany), 0.75 ng total DNA and 0.5 µM primers were incubated for 5 min at 95°C, followed by 40 cycles of 10 sec at 95°C, 10 sec at 58°C and 15 sec at 72°C in a LightCycler® 480 II instrument (Roche Diagnostics GmbH, Mannheim, Germany). Primer sequences, accession numbers of target genes/loci, size of amplification products, and primer efficiencies (that have been determined based on dilution series of total johansen Standard DNA harboring either plastome I-johSt or IV-atroSt) are provided in Table S17. ptDNA copy numbers were quantified for the plastid genes *rbcL*, *psbB* and *ndhI*, which are topographically well separated on the plastid genome, and normalized to three nuclear loci M02, M19, and *pgiC* (3, 21). The nuclear loci are only present once in the nuclear genome of johansen Standard, as judged from coverage analysis of Illumina libraries. Additionally, three markers for mitochondrial DNA (mtM03, mtM04, mtM06) (22) were included in the calculation. Data were analyzed with the LightCycler® 480 software v1.5.0 SP4 (Roche Diagnostics GmbH, Mannheim, Germany) employing the “Advanced Relative Quantification” method that incorporates primer efficiencies. Target/Reference values calculated by the software were used to determine the proportion of ptDNA per total DNA (including ptDNA, mtDNA and nuclear genome) by employing the approximate size of the nuclear (C1 about 1 GB) (4), mitochondrial (about 400 kb) (22, 23) and chloroplast (about 160 kb) (8) genomes, respectively. For illustration, arbitrary units were calculated by setting the I-johSt 5 DAS values of the plastid targets (*rbcL*, *psbB*, *ndhI*) to 1. Hence, the relative amount of ptDNA for all other genotypes and developmental stages is expressed as “fold I-johSt 5 DAS”: *((Target$^{pt}$/Ref)*plastome size [bp]/((Target/Ref)*plastome size [bp] + C1 nuclear genome size [bp] + (Target$^{mt}$/Ref)* chondriome size [bp]))/(value 5 DAS).* Significance of the differences between I-johSt and IV-atroSt/plastome I variants for each developmental stage was calculated with a one-sample *t*-test followed by multiple testing *p*-values correction according to Benjamini-Hochberg.

***Detection of RNA via RNA gel blot analyses***

Total RNA was isolated as described previously (2), with 2.5 µg separated in 1% (w/v) agarose gels under denaturing conditions and transferred to a nylon membrane (Hybond™-N; GE Healthcare, Chicago, IL, USA) by capillary action with 10x SSC (1x SSC: 0.015 M sodium citrate, 0.15 M NaCl). RNA was covalently linked to the membrane using a UV cross linker (Vilber Lourmat, Ebertharfzell, Germany). Transfer success and equal loading was visualized by methylene blue staining (0.03% methylene blue, 0.3 M Na-acetate). Membranes were incubated in Church buffer [0.5 M $NaH_2PO_4$, 7% SDS (w/v), 1 mM EDTA; pH 7.2] for 1 h. Hybridization was performed after addition of radiolabeled *accD* or *ycf2* probes overnight at 65°C. Double-stranded DNA probes were obtained by PCR amplification with gene specific primers AbaccDfor (5´-AGTATGGGATCCGTAGTCGG-3´) and accD_cons_rev (5´-ATTCAGCCGTTT GTGAACCCTC-3´) for *accD*, and the primers ycf2Vno6 (5´-TAATGATCGAGTGACATTGC-3´) and Ycf2_VP30rev (5´-CTCTTCGTCTTCCTCTTCAAGC-3´) for *ycf2*. Purification of PCR fragments was done with the NucleoSpin® Gel and PCR Clean-up Kit (Machery-Nagel GmbH & Co. KG, Düren, Germany) and random-prime $^{32}$P-labelling with the Megaprime DNA Labelling System (GE Healthcare, Chicago, IL, USA). Membranes were washed first with 1x SSC/0.2% SDS and two times with 0.5x SSC/0.2% SDS for 20 min each prior to exposition overnight. Radioactive signals were visualized with Typhoon TRIO+ ImageQuant (GE Healthcare, Chicago, IL, USA).

### Supplementary Information Text

**1. Generation of green variants with altered inheritance strength**

**2. The green plastome I variants do not display impaired growth, altered chloroplast morphology, nor a photosynthetic phenotype**

*2.1 No growth, germination or macroscopic phenotypes are present in the plastome I variants*

*2.2 Photosynthetic parameters are unaltered in the plastome I variants*

*2.3 Chloroplast sizes, numbers or volumes per cell are unchanged in the plastome I variants*

**3. Correlation mapping**

*3.1 Categorization of wild type and mutant plastomes into classes of inheritance strength*

*3.2 Selection of reference sequences for correlation mapping*

*3.3 Correlation mapping in the wild type plastomes*

*3.4 Correlation mapping in the green variants*

*3.5 Correlation analysis at selected loci*

*3.6. Phylogenetic independence of the correlation mapping analyses*

**4. Repeat structure, sequence evolution and divergence of *accD*, *ycf1*, *ycf2*, and *oriB***

**5. Variation at the chloroplast origins of DNA replication is not responsible for differences in chloroplast competition**

*5.1. Sequence variation in oriB cannot explain differences in inheritance strength*

*5.2 Changes in plastid DNA amounts during development do not correlate with inheritance strength*

*5.3 Nucleoid number and structure is identical in lines with different inheritance strength*

**6. Expression and transcript maturation of *accD* and *ycf2***

**7. ACCase activity in lines harboring chloroplasts of different inheritance strength**

**8. Predictability of inheritance strength based on lipid-levels**

*8.1. Chloroplast inheritance strength is independent of bleaching*

*8.2 The very weak variants III-V1 and III-V2*

*8.3 Classes of inheritance strength employed in the LASSO regression model*

*8.4 Predictability of inheritance strength based on lipid-level data as explanatory variables*

*8.5 The lipid classes DGDG, PG, PC, and PE are enriched for predictive lipids*

*8.6. A model for the predictability of inheritance strength based on lipid-levels*

### 1. Generation of green variants with altered inheritance strength

For functional validation of loci predicted by correlation mapping in the wild types (see Main Text and below), mutagenesis of the strong chloroplast genome I-johSt was conducted using the *Oenothera plastome mutator* (see Materials and Methods for details). Inheritance strengths of the obtained green variants were determined in crosses to the white chloroplast mutants I-chi or IV-delta as pollen or seed parent, respectively (Materials and Methods for details, Fig. 2, Table S10, and Main Text).

The progeny of three seasons were analysed for the two crossing series. From these experiments it appeared that the lines VC1 and V3g (together with the weak wild type IV-atroSt) have very low assertiveness rates in the F1. As already judged by eye, they form a distinct class from all other variants in both crossing directions (Fig. 2; Table S10). For the reciprocal cross, no significant differences from the strong wild type I-johSt were found for the variants V1c, V2f, and V3e, meaning that these variants had the same transmission efficiency as the wild type, although they underwent a mutagenesis approach and carry background mutations. This makes them a particularly valuable material to identify *plastome mutator*-induced mutations that do not affect chloroplast inheritance. All other variant plastids showed a significantly decreased competitive ability from at least one parent when compared to the wild type chloroplast genome I-johSt. In general, the plastome I variants cannot be grouped easily into the classes of Schötz (strong, intermediate and weak). Besides VC1 and V3g forming a separate weak group (see above), their transmission frequencies fall between the strong and intermediate wild types (cf. I-johSt and II-suavG of the I-chi cross; Fig. 2A). Hence, only two classes, a weak and a strong/intermediate one are present in the variants (analysed below in more detail).

The results of the classical experimental set up in which bleached chloroplast mutants are used, could be confirmed in a MassARRAY® approach employed for the crosses of the plastome I variants with the strong wild type plastome I-hookdV as male parent or the weak wild type IV-atroSt as the female parent (see Materials and Methods for details). Due to the detection threshold of the method (5-10%) when I-hookdV was transmitted through the pollen, most variants showed the same or slightly decreased transmission efficiency as their wild type I-johSt, with the progeny having increased amounts of paternal plastid DNA (ptDNA). However, only for VC1 and V3g is the difference of the ratio of paternal and maternal ptDNA in the pool large enough to result in the detection of a significantly lower assertiveness rate (Fig. 2B). The lines behave similarly in the other crossing direction, where most variants seem to be of wild type competitive ability. Again VC1 and V3g can clearly be confirmed as weak lines, while V3c and V3f (which appear as weak to intermediate when contributed by the female), show a higher transmission efficiency than I-johSt. This is the same reciprocal difference that is observed in the classical experimental set up (Fig. 2). Altogether, especially due to the detection limit, the classical approach using bleached chloroplast mutants gives more reliable results and allows a much finer discrimination of transmission efficiencies. Moreover, there is no qualitative difference in the assertiveness rates between the green wild types I-hook/IV-atroSt and their corresponding bleached mutants I-chi/IV-delta. This is in agreement with the classical literature (24) and investigated in more detail below.

### 2. The green plastome I variants do not display impaired growth, altered chloroplast morphology, nor a photosynthetic phenotype

To rule out the possibility that the observed differences in chloroplast inheritance strength result from secondary effects in the green variants we performed several controls: First, we monitored growth behavior of plants with the green chloroplasts of different inheritance strength in the common nuclear background of johansen Standard. Second, to access the physiological status of the material, we measured photosynthesis parameters. Third and last, we performed detailed microscopy to investigate chloroplast size, number per cell and morphology.

#### *2.1 No growth, germination or macroscopic phenotypes are present in the plastome I variants*

To ensure that the green variants are not impaired in development, cultures of johansen Standard plants harboring various variant chloroplasts were compared side-by-side to plants with their strong wild type chloroplast genome I-johSt and the weak one IV-atroSt. It appeared that seeds from all plant lines germinated at 100% within 3 days after sowing (DAS). After transfer to soil, no differences in growth were observed during whole plant development under standard greenhouse conditions. Also no macroscopic phenotype such as altered leaf coloration was observed (Fig. S7).

#### *2.2 Photosynthetic parameters are unaltered in the plastome I variants*

To gain insights into the physiological status of our materials, we determined several photosynthetic parameters and plotted them against competitive ability (Fig. S8). From these analyses it became clear that differences in photosynthesis capability, if present at all, cannot be interpreted as a function of inheritance strength: We could not detect significant differences between plants nor dependencies of inheritance strengths on chlorophyll content per leaf area or for chlorophyll a/b ratio. The latter reflects the ratio of the photosynthetic reaction centers (exclusively binding chlorophyll a) to the antenna proteins (which bind both chlorophyll a and b). Also $F_V/F_M$, the maximum quantum efficiency of photosystem II (PSII) in the dark-adapted state, did not show any changes with inheritance strengths. All measured values were above 0.8 indicating that PSII was intact and that its antenna proteins were efficiently coupled to the reaction centre. There was a minor tendency towards a decrease of leaf respiration in darkness with higher assertiveness rates. However, neither for leaf assimilation rates measured at the growth light intensity of 200 µE $m^{-2}$ $s^{-1}$, nor for assimilation capacity measured under light-saturated conditions, were changes dependent on competitive ability observed. Similarly, for other photosynthetic parameters tested, including leaf absorptance, the chlorophyll a fluorescence parameters qN (non-photochemical quenching, a measure for the thermal dissipation of excess excitation energy in the antenna bed of PSII) and qL (a measure of the redox state of the PSII acceptor side), no clear differences dependent on competitive ability were found.

*2.3 Chloroplast sizes, numbers or volumes per cell are unchanged in the plastome I variants*

To test if differences in competitive ability are a side effect of a putative chloroplast division phenotype, our strong wild type I-johSt was compared to three lines with weak transmission efficiency (VC1, V3g and IV-atroSt). For this, we performed light microscopy using DIC optics at a developmental stage where the first 3-4 true leaves have developed (25 DAS). The chloroplast numbers in spongy mesophyll cells varied between 40-70 plastids per cell. This variance was found in all lines. None of the analysed weak lines showed a significant difference compared to I-johSt in chloroplast morphology, average chloroplast number per cell or chloroplast volume index (length x width$^2$) (20). For chloroplast number per cell, one-way analysis of variance (ANOVA) yielded $p = 0.93$ among all four lines (Fig. S9). Adjusted $p$-values obtained by $t$-test and multiple testing correction in the comparison of each single line with I-johSt again did not uncover significant differences (Table S15). Very similar results were obtained for the chloroplast volume, for which one-way ANOVA gave a value of 0.51 in the comparison of all four lines. Comparing I-johSt with the weak plastomes also did not uncover significant differences, as judged from multiple $t$-testing (Table S16)

**3. Correlation mapping**

As described above, the green variants do not display any phenotype other than an altered inheritance of the chloroplast in crosses. Together with the wild type chloroplasts of different inheritance strengths, this makes them a valuable material to pinpoint molecular loci for chloroplast transmission encoded on the plastome. In contrast to algae or fungi, however, organelle genomes of higher plants or animals are not amendable to linkage mapping (25). Consequently, in these materials, identification of functionally relevant loci can only be based on correlation of a polymorphism within a given sequence interval to a phenotype in a mapping panel. To our best knowledge, this has been done only manually so far (26, 27), which somewhat limits these analyses to a manageable number of organelle sequences, as well as to simple phenotypes, such as the presence or absence of sterility (28). We therefore developed a novel mapping approach that fills this methodological gap. Conceivably, this approach could be applied to map loci conferring cytoplasmic male sterility (28), mitochondrial diseases (29), cytonuclear incompatibility or to analyze adaptive cytoplasms (30-32).

The method is based on Spearman's rank and/or Pearson's correlation (Materials and Methods), with the latter capturing linear dependencies more directly. Since (i) presence or absence of linear dependencies in our data structure is a matter of speculation, and (ii) as a rank-based correlation metric, Spearman correlation yields more statistically robust results that are less influenced by outliers, we have used both approaches. For this, we calculated sequence divergence (total count of nucleotide changes, i.e. SNPs, insertions and deletions) in respect to a reference sequence (see below) for every sequence in an alignment at a given alignment window. The value thus obtained is then correlated with a phenotype. In our case, this is a class of inheritance strength or a percentage value expressing transmission efficiency of a given chloroplast genome (see above and Materials and Methods for details). For example, if the reference sequence represents a strong chloroplast genome and, relative to it, certain weak plastomes contain polymorphisms in the same alignment window, this window is identified as highly relevant for inheritance strength (Fig. 1A; Fig. S17). Subsequently, individual polymorphisms or regions within this window are analyzed separately (Fig. 1B). Since full organelle genomes are analyzed, more than one relevant site can be identified. However, as for any other association mapping approach, the presence of two or more genetically independent loci that confer the same phenotype can complicate the conclusion. Perfect correlation coefficients of 1 or -1 might not be achievable at a single site.

Another common challenge of all genome wide association studies, a lack of resolution to identify functionally relevant loci due to linkage disequilibrium, is especially challenging in a non-recombining system such as an organelle genome. Here, linkage to phylogeny is extreme and even absolute correlation at a single site may be due to genetic hitchhiking via linkage disequilibrium, and not necessarily due to functional relevance. However, also the opposite is true. Non-independence from the phylogeny does not necessarily stand against functionality. Ideally, this problem can be partially circumvented, if phylogenetic independence of the correlation between a given trait (e.g. inheritance strength) and a sequences window can be shown. This indicates functionality, since trait and sequence windows must have evolved at least twice independently. Phylogenetic independent contrasts (PIC) or related methods such as phylogenetic generalized least squares (PGLS) test this null-hypothesis, which motivated us to implement a phylogenetic control in our correlation mapping approach (SI Materials and Methods). However, since a lack of sufficient independence from phylogeny does not stand against functionality (see above), in our context, these methods are only informative for the subset of cases where independent evolution indeed happened and cannot replace experimental verification of predictive loci.

*3.1 Categorization of wild type and mutant plastomes into classes of inheritance strength*

The datasets that measure inheritance strength of wild type chloroplasts or the green variants were either obtained from the literature or produced in this work. They represent percentage values of heteroplasmic seedlings in an F1 generation that reflect inheritance strength of a given chloroplast genome (Tables S7 and S10; see above). The numbers can be directly applied to Spearman's/Pearson's correlation. If datasets of more than one crossing series are to be combined, clustering of the crossing data into classes is necessary.

For the wild type plastomes, we used the original data of Franz Schötz, where two sets of crosses "biennis white" and "blandina white" are available (33, 34) (Table S13; Materials and Methods); inheritance strength of 25 wild type chloroplasts was determined using these previously described tester lines. Clustering of the two datasets with the *k*-means algorithm using the optimal number of centers ($k$ = 3) confirmed the original classifications suggested by Schötz, with the exception of the I-bauriSt and II-corSt plastomes (Fig. S16A; for details see Materials and Methods). These plastomes were borderline genotypes in Schötz's classification system, and according to our data, they might be reassigned. Besides these minor discrepancies, clustering supports the presence of the three classes of inheritances strengths (strong, medium and weak) among the wild type plastomes of *Oenothera*, as previously described.

The clustering of the green variants is less clear. When data from the I-chi and IV-delta crossing experiments are combined, the pamk function identified $k$ = 2 as the optimal number of clusters, clearly separating the weak from the stronger materials (Fig. S16B). However, finer clustering of the stronger variants leads to ambiguous class membership. This is likely due to the higher variation in the IV-delta crosses compared to the I-chi crosses (Fig. 2 and above). This seemingly weakens the combination of the two datasets. Finally, we chose four clusters to classify the variants: (i) At this number of clusters, individual samples swap at the lowest rate between the clusters, when *k*-means (a clustering method that includes a random element) was applied repeatedly. (ii) This number of classes best reflects the transmission abilities of the variants, since many variants are somewhat in between the strong and the intermediate genotypes (see positions of the strong and intermediate wild types I-johSt and II-suavG in the I-chi crosses; Fig. 2A and above). This justifies the definition of a fourth class and material of that kind does not exist in the wild types (cf. Fig. S16A vs. 16A). Hence, based on the wild type classes, the variants are classified as follows: class 1 = strong, class 2 = strong to intermediate, class 3 = intermediate, and class 4 = weak.

*3.2 Selection of reference sequences for correlation mapping*

For correlation mapping in the wild types the chloroplast genome of I-hookdV was chosen as a reference for two reasons: The plastome is the strongest one known (Table S13), but also the most derived one as judged from phylogenetic analyses (Fig. S1, SI Materials and Methods). Based on this it is a natural choice, since every polymorphism in respect to this reference should potentially make a chloroplast genome weaker. A similar argument applies to the reference I-johSt in the correlation analysis of the green variants. This plastome is the progenitor of these lines.

*3.3 Correlation mapping in the wild type plastomes*

Pearson's correlation generally identified more windows than Spearman's, but both predict essentially the same regions relevant for inheritance strengths. Interestingly, there was no notable difference between the methods if either *k*-means classes or the "biennis/blandina white" crossing data were used for correlation (Fig. 1A, Fig. S1, and Dataset S1). Largely based on theoretical considerations (presence of three clearly ranked classes in the wild types and stronger experimental base if the "biennis white" and "blandina white" crossing experiments are combined; see above), we discuss here Spearman's rank correlation to *k*-means classes in more detail. According to the latter, sequence windows in the *ycf1* and *ycf2* genes (between alignment positions 99011-100000 and 134641-135640) show nearly absolute correlation to inheritance strengths (*rho* = -0.99, $p < 0.0005$; Dataset S1). In both genes, the correlation oscillates from *rho* = 0.86 to -0.99 ($p < 0.0005$), and the positive and negative correlation should be interpreted as equally important. Another nearly absolute correlation (*rho* = 0.98, $p < 0.0005$) was measured in alignment windows containing the promoter, 5'-UTR and 5'-end of *accD* (positions 63501-64760). A window further upstream containing the same features (positons 63391-64490) also correlates with *rho* = 0.96, $p < 0.0005$ (Fig. 1B). However, highly significant correlations of 0.96 were also found in intergenic regions of photosynthesis genes and/or tRNA genes, for example between *ycf3* and *psaA* (encoding a photosystem I assembly factor and core subunit, respectively) (35). In addition, significant correlations were measured from the spacers of the photosystem II and cytochrome $b_6f$ subunit genes *psbE* and *petL*, and in a sequence interval contacting *trnR-UCU* and *trnG-UCC*. In contrast, no significant correlation was observed for *oriA*. For *oriB*, three sequence windows (partially) containing the *oriB* correlate with 0.90, 0.88 (both $p < 0.0005$) and 0.81 ($p< 0.005$). If Pearson's correlation to *k*-means classes is applied to the wild type data, the described pattern can be reproduced but more windows with significant correlation are identified (Fig. 1 and see above). The highest observed Pearson correlation in the wild type dataset is $r = 0.96$ ($p < 0.0005$) in a sequence window again containing the promoter, 5'-UTR and the 5'-end of *accD* (Fig. 1B, Dataset S1).

*3.4 Correlation mapping in the green variants*

When correlation mapping results are compared between the wild type and the green variants, the most striking difference in the variants is the loss of significance after *p*-value adjustment for Spearman's but not for Pearson's correlation. Here, windows with significant correlations were obtained (cf. Fig. 1 vs. 3 and Fig. S1 vs. S10, Dataset S1). This is probably because the rank-based Spearman correlation is less influenced by the VC1 and V3g data points. These two single genotypes, however, form the weak and, therefore, most predictive class, whereas the other variants do not differ noticeably from their wild type progenitor (cf. Fig. 2 and Fig. S16B; also discussed above). This leads to a relatively weak correlation to inheritance strengths which appears to be an under-estimation and a consequence of the multiple testing correction (> 13,000 tests). The weak correlations also contradict the genetic observations, which clearly indicate that the plastomes of the green variants must contain mutated loci for inheritance strength. A similar argument applies to correlation of the *k*-means classes in the variants. As discussed above, definition of these classes is less clear than in the wild type, which weakens their predictive power. We therefore think that the Pearson correlation of the I-chi crosses (which yield a better resolution than the reciprocal IV-delta crosses; Fig. 2 and above) represents the best approach to identify the relevant loci that alter inheritance strength in this material. Notably, this approach yields the most significant correlations, but all approaches (including Spearman) identify the same regions in the plastome with the highest correlation values (Fig. S10, Dataset S1).

In the variants, Pearson's correlation of the I-chi crosses predicts a sequence window in the 5'-end of *accD* as significantly correlated to inheritance strengths ($r = 0.78$ and $p < 0.005$). The strongest correlation for this dataset is observed for the 5'-UTR of *ycf2* ($r = 0.91$, $p < 0.0005$). In addition, a highly repetitive region in the coding region of the same gene also shows good correlation values ($r = 0.71$; $p < 0.05$; Fig. S10, Dataset S1). Two insertions/deletions (indels) in *ycf1* are also significant ($r = 0.61$; $p < 0.05$). They represent a single insertion and a deletion in the weak variant VC1, located relatively close to each other (see below). The functional relevance of the two mutations for inheritance strength can be questioned, however. Since the second weakest variant of the dataset displays a wild type *ycf1* sequence (Fig. S6A), the above described mutations are likely a result of the large amount of background mutations present in VC1 (see Materials and Methods for details). For the same reason, a contribution to the phenotype by *oriB* can be excluded ($r = 0.41$, $p = 0.25$) in the variants. Taken together, our results narrow down the regions identified in the wild types to the two genes *accD* and *ycf2*. All other mutated loci in the variants seem to be of minor importance.

#### *3.5 Correlation analysis at selected loci*

When correlation mapping is applied to selected loci within the above identified alignment windows, the general observation is that correlation values drop to some extent (cf. Dataset S1). This is probably best explained by looking at the highly correlating sequence intervals spanning the promoter, 5'-UTR and 5'-end of *accD* in the wild types (Fig. 1B). When analyzed as functional units (promoter/5'-UTR region and protein N-terminus; Fig. S4), correlation of the individual segments (promoter/5'-UTR region: $r = 0.80$ or $rho = 0.74$; $p < 0.005$ for both; full N-terminus: $r = 0.78$ or $rho = 0.60$; $p < 0.005$ or $p < 0.05$) is much lower than for the original sequence intervals ($r = 0.94$ or $rho = 0.96$ and $r = 0.96$ or $rho = 0.98$ with $p < 0.005$ for all), which led to the identification of these regions. As discussed below, experimental evidence is available that promoter/5'-UTR and N-terminus interact to affect the inheritance phenotype, while the individual regions display weaker correlations.

In spite of these complications, to get an impression of how well certain coding or promoter/5'-UTR regions, as well as segments of *oriB* correlate with inheritance strengths, we calculated correlation values for *accD*, *ycf1*, *ycf2*, and *oriB* for polymorphisms that are present in both wild type and the variants (Figs. S4-S6, Dataset S1). We also included two prominent sites in the *ycf2* gene present in the wild type, for which we found no mutation in the variants. Please note that at three sites (AccD N-terminus, the AccD site 2 and the *ycf2* promoter/5'-UTR region) in addition to Person's correlation, the Spearman's correlation analysis yields significant correlations in the variants. This is in contrast to the whole plastome approach described above, where *p*-value corrections due to multiple testing were applied.

The best correlating region in both sequence sets (wild type and variants) is site 2 of the AccD N-terminus (Fig. S4B). Its prediction is extremely robust in that significant Pearson's and Spearman's correlations were obtained for all crossing series and *k*-means classes (Fig. S4, Dataset S1). Less clear is the contribution of *ycf2*. In the wild types, the *ycf2* site 1 and site 2, but not site 3 can be associated with inheritance strength, but in the green variants, site 3 exerts the most influence on the competitive ability of chloroplasts (Fig. S5B and below).

In summary, the refined analyses at selected loci clearly confirm the contribution of *accD* on inheritance strengths and might have even identified the most important region. It also shows that *ycf2* may contribute to the phenotype. Without chloroplast transformation in the evening primrose, a technology currently not available, the influence of the individual sites remains speculative.

*3.6. Phylogenetic independence of the loci predicted by correlation mapping*

To address the possible impact of phylogenetic non-independence on the association between total sequence divergence and inheritance strength, we implemented a phylogenetic control in our correlation mapping analysis. In principal, we expect loci for inheritance strength to be to some extent independent of the phylogeny, since plastome phylogeny in *Oenothera* is (((I,II) III) IV), whereas inheritance strengths follows the distribution (((I,III) II) IV) (Fig. S1).

For the wild type dataset det, for which it is easy to reconstruct a phylogeny, we performed this test using a phylogenetic generalize least squares model (PGLS), statistically equivalent to phylogenetic independent contrasts (PIC) but with a much more flexible implementation (36). This allowed for relatedness between each of our 14 lineages to be accounted for in the association between divergence and inheritance strength, however, both PIC and PGLS analyses rely on the assumption of a continuously distributed response variable (12, 37). Unfortunately, our discrete *k*-means clustering approach that best reflects the joint inheritance strength of our two sets of crosses, "biennis white" and "blandina white" (see above) does not fit this assumption. Instead, we conducted the analysis with continuous measures of inheritance strength, % of variegated seedlings from the biennis and blandina crosses, despite these frequencies not always assigning the same rank order to our wild-type chloroplast lineages (Table S7).

Consequently, for the blandina crosses, this analysis was not informative. Before *p*-value correction, only some minor sequence variation around *trnM-CAU* and *trnV-UAC*, as well as the small single sequence (SSC) and the inverted repeat A ($IR_A$) junction were found to be phylogenetically independent (Dataset S1). The latter is displayed at the very end of the linear plastomes map in Fig. S1, highly divergent not only in evening primroses and likely depleted of functional sequences motifs (8). For the biennis crosses, the SSC/$IR_A$ junction and five new regions were identified, four of them again comprising minor sequence variation in intergenic regions, not being able to explain the inheritance phenotypes (Dataset S1 and S2). The fourth region, however, is *ycf2* and here two of the previously predicted sites (see above) were shown to be significant after phylogenetic correction (site 2 and site 3; $p < 0.05$ and $p < 0.05$, respectively; Fig. S1, Dataset S1). However, they did not survive statistical significance after false discovery rate correction, most likely because of a lack of power, i.e. with a small tree there are insufficient numbers of independent events along a relatively small phylogeny to detect the significant association [only one event, comparing phylogeny (((I,II) III) IV) to inheritance (((I,III) II) IV)]. In other words, the phylogenetically-controlled association analysis alone does not withstand statistical scrutiny.

Interestingly, *acc*D is missing from the phylogenetic independent regions. This might support the view of at least two independent loci determining inheritance strengths, one of which (*ycf2*) is not linked to the phylogeny. In summary, we think that implementation of PGLS or related methods has the potential to significantly improve the correlation mapping method, although this needs to be explored with a much broader base.

**4. Repeat structure, sequence evolution and divergence of *accD*, *ycf1*, *ycf2*, and *oriB***

The four genes or loci partially span rapidly evolving regions of the *Oenothera* plastome that are characterized by large repetitive regions. Those can be of up to 1 kb in size as is the case for site 3 in the *ycf2* gene. They are comprised mostly of tandem or direct repeats (and less pronounced palindromes or inverted repeats) as described earlier (8). Due to their repetitive nature, these regions are very prone to replication slippage (2, 38) and sequence divergence at these regions substantially contributes to the overall sequence variation of the *Oenothera* chloroplast DNA (Greiner et al. 2008. Fig. 3 therein) (8). The presence of repeats also makes them a preferred target of the *plastome mutator* allele (39, 40).

Sequence evolution is extremely fast at these repeats. In case of the repetitive regions of *accD* and *ycf1* phenotypically neutral spontaneous mutations were isolated repeatedly at very similar sites (2). Moreover, the *oriB* (which is essentially located in the *rrn16 - trnI-GAU* spacer) is used as a hypervariable marker allele that allows discrimination among a huge variety of *Oenothera* strains (3). The repeat structure of the *oriB* region was analysed earlier and is comprised of 7 direct repeat classes that can be divided into various subtypes (39, 41) (and below). In the *accD* gene mostly tandem or direct repeats span the promoter/5’-UTR and N-terminal region (Fig. 1); all three of these segments are considered to contribute to the regulation of the gene (42-46). In fact, sequence variation induced by theses repeats is so high that upstream of the *accD* start codon, a window of about 1.4 kb cannot be aligned between the weak plastome IV and the stronger plastomes I-III (Dataset S2). This is to some extent also observed for site 2 in the N-terminus of the wild type AccD. In plastome IV major portions of this site is missing and about half of the remaining sequence is polymorphic (Fig. S2A). In the *ycf2* coding sequence, the most prominent repeats are present in sites 2 and 3. At the first site the number of PEKRKEKK tandem repeats can be correlated with inheritance strengths in the wild types, but not in the variants. The situation is reversed for site 3, in which tandem repeats exist as two subtypes 5’-GAGGAAGtAGAAGGGACAGAA-3’ and 5’-GAGGAAGgAGAAGGGACAGAA-3’ associated with a GAT linker, and correlate with inheritance strengths in the variants, but not in the wild types (Fig. S2).

### 5. Variation at the chloroplast origins of DNA replication is not responsible for differences in chloroplast competition

As elaborated above, our correlation mapping already points to a connection of lipid biosynthesis and chloroplast competition. However, one might still argue that *a priori* differences in the origins of replication are the simplest mechanistic explanation for organelle competition. At least some evidence supporting this claim is available for yeast and *Drosophila* (47-49). The location and repetitive nature of *oriB* in evening primroses (see above) are reminiscent of the non-coding displacement loop (D-loop) of metazoan mitochondrial DNA (mtDNA). In many animal taxa the D-loop is the most variable sequence of mtDNA and is in the proximity of tRNA or rRNA genes (47, 50, 51).

#### *5.1. Sequence variation in oriB cannot explain differences in inheritance strength*

Previous work in the evening primrose did not support an involvement of the origins of replication in chloroplast competition. First, the number of D-loop initiation sites (i.e. *oris*) does not differ between weak and strong plastomes and their locations in the chloroplast genome is identical (52). Second, in a previous association mapping study that investigated the hypervariable repeat region of *oriB*, a short repeat series was identified as the sole determinant that could explain the difference between the strong and intermediate plastomes I, III and II on one side, and the weak plastome IV on the other side (41). The sequence (5’-ACGACACGACGATTAGATTAGCTCATTGGTAGGACGACGATTAGCTCATTGGTAGGACGACG-3’) is 62 bp in size and is capable of forming of weak hairpins. Our study, analyzing a greater number of plastome sequences, confirms the absence of this sequence in the weak plastome IV. However, in none of the green plastome I variants with altered inheritance strength is the sequence partially or fully deleted. Moreover, the very weak variants of plastome III (Main Text and see below) do not carry a single mutation in one of the two origins of replication (Dataset S2). We therefore do not think that a genetic determinant within the *oriB* of *Oenothera* is able to explain the huge differences observed in competitive behavior.

To substantiate this view, also on the level of DNA, we investigated the dynamics by which ptDNA increases during plant development in more detail. In addition, we analyzed chloroplast nucleoid structure and number per cell.

*5.2 Changes in plastid DNA amounts during development do not correlate with inheritance strength*

To investigate if differences in ptDNA increase during development and/or if changed ratios of plastid/nuclear DNA are able to explain chloroplast competition, we performed quantitative real-time PCR. In general, plastid DNA amounts are not static during ontogenies (18, 53). They increase as leaves grow, starting from 0.4% in meristematic tissue to more than 20% in mature leaves (17). If differences were observed in DNA abundance during development in different *Oenothera* lines harboring chloroplasts with different inheritance strengths, it might hint towards an aspect of DNA replication such as replication speed as an underlying mechanism for plastid competition. To monitor this process we analysed total DNA of the johansen Standard strain equipped with the strong and the weak wild type chloroplast I-johSt and IV-atroSt, respectively. In addition, we included selected lines of our plastome I variants: V1c, V2a, V2g, and V3e (all strong), V3c (strong to intermediate), V3d (intermediate) and VC1, V3g (both weak). Plant tissues of different developmental stages were analysed, seedlings 5 DAS (when seeds have just germinated and cotyledons have developed), plantlets 21 DAS (after development of the first two true leaves), and the second true leaf of young rosettes 32 DAS. ptDNA amounts were calculated relative to I-johSt (5 DAS) per one haploid genome (for details see SI Materials and Methods).

Depending on the plastome target region, at 5 DAS a small increase of ptDNA amounts was observed in the lines V1c, V3e, V3c, VC1 and V3g. These differences, however, are not significant. At 21 DAS the weaker variants V3c, V3g and VC1 showed an increase in relative ptDNA amounts compared to wild type I-johSt, but only for the target *ndhI* which was again not significant. In general, from 5 to 21 DAS only a minor or no increase of plastid DNA amount was observed for each particular line and each plastid target, while in the young rosette at 32 DAS the ptDNA amount doubled (Fig. S11). These results echo previous work in *Arabidopsis* and sugar beet (17, 53). Since the same results were obtained for plant lines carrying strong and weak plastids, no developmental difference in ptDNA copy numbers correlates with differential transmission efficiencies.

In summary, all minor increases in ptDNA amount are not significant nor do they correlate with transmission efficiencies nor with the DNA variations described previously (Fig. S6B). Moreover, no differences can be detected in IV-atroSt compared to wild type I-johSt, although plastome IV is the weakest of all genotypes tested. Therefore, the ptDNA amounts in vegetative tissues do not indicate different replication speeds, suggesting that replication *per se* is not the underlying mechanism for different transmission efficiencies.

*5.3 Nucleoid number and structure is identical in lines with different inheritance strength*

Under the premise that ptDNA amounts are constant, there is still the possibility that strong and faster replicating plastomes have altered numbers of nucleoids, which could impact their ability to divide. To exclude this possibility we quantified nucleoids in the central laminal region of the first true leaf 25 DAS. After staining with DAPI, nucleoids were clearly visible as small dots with their fluorescence sharply delimiting them from the dark cellular background, even when forming tight associations like clumps or threads (Figs. S12 and S13). One strong (I-johSt) and three weak lines (V3g, VC1, and IV-atroSt) were investigated. The mean number of nucleoids per chloroplast ranges between 17.7 and 18.1 with no significance differences between the lines. One-way ANOVA gave $p = 0.53$; multiple $t$-tests comparing I-johSt with each of the weaker plastomes did not point to significant differences as well (Fig. S14 and Table S14). Moreover, we did not observe any difference in nucleoid morphology between the lines.

**6. Expression and transcript maturation of *accD* and *ycf2***

Since the polymorphisms in *oriB* cannot explain differences in competitive ability, we investigated the accumulation of *accD* and *ycf2* transcripts. For this, we used leaves of plants 26 DAS from the lines I-johSt, V1c, V3e, V2g (all strong), V3c (strong to intermediate), and VC1, V3g and IV-atroSt (all weak; Fig. S14). A probe specific for the conserved C-terminal part of *accD* detected the mature transcript at about 3 kb. No differences in transcript accumulation were observed between I-johSt and the plastome I variants. However, for IV-atroSt two additional bands running below the mature transcript were present. Moreover, the mature transcript clearly over-accumulated in this weak, but phylogenetically more distant plastome. This transcript over-accumulation appears to be a result of the high sequence variation observed in the *accD* promotor/5'-UTR that strongly correlates with inheritance strength (see Fig. 1, Fig. S4A and above). A similar analysis was conducted for *ycf2*, where again a probe specific for the C-terminal part of the gene detected the mature transcript at the expected size of about 9 kb. The very small differences in size between the lines perfectly mirrors the occurrence of in-frame deletions in the lines IV-atroSt, V3c, V3g, and VC1 (Dataset S2). In IV-atroSt as well as in the plastome I variants, no difference in accumulation of the mature transcript compared to I-johSt was found. However, transcript stability/processing seems to vary between the strong plastome I-johSt and the weak IV-atroSt. Interestingly, whereas the strong variants V1c, V3e and, to same, extent V2g showed exactly the same transcript pattern as the wild type, the weak variants showed a pattern more similar to IV-atroSt. This indicates a correlation between transmission efficiency of mutations in site 3 of *ycf2* (cf. Fig. S3), which might result from altered mRNA degradation and/or processing.

**7. ACCase activity in lines harboring chloroplasts of different inheritance strength**

As the above described analysis indicates an involvement of *accD* and/or *ycf2* in the inheritance phenotype, we decided to determine ACCase activity in our lines. From these measurements, it appeared that the strong variants (V1c, V3e, V2a, and V2g) display a similar or even lower ACCase activity than their wild type I-johSt. The same holds true for the strong to intermediate or intermediate genotypes (V3c and V3d). In the weak materials, a 2-3 fold increase of ACCase activity is observed for VC1 and IV-atroSt, although V3g shows wild type enzyme activity (Fig. 4A). Although there is no simple linear correlation between inheritance strengths and ACCase activity, the strong increase in VC1 and IV-atroSt is hard to ignore. In fact, both inheritance strength and ACCase activity seem to depend on the particular mutation pattern: (i) Mutations in *ycf2* seem to influence ACCase activity, as judged from the variant V3e that is wild type for the *accD* segments but is mutated in *ycf2* (Fig. 4A, yellow box). (ii) Larger mutations in the AccD N-terminus have higher ACCase activity, whereas the presence of a more diminutive AccD N-terminus correlates with lower activity (Fig. 4A, cf. blue boxes vs. the remaining pattern). (iii) There must be an influence of *ycf2* on inheritance strength (cf. Fig. 4A, green boxes associated with the weaker materials). Hence, if ACCase and/or Ycf2 result in altered levels of lipids, one would expect that lipid composition is predictive of inheritance strengths.

**8. Predictability of inheritance strength based on lipid-levels**

To test for predictability of inheritance strength from lipid level data, we analyzed 16 chloroplast genotypes of different inheritance strength in a LASSO regression model (Table S3; Materials and Methods). Since chloroplast inheritance strength is independent of photosynthetic competence (see below), we included pale lines. The aim was to enrich the lipid signal responsible for inheritance strength, i.e. to deplete for the structural lipids of the thylakoid membrane (54); also see Main Text. Namely, we used our bleached *psaA* mutants I-chi and IV-delta impaired in photosystem I assembly, as well as the pale green *virescent* genotypes III-lamS, III-V1 and III-V2 (Tables S3 and S9; Fig. S15). Such materials were previously shown to have perturbed thylakoid membrane formation (55-58). Moreover, as elaborated in the following chapters, we could confirm the independence of a pale phenotype from inheritance strength with these plastomes.

*8.1. Chloroplast inheritance strength is independent of bleaching*

Intuitively one might expect that bleached chloroplast mutants would be less successful in crosses than their corresponding green wild types. However, previous analyses in evening primroses showed that differences in chloroplast inheritance strength are largely independent of the chloroplast mutant used for the analyses (59, 60). At least for *Oenothera*, it is therefore generally accepted that mutations in a chloroplast genome that result in bleaching essentially do not affect chloroplast assertiveness rates (24) (also see Fig. 2A,C vs 2B,D and above). Due to technical limitations, however, this hypothesis was never tested directly. Since closure of this gap is of general relevance for this work, and to provide further evidence that chloroplast inheritance strength is largely independent of the photosynthetic status of the chloroplast, we directly compared the wild type chloroplast I-hookdV and its bleached derivative I-chi, as well as IV-atroSt and the corresponding mutant IV-delta. For this, we investigated appropriate F1 populations crossed to the chloroplast genomes I-johSt, VC1, and V3g with the MassARRAY® system (Fig. S18; see Materials and Methods for details on the material). As expected, transmission efficiencies had the same range for nearly all six pairs of crosses under investigation. Only in one cross with VC1 as a mother, the bleached mutant I-chi actually behaved stronger than its corresponding green wild type.

Taken together, we could confirm that chloroplast assertiveness rates are independent of photosynthetic capability. Moreover, these results make it very unlikely that the differences in inheritance strengths observed for the mutated plastome I variants (all sharing a green phenotype), are due to a secondary effect.

*8.2 The very weak variants III-V1 and III-V2*

While the plastome I variants and their wild type I-johSt are native in and compatible with the nuclear background of the johansen Standard race, III-lamS and its *plastome mutator* variants III-V1 and III-V2 are foreign and incompatible in this background, meaning that tissues carrying them do not develop a normal green color (Materials and Methods, Fig. S15, Table S11). The wild type III-lamS plastome appears to be strong, as judged from crosses to I-johSt as pollen donor, its derivative variants III-V1 and III-V2 are weak (cf. Fig. 2A vs. Fig. S19; cf. Table S10, S11 and S13) (60). Although the fraction of plants showing biparental inheritance in the crosses of III-V1 and III-V2 to I-johSt as pollen donor are somewhat low (37.3% and 38.0%, respectively) for a combination of a weak and a strong plastome (Fig. 2, Table S10) (24), the striking difference of these crosses to all other crosses described is that some seedlings contain only paternal chloroplasts (Fig. S19; Table S11). As mentioned previously, biparental inheritance in the evening primrose shows maternal dominance, in which progeny are either homoplasmic for the maternal chloroplast or heteroplasmic for the maternal and the paternal chloroplasts, but they are never homoplasmic for the paternal one. The appearance of homoplasmic offspring having the paternal chloroplast in the III-V1/III-V2 crosses to I-johSt is the only reported case in the evening primrose where an exception to maternal dominance occurs. This justifies the definition of a new inheritance class for these plastomes.

*8.3 Classes of inheritance strength employed in the LASSO regression model*

To predict chloroplast inheritance strength from lipid-level data, the genotypes of the plants needed to be ranked according to their inheritance strengths (Materials and Methods; Table S3). For the green variants (V1c, V2a, V2g, V3e, V3c, V3d, VC1, and V3g) and the wild types I-johSt and IV-atroSt, the existing *k*-means classes 1 - 4 already employed in our association mapping approach were used (see above). The remaining plastomes (I-chi, IV-delta, I-hookdV, III-lamS, III-V1 and III-V2) were rendered consistent with this framework based on the classification of Schötz and our own data. This adds the plastome I-chi, its wild type I-hookdV and III-lamS to the strong class 1. The mutant IV-delta was placed into the weak class 4. As a result of the exceptions to maternal dominance, when III-V1 and III-V2 were seed parents, these plastomes are placed in a new class 5 (very weak). Taken together, the 16 genotypes are classified into five classes of descending inheritance strengths (strong = 1, strong to intermediate = 2, intermediate = 3, weak = 4, and very weak = 5). Material in class 1 and class 4/5 are over-represented, since (as for the inclusion of the bleached material; see above) we expect to enhance the signal for predictive lipids (Table S3)

*8.4 Predictability of inheritance strength based on lipid-level data as explanatory variables*

The rationale of the predictive approach is as follows: To test for predictability of inheritance strength based on lipid-level data, a linear model (LASSO) was trained and its performance tested in a cross-validation setting on two randomly selected genotypes with differing inheritance strength (see Materials and Methods). If the proposed regression model has predictive power, the actual inheritance strength-values associated with the two test genotypes should be positively correlated with their predicted ones. Note that for each genotype repeated measurements were available. Thus, regression was performed over more than two points and correlation coefficients could assume absolute values differing from 1. Testing was done in a cross-validation setting, i.e. the two test genotypes were not included in the model training. This procedure was repeated 100 times, with each run corresponding to two new randomly selected genotypes of differing inheritance strength and all others used for model training.

If, indeed, inheritance strengths can be predicted based on lipid levels, on average, a positive correlation (Pearson correlation coefficient, *r*) between actual and predicted inheritance strength-values of the two test set genotypes should be obtained. To test for this outcome, 100 Pearson correlation coefficients are classified as positive (success) or negative (failure). Then, they were compared to the null hypothesis of no predictive value, corresponding to a 50% chance of obtaining a positive correlation and significant deviations from this expected probability tested by performing a binomial test.

*8.5 The lipid classes DGDG, PG, PC, and PE are enriched for predictive lipids*

From the 100 cross-validation runs, using the combined dataset from three independent experimental series (Table S3), a median Pearson correlation coefficient (*cvR*) between actual and predicted inheritance strength values of $cvR_{median}$ = 0.7 was obtained (Fig. 4B) with 82 being positive, i.e. successful predictions. This corresponds to $p_{binomial} = 2.17 \times 10^{-9}$ vs. the null hypothesis of 0.5 (no predictive value). Thus, lipid levels proved indeed predictive relative to inheritance strength.

Individual lipids were ranked with regard to their predictive value based on the coefficients by which they entered the regression model (Fig. 4C; Table S4). Averaged over all 100 cross-validation runs, 20 lipids/molecules were identified as predictive as judged by their average absolute weight. They were considered predictive if their absolute average weight was greater than one standard deviation (SD = 0.7) of the average weights of all 102 lipids/molecules. The individual predictive lipids belong to the membrane lipid classes MGDG, DGDG, PG, PC, and PE, as well as to the storage lipid class TAG. Among those, DGDG, PG, PC, and PE were found enriched (odds ratio > 1), albeit statistical significance could not be established (Table S3).

The classes MGDG and DGDG mostly represent plastid lipids located in both the thylakoid membrane and the envelope. PGs and PCs are found in plastid and extraplastidial membranes, although for the chloroplast, PCs are specific to the envelope. PEs are present in the plasma and mitochondrial membranes and, as the storage lipids TAG, essentially absent from chloroplasts (61).

*8.6. A model for the predictability of inheritance strength based on lipid-levels*

Taking into account the data above and of Fig. 4, we propose the following model to explain how certain changes in lipid abundance influence inheritance strengths: Increased activity of acetyl-CoA carboxylase in the chloroplast (Fig. 4A) increases fatty acid concentrations and subsequently fatty acid export to the endoplasmic reticulum (ER). The combination of increased fatty acid synthesis and export leads to an upregulation of the eukaryotic lipid biosynthesis pathway in the ER (62). This is seen in increased amounts or shifted proportions of diverse lipid classes, including phospholipids or storage lipids. Since PC is the dominant phospholipid class of the chloroplast outer envelope (63) those changes affect the structural and physiological properties of the envelope. It in turn impacts chloroplast division and/or stability processes, thus ultimately determining inheritance strengths (see Main Text). Conceivably, the Ycf2 protein, which is located in the envelope (64), might be responsive to changes in the lipid composition of the envelope and/or ACCase activity, thus influencing growth and division of the chloroplast. The observed shift in the proportions of storage lipids or other changes in the extraplastidial lipidome (cf. PEs or TAGs in Fig. 4C) might occur in response to altered fatty acid pools, although especially the storage lipids are likely not relevant for the variation in chloroplast inheritance strengths (see Main Text).

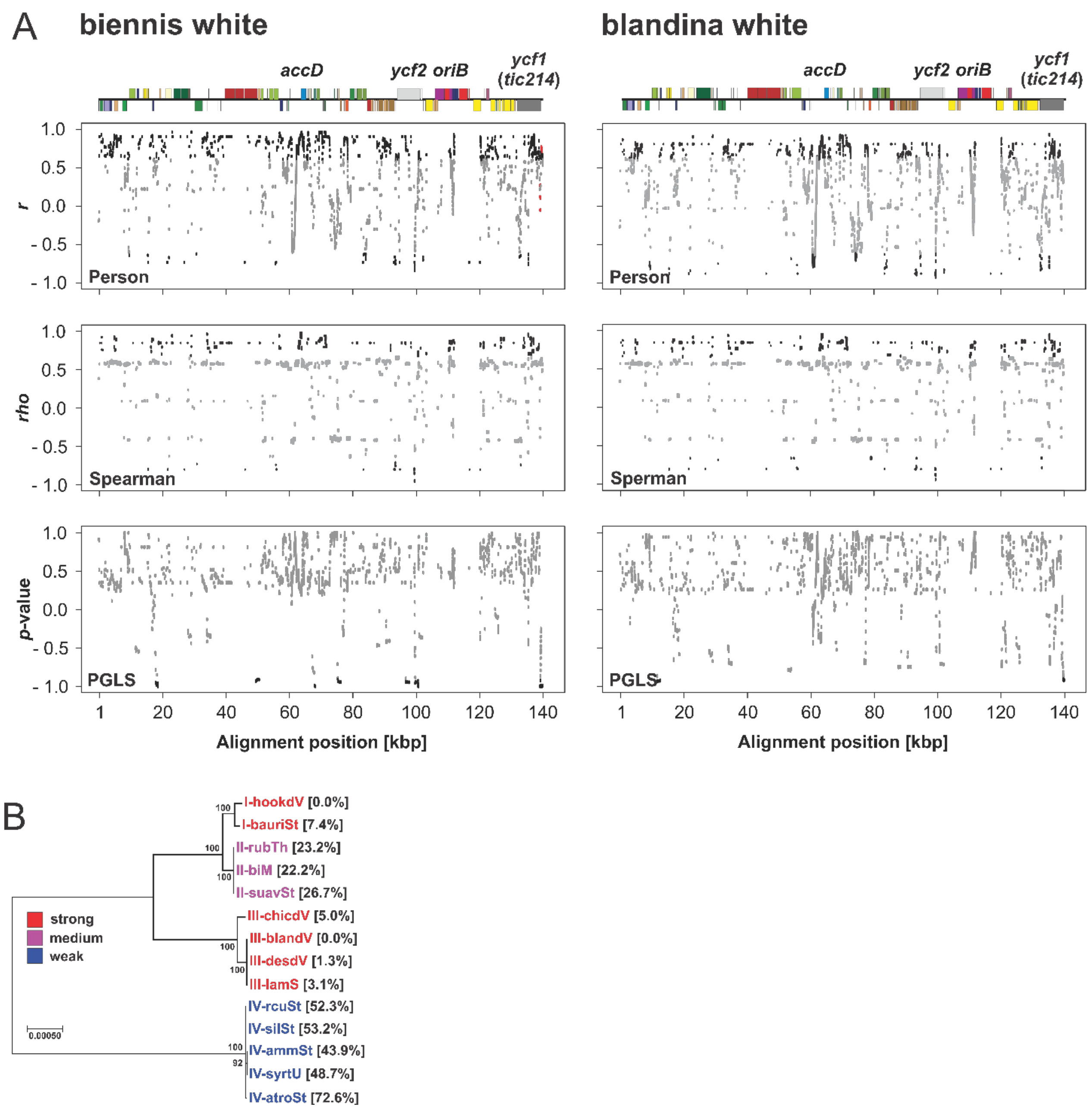


**Fig. S1.** Correlation mapping and phylogenetic independence of predicted sites in the wild type chloroplast genomes using the "biennis/blandina white" crossing data. (*A*) Pearson's correlation, Spearman's correlation and PGLS. Relevant genes or loci with significant correlation are noted on the linear plastome maps above. Non-significant correlations before (PGLS) or after (Pearson's/Spearman's correlation) *p*-value adjustment ($p > 0.05$) are displayed in grey. Alignment windows whose corrected *p*-values of the PGLS analyses still remained significant after correction are marked in red in the original Pearson/Spearman correlation mapping plot. For details see SI Text. (*B*) ML tree of the 14 wild type chloroplast genomes used for the analysis above and their inheritance strength relative to "biennis white" (cf. Table S5). Tree is drawn to scale, with branch lengths measured in the number of substitutions per site. Numbers at branch points represent bootstrap values.

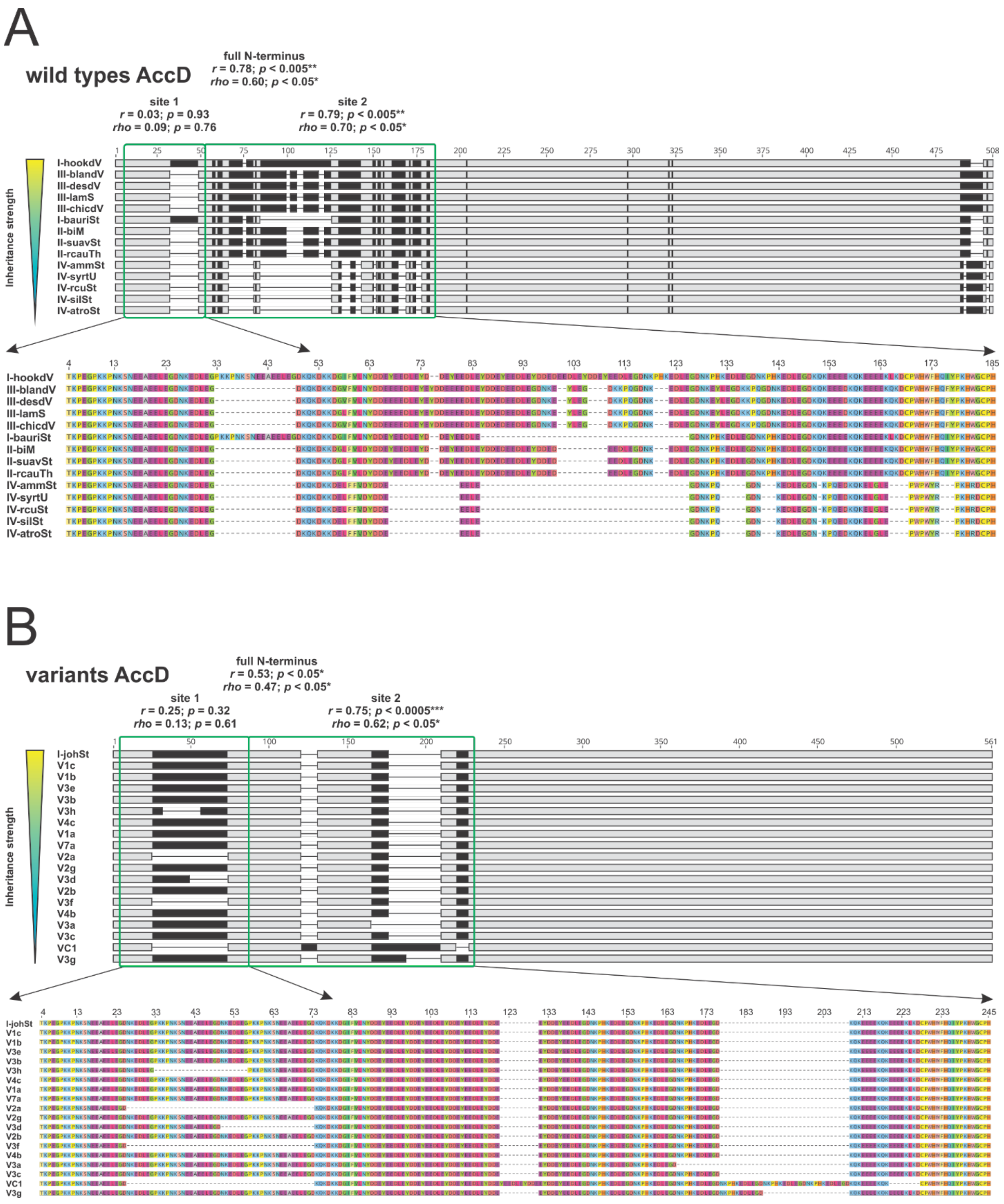


**Fig. S2.** Amino acid sequence of the AccD N-terminus and correlation to inheritance strength (Pearson's/Spearman's correlation to *k*-means classes for the wild types and to I-chi crosses for the variants). Individual sequences are sorted according to their competitive ability. Polymorphic regions are indicated in in black, alignments of identical sequences in grey. (*A*) Wild types. (*B*) Variants. Sequence variation in both sequence sets is conferred by large tandem or direct repeats.

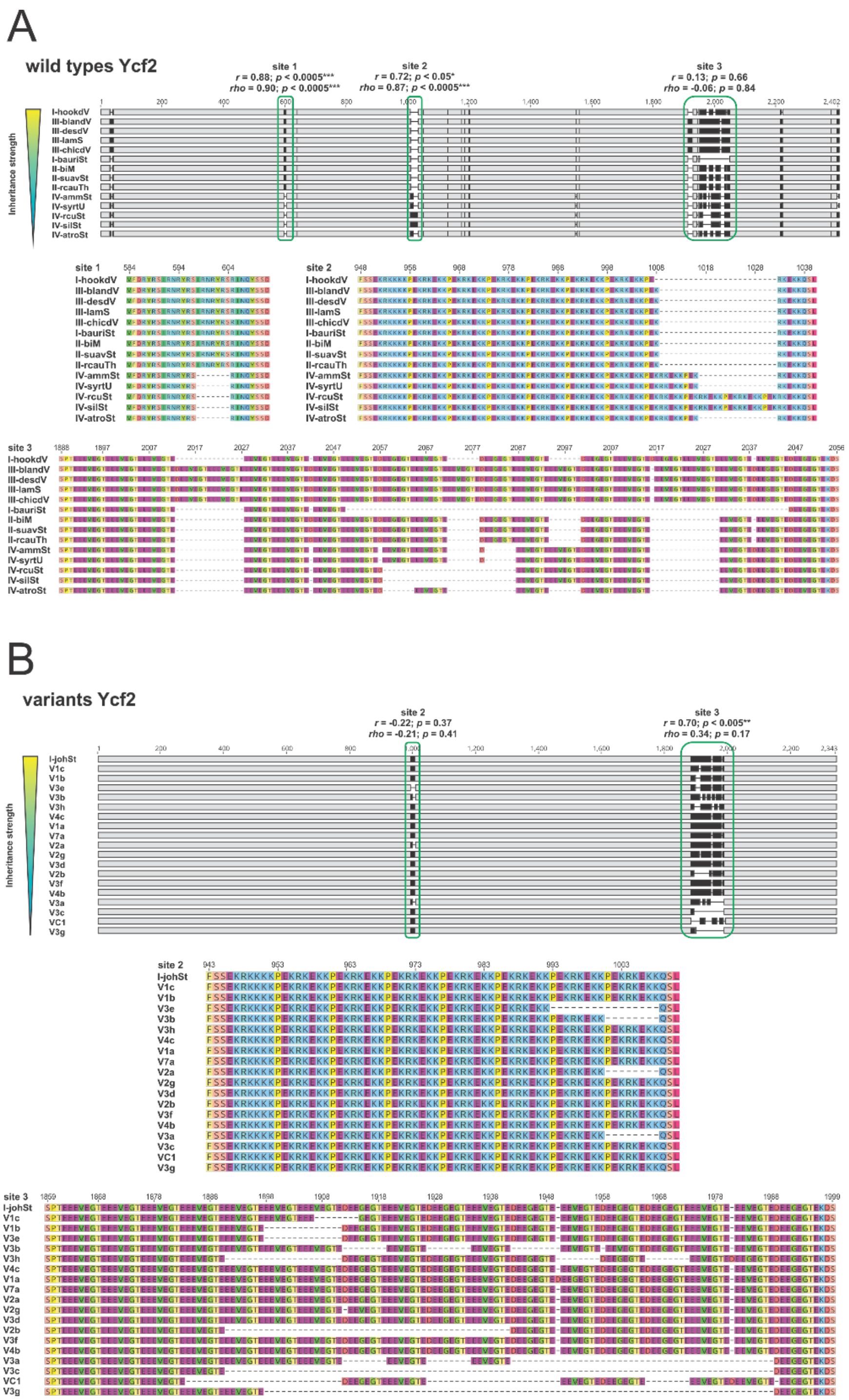


**Fig. S3.** Amino acid sequence of the Ycf2 protein and correlation to inheritance strength (Pearson's/Spearman's correlation to *k*-means classes for the wild types and to I-chi crosses for the variants). Individual sequences are sorted according to their competitive ability. Polymorphic regions are indicated in in black, alignments of identical sequences in grey. (*A*) Wild types. (*B*) Variants. Sequence variation in both sequence sets is conferred by large tandem or direct repeats. Note that sites 1 and 2, but not site 3 correlate with inheritance strengths in the wild types, whereas multiple deletions in site 3 are associated with the weaker inheritance phenotype of the variants.

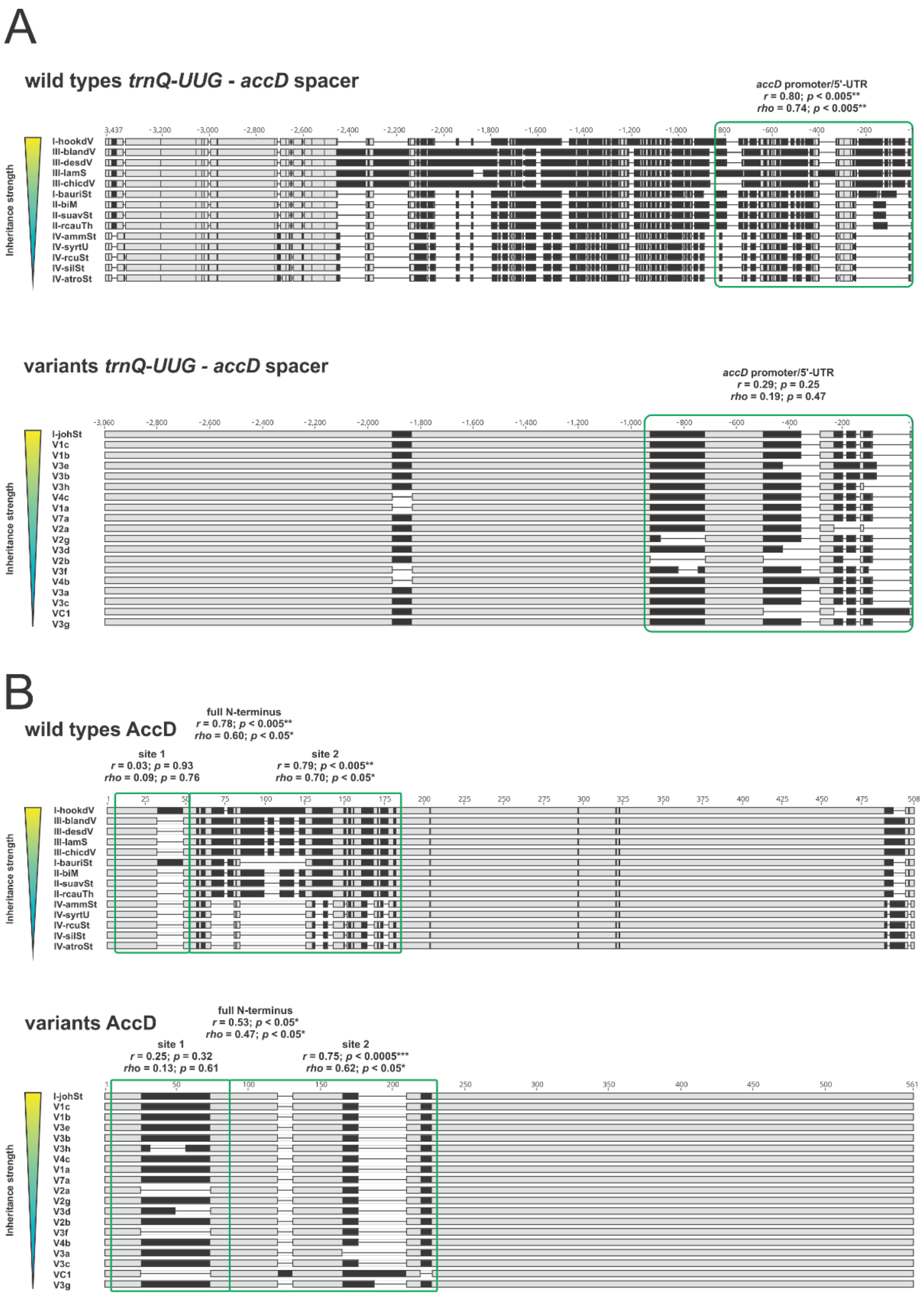


**Fig. S4.** Pearson's/Spearman's correlation to *k*-means classes (wild type) and I-chi (variants) at selected sites of the *accD* gene. Individual sequences are sorted according to their inheritance strength. Polymorphic regions are indicated in in black, alignments of identical sequences in grey. (*A*) *trnQ-UUG* - *accD* spacer (*accD* promoter/5'-UTR). (*B*) *accD* coding region. For better presentability, the protein alignment is shown. For details see Main Text and SI Text.

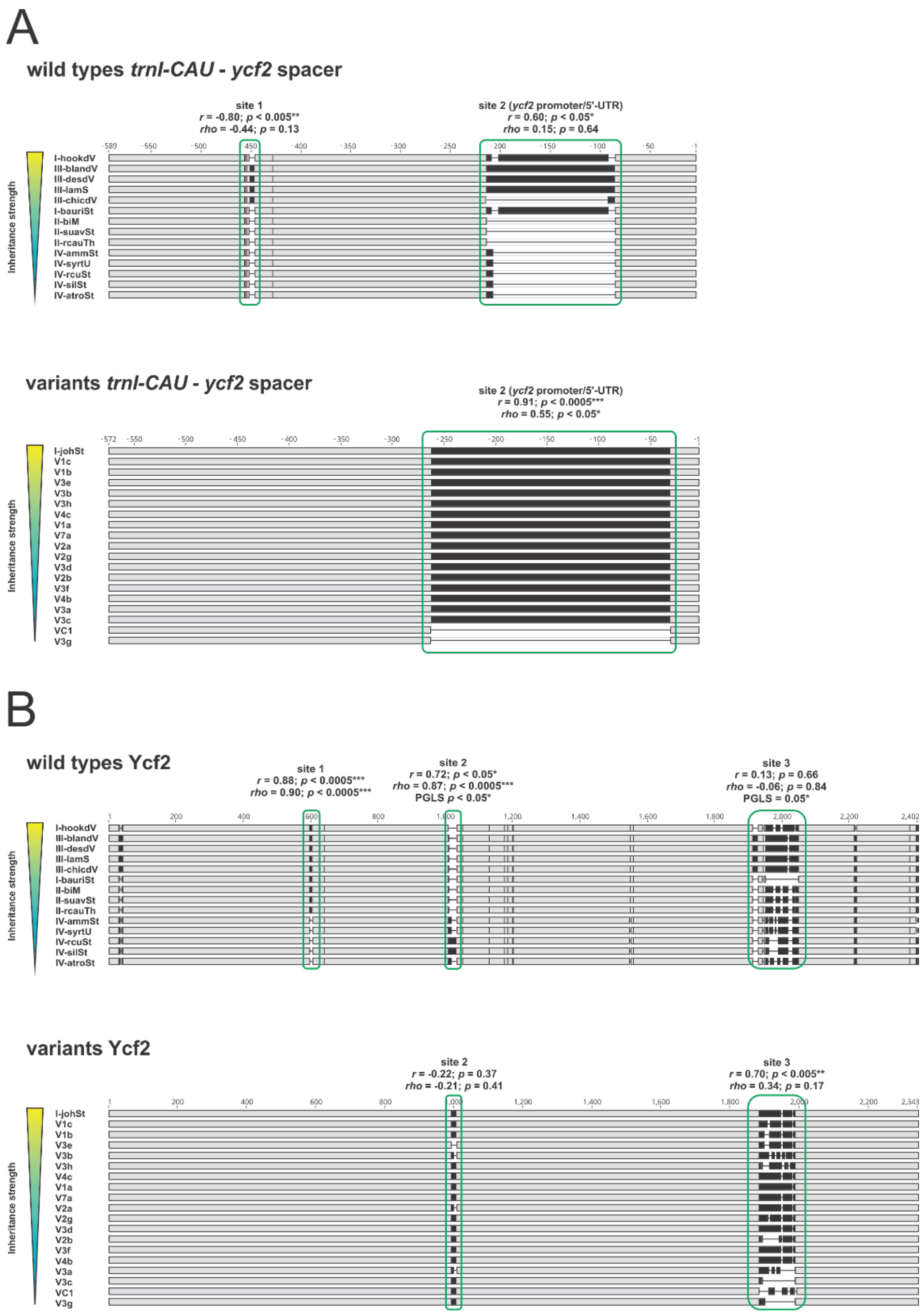


**Fig. S5.** Pearson's/Spearman's correlation to *k*-means classes (wild type) and I-chi (variants) at selected sites of the *ycf2* gene. Individual sequences are sorted according to their inheritance strength. Polymorphic regions are indicated in in black, alignments of identical sequences in grey. (*A*) *trnI-CAU* - *ycf2* spacer (*ycf2* promoter/5'-UTR). (*B*) *ycf2* coding region. Note that sites 1 and 2, but not site 3 correlate with inheritance strength in the wild types, whereas site 3 is associated with the inheritance phenotype of the variants. For better presentability, the protein alignment is shown. For details see Main Text and SI Text.

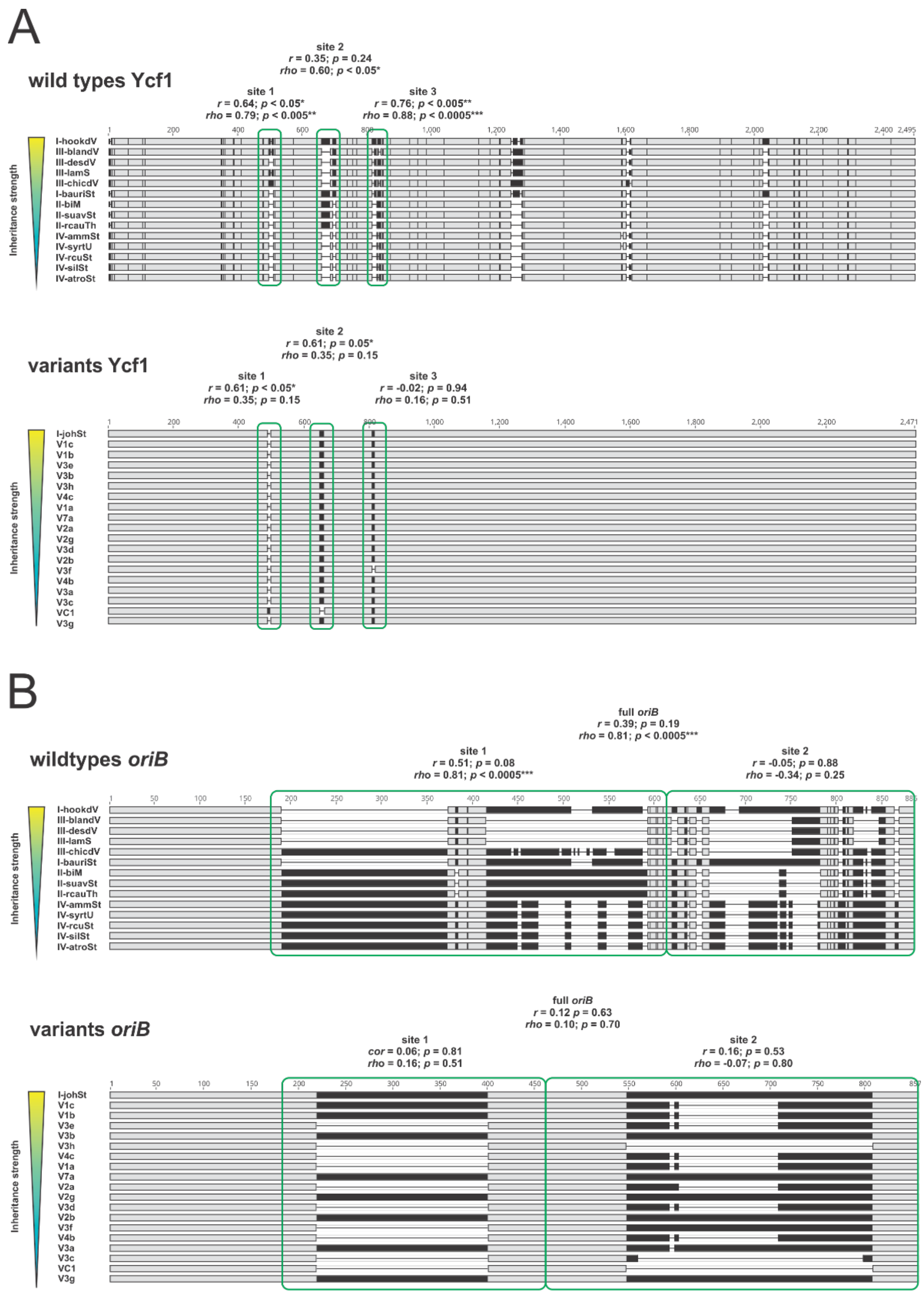

**Fig. S6.** Pearson’s/Spearman’s correlation to *k*-means classes (wild type) and I-chi (variants) at selected sites of the *ycf1* coding region and *oriB*. Individual sequences are sorted according to their inheritance strength. Polymorphic regions are indicated in in black, alignments of identical sequences in grey. (*A*) *ycf1* coding region. For better presentability, the protein alignment is shown. (*B*) *oriB*. For details see Main Text and SI Text.

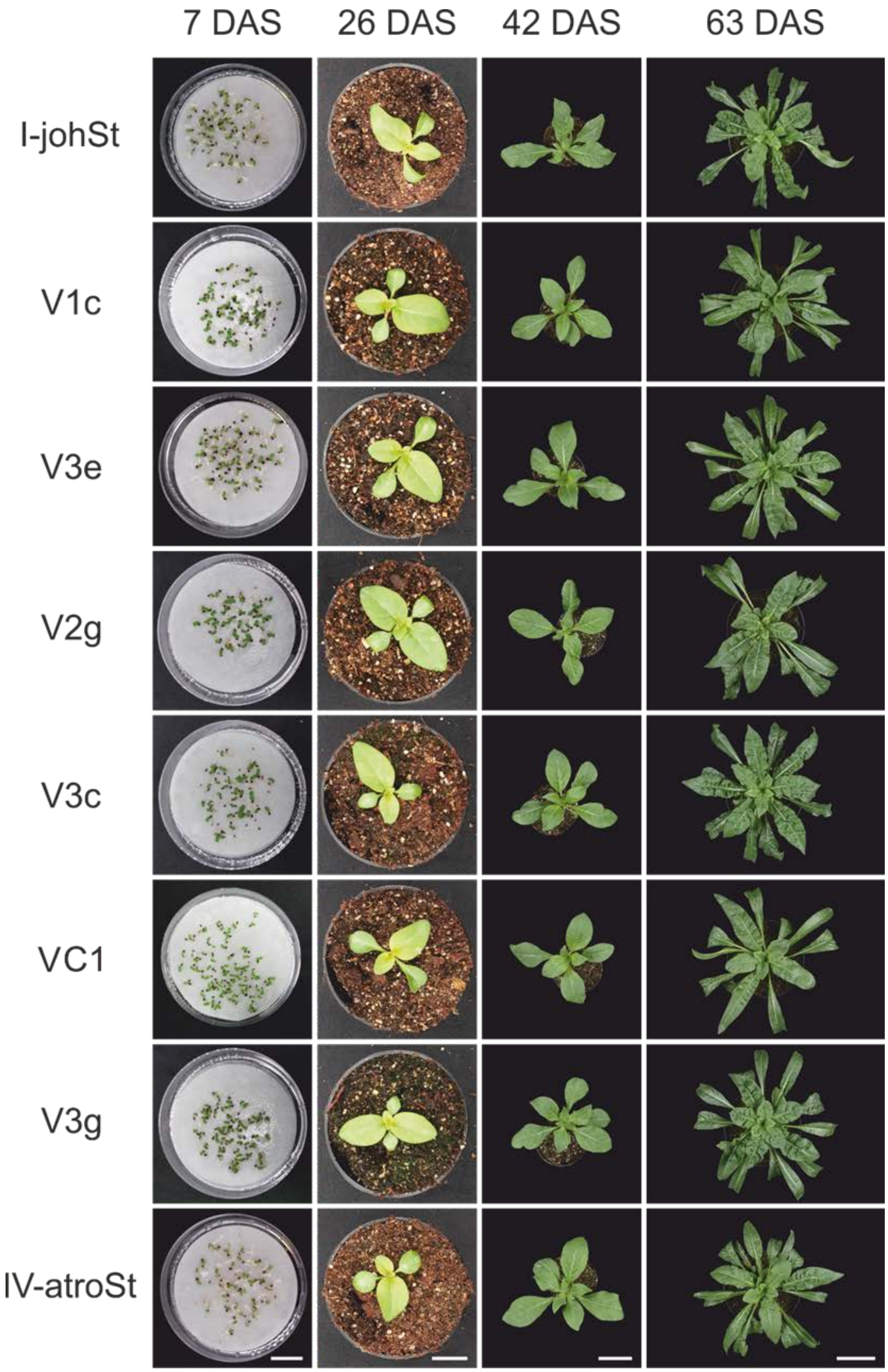


**Fig. S7.** Developmental series of johansen Standard plants with wild type or variant chloroplast genomes of different inheritance strength. Columns from left to right. First column: seedlings 4 days after geminating or 7 days after sowing (DAS); scale bar = 2 cm. Second column: plantlets 26 DAS; scale bar = 2 cm. Third column: end of early rosette stage, 42 DAS; scale bar = 4 cm. Fourth column: mature rosettes, 63 DAS; scale bar = 10 cm.

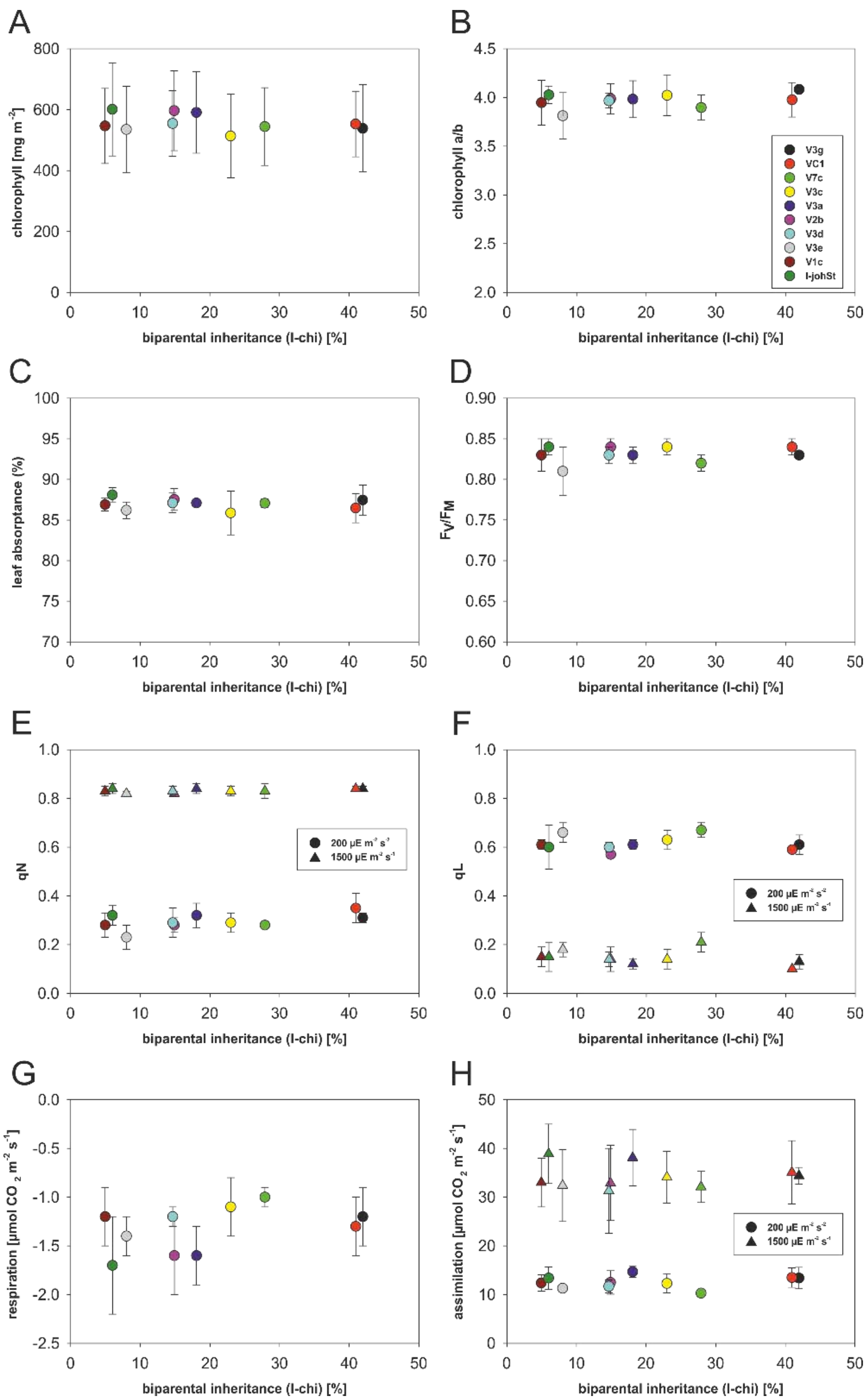


**Fig. S8.** Photosynthetic parameters of fully expanded leaves from 7-8 week-old I-johSt plants and several plastome I variants. (*A*) Chlorophyll content. (*B*) Chlorophyll a/b ratio. (*C*) Leaf absorptance. (*D*) Fv/Fm. (*E*) qN under low light intensity at 200 µE $m^{-2}$ $s^{-1}$ and saturating light intensity at 1,500 µE $m^{-2}$ $s^{-1}$. (*F*) qL under low light intensity 200 µE $m^{-2}$ $s^{-1}$ and saturating light intensity at 1,500 µE $m^{-2}$ $s^{-1}$. (*G*) Respiration measured after 30 min of dark adaptation. (*H*) $CO_2$ assimilation under low light intensity at 200 µE $m^{-2}$ $s^{-1}$ and saturation light intensity of 1500 µE $m^{-2}$ $s^{-1}$. Values are plotted against inheritance strength (biparental inheritance I-chi [%], for details see Fig. 2 and SI Text).

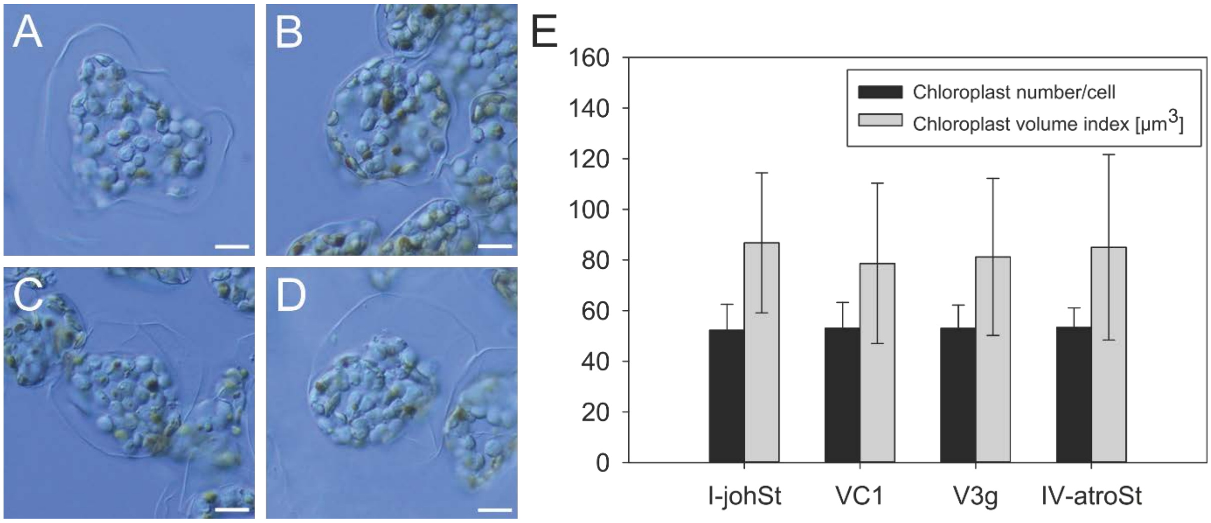


**Fig. S9.** Chloroplast size and number per cell of johansen Standard lines with wild type or variant chloroplasts of different inheritance strength. Representative cells of (*A*) I-johSt (strong wild type). (*B*) VC1 and (*C*) V3g (weak variants derived from I-johSt), and (*D*) IV-atroSt (weak wild type). (*E*) Comparison between the lines. Note lack of statistically significant differences (Tables S15 and S16; SI Text for details). Scale bar = 10 μm

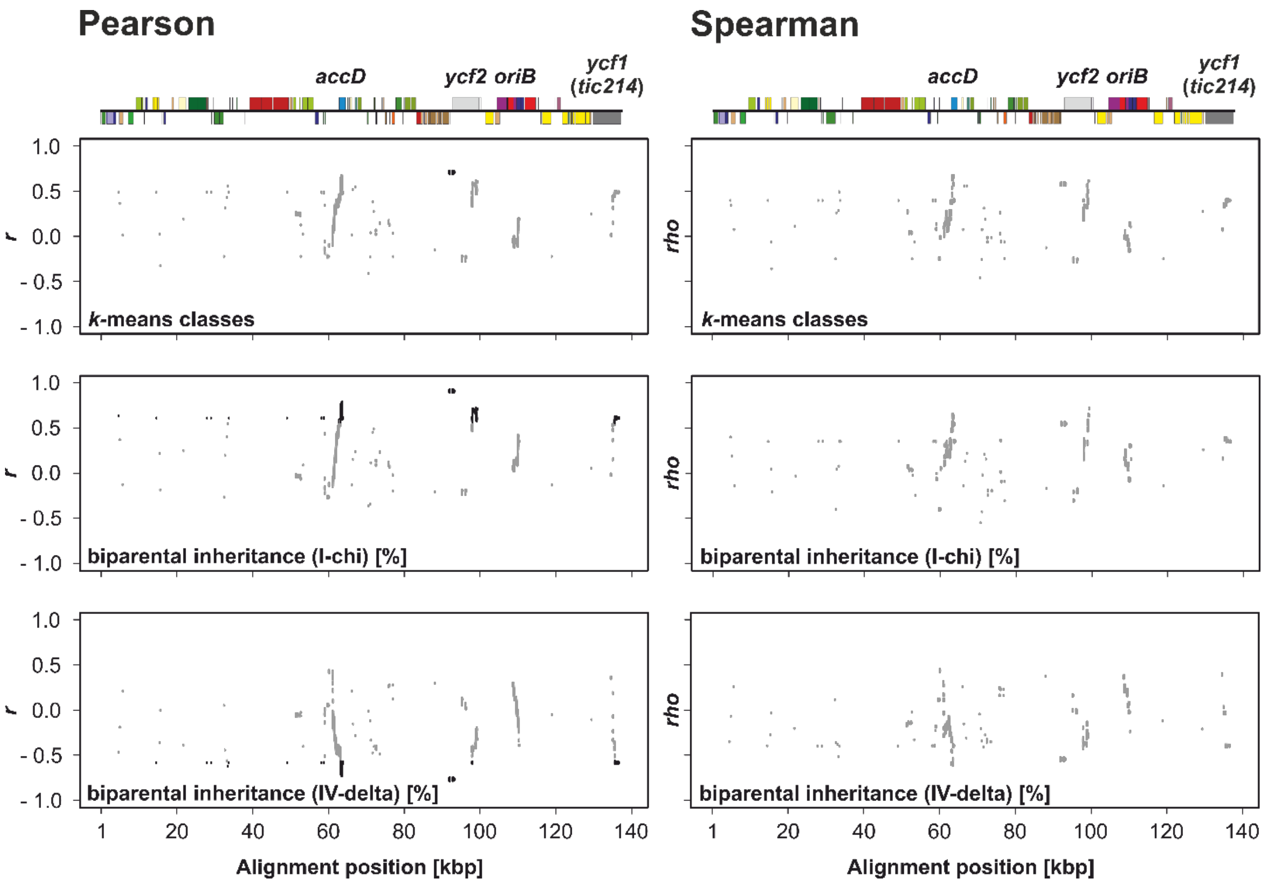


**Fig. S10.** Correlation mapping in the plastome I variants. Pearson's and Spearman's correlation to the *k*-means classes and the I-chi/IV-delta crossing data. Relevant genes or loci with significant correlation are designated in the linear plastome maps above. Non-significant correlations after *p*-value adjustment ($p > 0.05$) are displayed in grey. Note that regions positively correlated in the I-chi cross are negatively correlated in the reciprocal IV-delta cross. Also note that PGLS analysis is not applicable to a set of independent mutants. For details see SI Text.

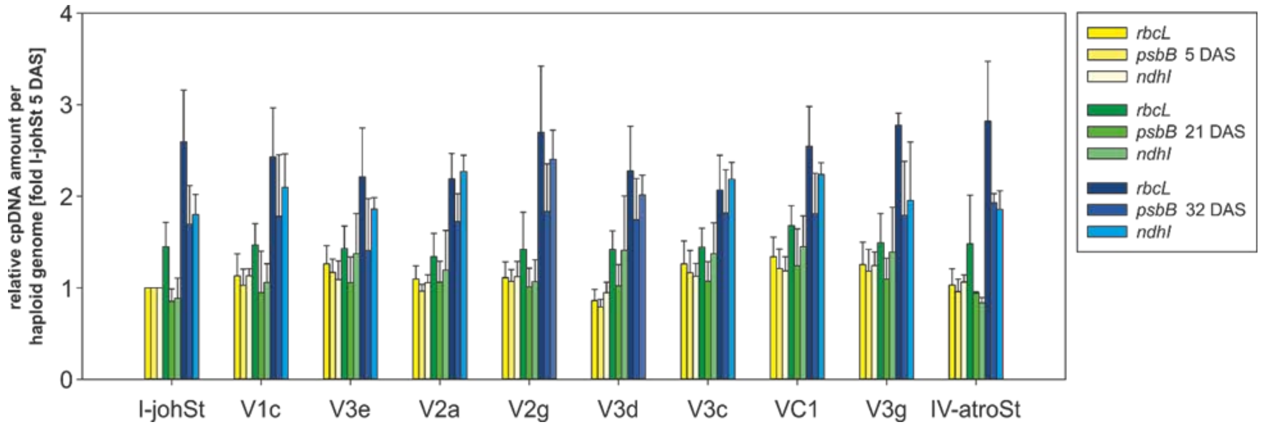


**Fig. S11.** Relative ptDNA content of the *rbcL*, *psbB* and *ndhI* loci at developmental stages 5 DAS (cotyledons), 21 DAS (cotyledons and first and second true leave) and 32 DAS (second true leaf) as judged from quantitative real-time PCR. Data for plastid markers were normalized to the mean amounts of three nuclear markers (M02, M19, and pgiC) and expressed as relative values [fold I-johSt 5 DAS]. For details see SI Materials and Methods. No statistical significant differences between the lines at a given developmental stage are observed (also see SI Text).

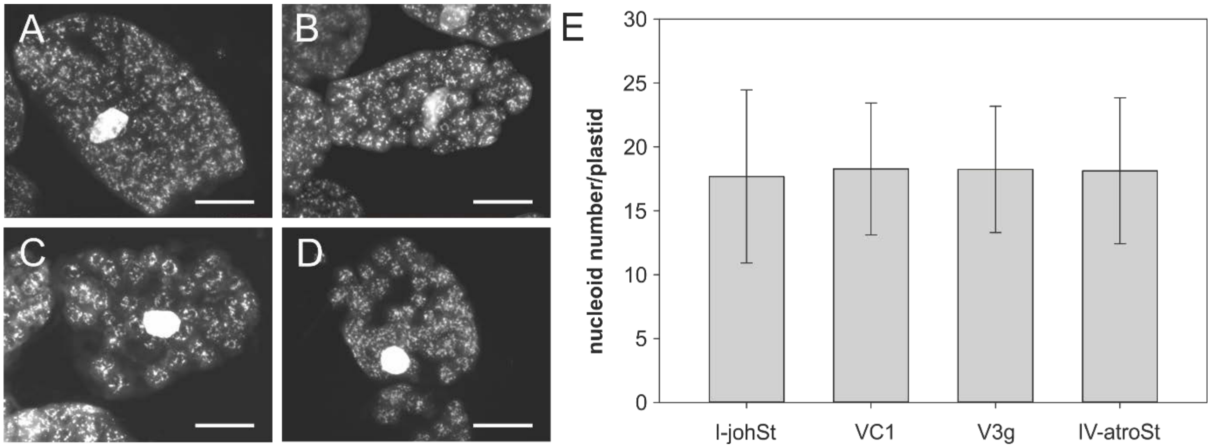


**Fig. S12.** ptDNA nucleoids visualized by DAPI and summary of nucleoid counts in chloroplasts of different inheritance strengths. Representative cells of (*A*) I-johSt (strong wild type). (*B*) VC1 and (*C*) V3g (weak variants derived from I-johSt), and (*D*) IV-atroSt (weak wild type). (*E*) Comparison of nucleoid number/plastid in the lines. Note lack of statistically significant differences (cf. Fig. S13, Table S14, SI Materials and Methods, and SI Text). Scale bar = 10 µm

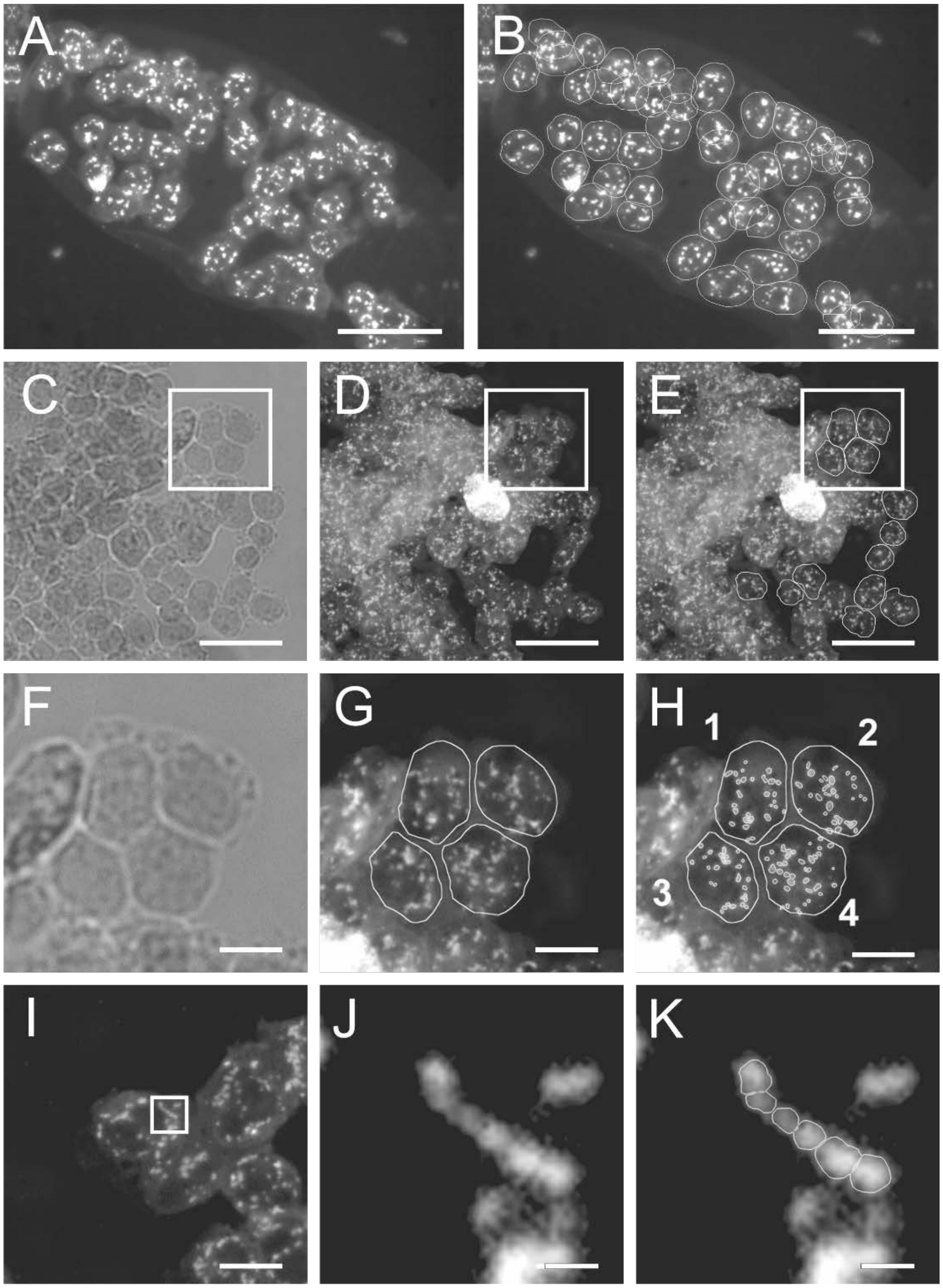


**Fig. S13.** Representative examples of nucleoid counting in DAPI-stained chloroplasts. (*A*) Original image of VC1 cells. (*B*) Manual delimitation of single chloroplasts in (*A*). Only chloroplasts with non-overlapping nucleoids were analysed. (*C*-*K*) Nucleoid counting in IV-atroSt chloroplasts. (*C*-*E*) Regions from which cell fragments in (*F*-*H*) derive. (*F*-*H*) Chloroplast delimitation and counting. (*F*) Bright field. (*G*) Delimitation of chloroplast. (*H*) Counting. Chloroplast 1: 26 nucleoids, chloroplast 2: 28 nucleoids, chloroplast 3: 27 nucleoids, and chloroplast 4: 39 nucleoids. (*I*-*K*) Detangling of nucleoids aggregated in clumps and/or threads. (*I*), Region from which the nucleoid thread in j and k derives from. (*J*) Nucleoid thread. (*K*) Delimitation and counting of single nucleoids in the thread. Scale bar = 10 µm in (*A*-*E*), 2.5 µm in (*F*-*H*) and 0.25 µm in (*I*-*K*).

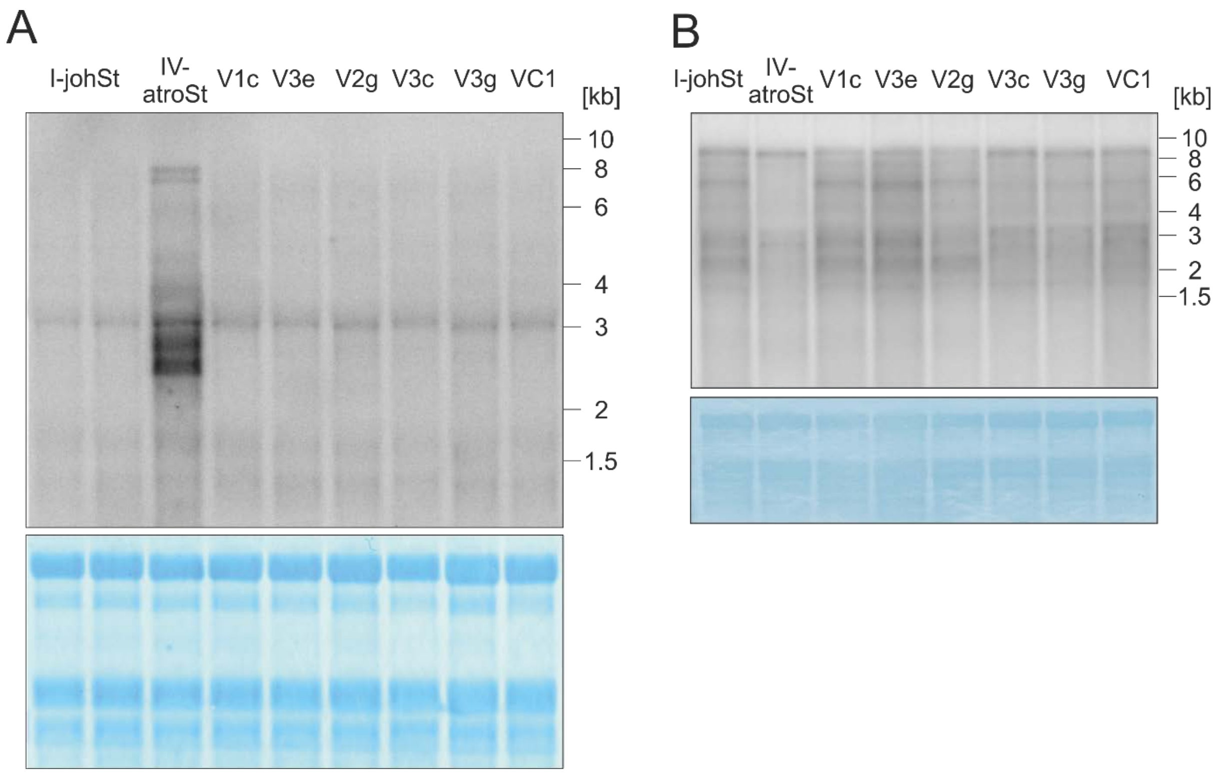


**Fig. S14.** Transcript accumulation of the *accD* and *ycf2* mRNA in lines of different inheritance strengths. (*A*) *accD*. Note the strong over-expression in the weak wild type IV-atroSt, whereas the plastome I variants do not differ from their wild type I-johSt. (*B*) *ycf2*; mature transcript (9 kb) and degradation/processing products. The latter differ between the strong and the weak wild types (I-johSt vs. IV-atroSt). The patterns are similar in the stronger (V1C, V3e and V2g) and weaker (V3c, V3g, and VC1) materials, correlating with the presence of mutations in site 3 of *ycf2* (Fig. S2).

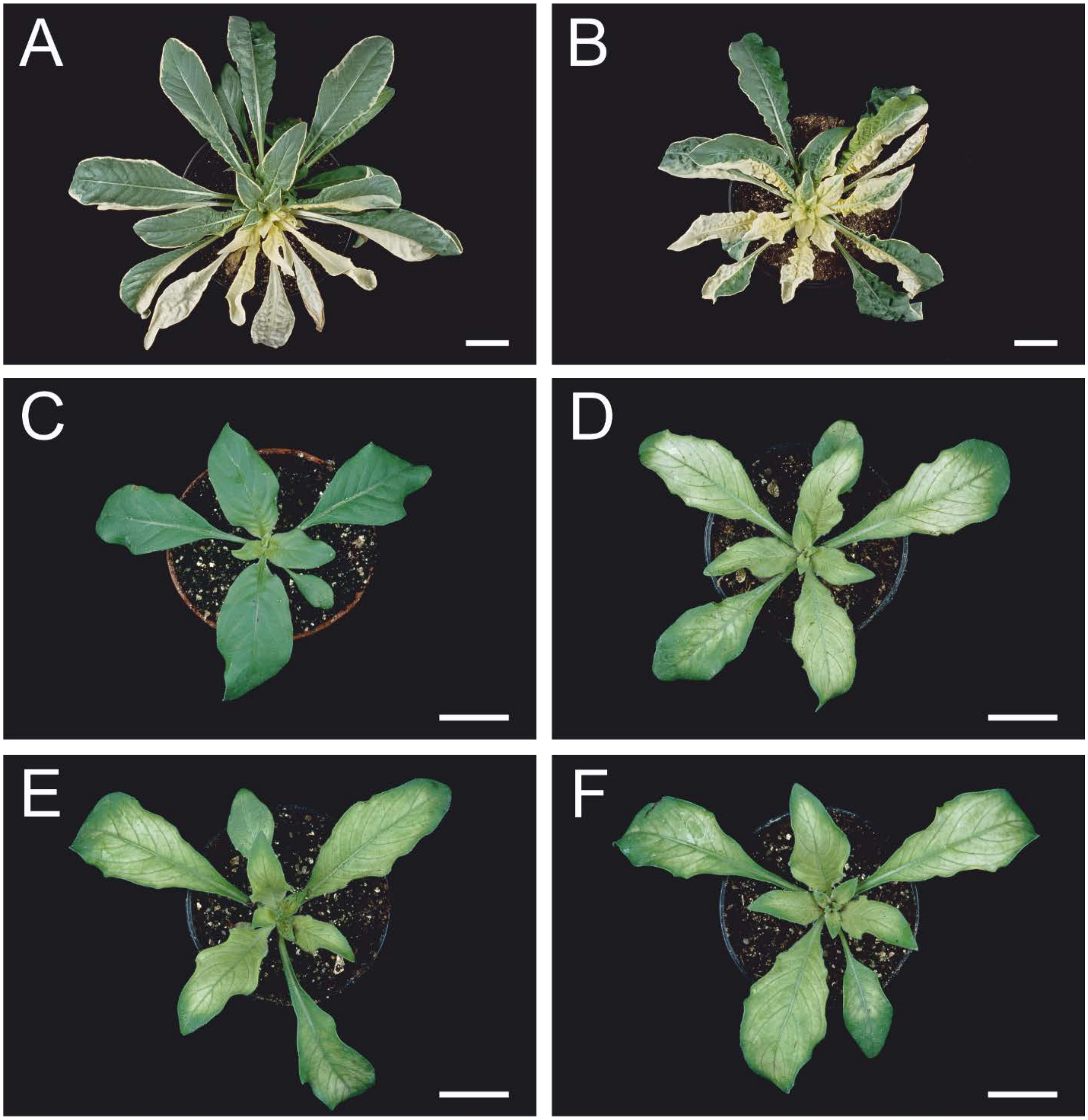


**Fig. S15.** Plants with chloroplast genotypes resulting in bleaching used in this work. (*A*,*B*) Variegated lines combining wild type (green) and mutated (white) chloroplasts in one individual. (*C*-*F*) Homoplasmic material with only one chloroplast type. (*A*) Plastome mutant I-chi (white) and wild type IV-atroSt (green). (*B*) Plastome mutant IV-delta (white) and wild type I-johSt (green). (*C*) Chloroplast genome I-johSt in its native nuclear background. (*D*-*F*) *Virescent* phenotype of III-lamS and derived variants in the background of johansen Standard. (*D*) Wild type genotype III-lamS, (*E*) III-V1, and (*F*) III-V2; two independent very weak variants derived from the strong plastome III-lamS. (*A*,*B*) Mature rosette. (*C*-*F*) Early rosette stage. Scale bar = 5 cm in (*A*,*B*) and 2 cm in (*C*-*F*)

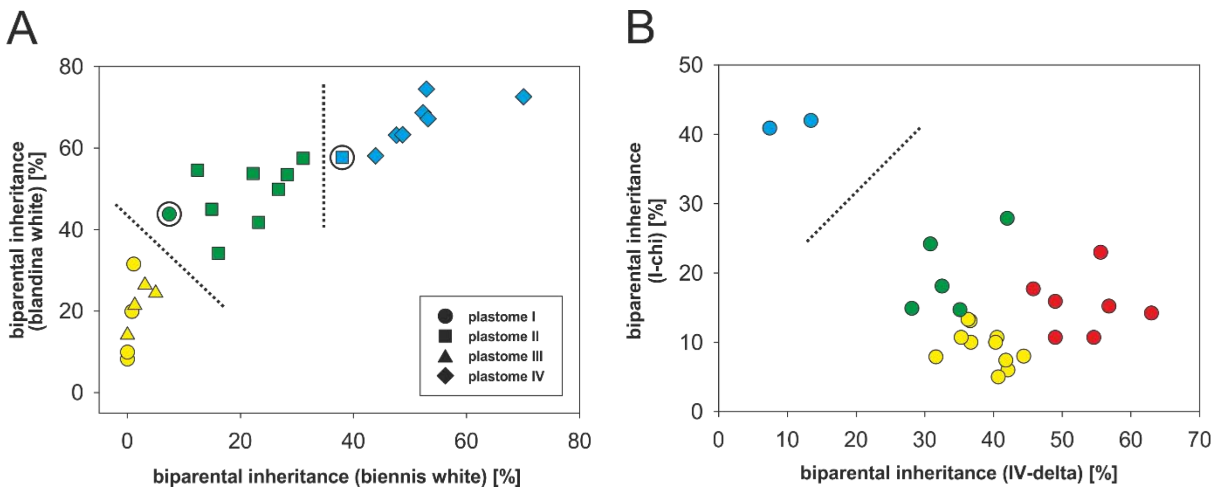


**Fig. S16.** Inheritance strength of the wild type and variant plastomes based on crossing data with white tester chloroplasts. The two crossing series, “biennis/blandina white” (wild types) and I-chi/IV-delta (variants), are plotted against each other. (*A*) Wild type plastomes of Table S11: *k*-means clustering with three centers identified by the pamk function (dotted lines) confirms the classes of Schötz (strong, intermediate and weak), indicated as yellow, green, and blue and comprised of plastome I and III, plastome II, and plastome IV, respectively. The two exceptions I-bauriSt and II-corSt are circled. (*B*) Plastome I variants of Table S8: *k*-means clustering with two centers (again obtained from the pamk function) clearly separates the dataset (dotted line), whereas finer clustering into four groups results in a new class (red) not present in the wild types (strong = yellow, strong to intermediate = red, intermediate = green, weak = blue). For details see SI Text.

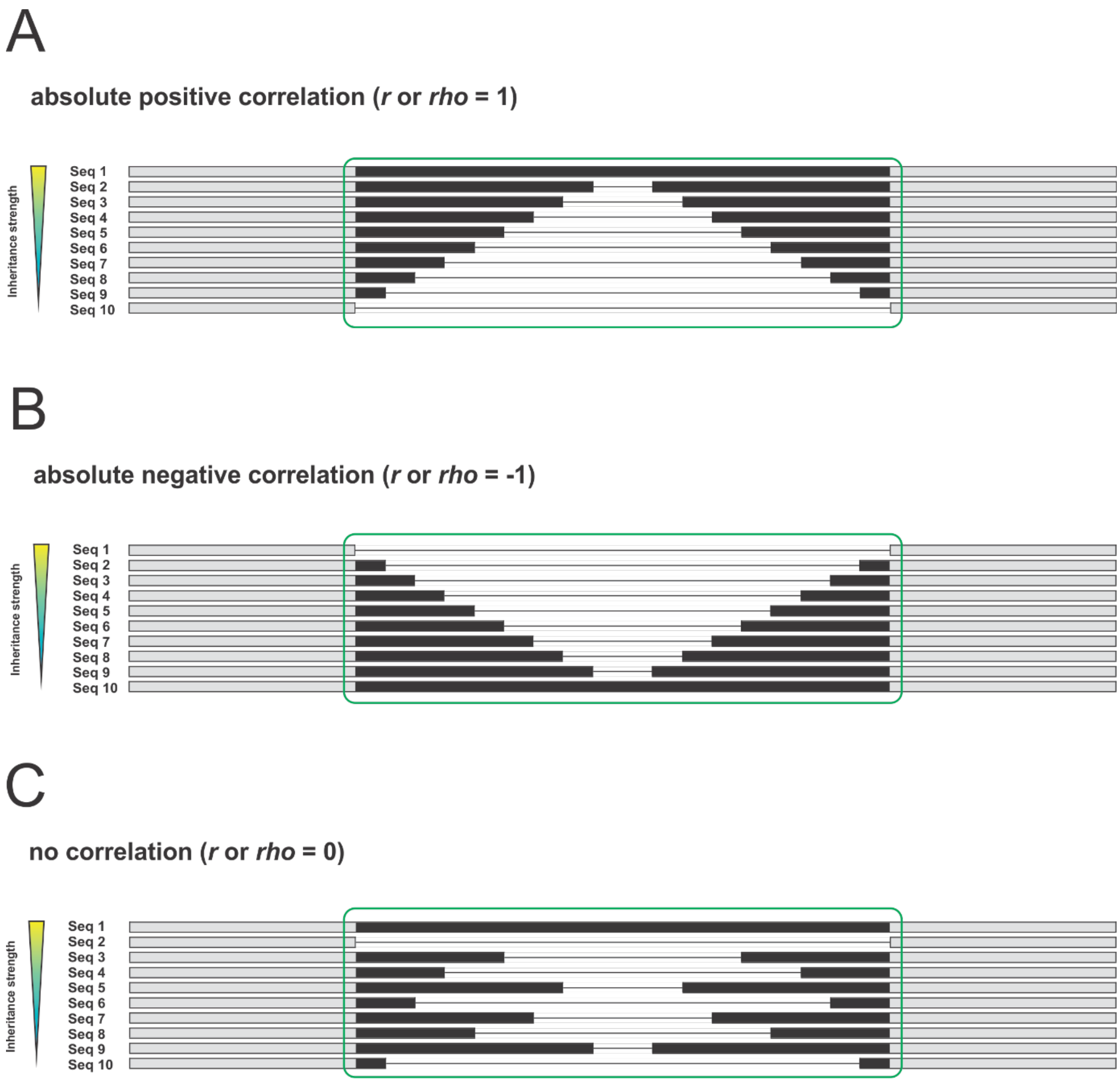


**Fig. S17.** Principal of the correlation mapping approach. Individual sequences are sorted according to their inheritance strength. Polymorphic regions are indicated in in black, alignments of identical sequences in grey. Sequence alignment window with absolute positive (*A*) absolute negative (*B*) and no correlation (*C*) to inheritance strengths. For details see Materials and Methods and SI Text.

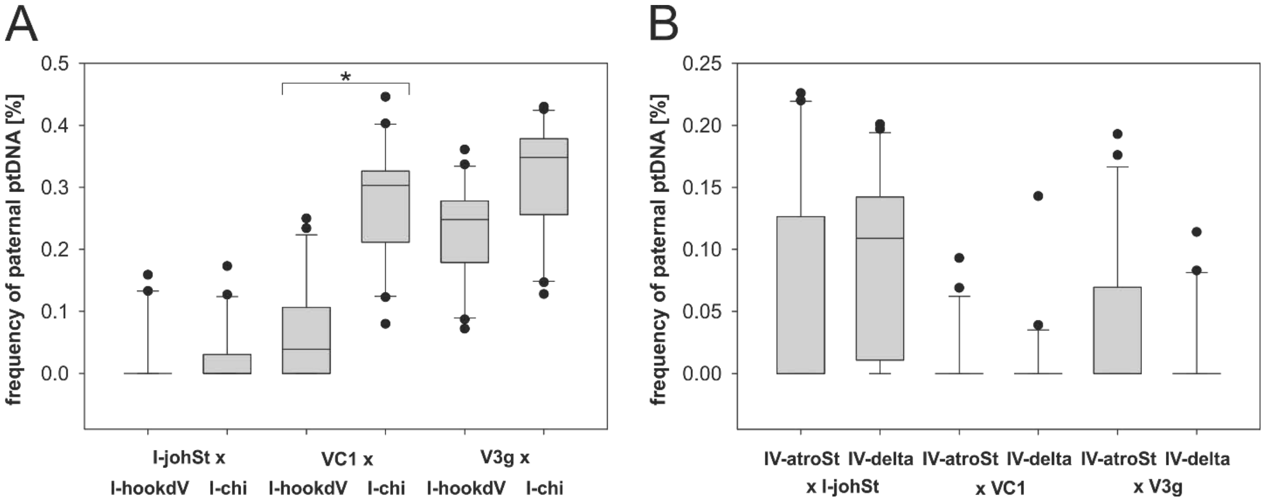


**Fig. S18.** Transmission efficiencies of wild type and derived bleached mutant chloroplasts in crosses to I-johSt, VC1 and V3g. (*A*) Wild type/mutant pair (I-hookdV/I-chi) as pollen parent. (*B*) Wild type/mutant pair (IV-atroSt/IV-delta) as seed parent. Box-plots represent frequencies of the paternal ptDNA [%] measured by the MassARRAY® system. Significance of difference (VC1 x I-hookdV vs. VC1 x I-chi) was tested with Kruskal-Wallis one-way ANOVA on ranks (* $p < 0.05$).

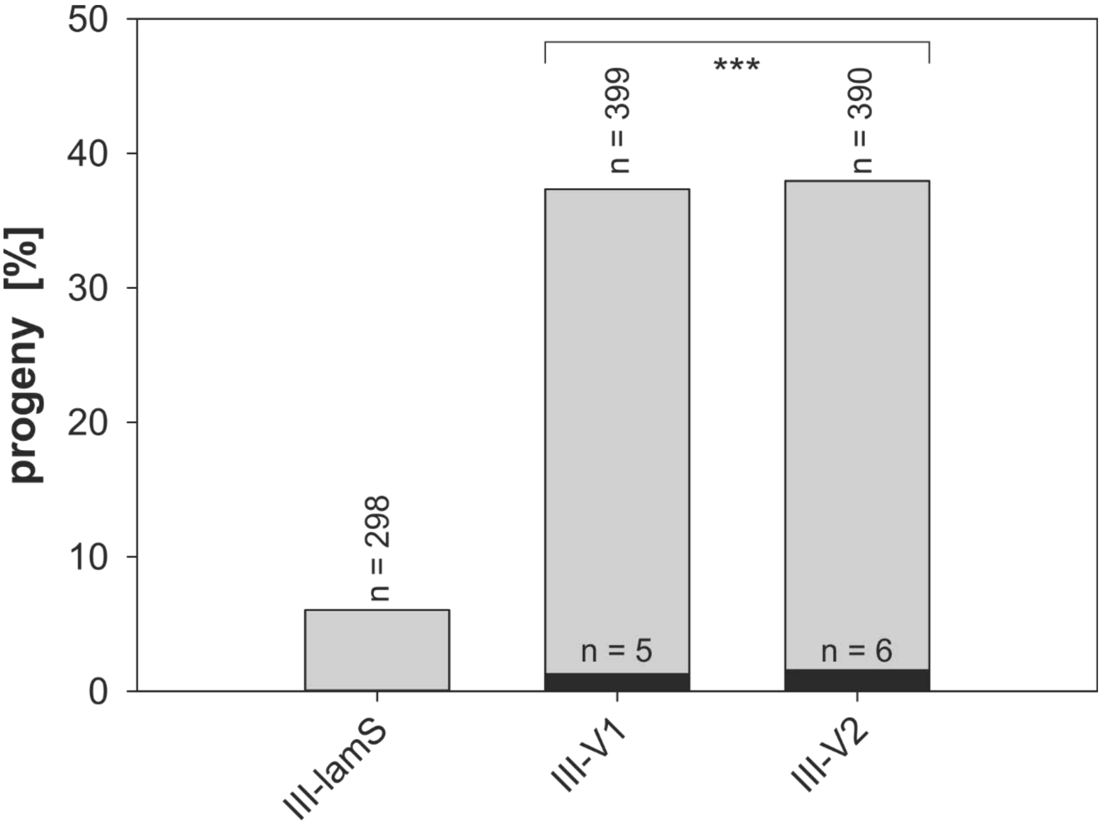


**Fig. S19.** Transmission efficiencies of wild type plastome III-lamS and its variants III-V1 and III-V2 in crosses to I-johSt as pollen donor. Depicted in grey is the average percentage of variegated progeny (= biparental inheritance [%]) obtained from two seasons (2015 and 2016). Significance of differences between variegated progeny of III-lamS and the variants was calculated by Fisher`s exact test (*** $p < 0.0001$); n black bars = total number of seedlings homoplasmic for the paternal chloroplast, n grey bars = total number of seedlings counted in the experiment.

**Table S1.** Summary of the mutations found in the green variants of I-johSt[1)]

| Variant | Inheritance class[2)] | Alignment position[3)] | Type | size [bp] | Allele in WT I-johSt | Allele in mutant I-johSt variant | Gene/Region | Type/Impact[4)] |
|---|---|---|---|---|---|---|---|---|
| V1a | 2 | 60935 - 61008 | DEL | 74 | CATAG…GTTGA | - | *trnQ-UUG* - *accD* spacer | intergenic, larger deletion |
| V1a | 2 | 67064 | INS | 1 | - | A | *ycf4* - *cemA* spacer | intergenic, oligo(N) stretch |
| V1a | 2 | 98590 - 98592 | INS | 3 | - | GAT | *ycf2* | 1 AS insertion at site 3 of Ycf2 |
| V1a | 2 | 98600 | SNP | 1 | T | G | *ycf2* | codon 1952, V:GTA to G:GGA |
| V1a | 2 | 109065 - 109246 | DEL | 182 | TCCCC…GCATG | - | *oriB* | large deletion in *oriB* |
| V1a | 2 | 109439 - 109443 | DEL | 5 | ACATA | - | *oriB* | small deletion in *oriB* |
| V1a | 2 | 109449 - 109553 | DEL | 105 | GGATG…GGAGC | - | *oriB* | large deletion in *oriB* |
| V1b | 1 | 54036 | DEL | 1 | A | - | *atpF* intron | intron, oligo(N) stretch |
| V1b | 1 | 71172 | INS | 1 | - | T | *psbE* - *petL* spacer | intergenic, oligo(N) stretch |
| V1b | 1 | 71718 | INS | 1 | - | A | *psbE* - *petL* spacer | intergenic, oligo(N) stretch |
| V1b | 1 | 73173 | DEL | 1 | G | - | *trnP-UGG* - *psaJ* spacer | intergenic, single bp deletion |
| V1b | 1 | 73177 - 73182 | DEL | 6 | TAAGAA | - | *trnP-UGG* - *psaJ* spacer | intergenic, small deletion |
| V1b | 1 | 76536 | SNP | 1 | C | G | *rps12* - *clpP* spacer | intergenic, SNP |
| V1b | 1 | 98434 - 98475 | DEL | 42 | GAGGA…CAGAA | - | *ycf2* | 14 AS deletion at site 3 of Ycf2 |
| V1b | 1 | 109439 - 109443 | DEL | 5 | ACATA | - | *oriB* | small deletion in *oriB* |
| V1b | 1 | 109449 - 109553 | DEL | 105 | GGATG…GGAGC | - | *oriB* | large deletion in *oriB* |
| V1c | 1 | 22673 | INS | 1 | - | T | *ycf3* - *psaA* spacer | intergenic, oligo(N) stretch |
| V1c | 1 | 33274 | INS | 1 | - | A | *psbD* - *trnT-GGU* | intergenic, oligo(N) stretch |
| V1c | 1 | 54036 | DEL | 1 | A | - | *atpF* intron | intron, oligo(N) stretch |
| V1c | 1 | 59739 - 59764 | INS | 26 | - | CTAGT…TAGAA | *psbK* - *trnQ-UUG* | intergenic, deletion |
| V1c | 1 | 71172 | INS | 1 | - | T | *psbE* - *petL* spacer | intergenic, oligo(N) stretch |
| V1c | 1 | 71718 | INS | 1 | - | A | *psbE* - *petL* spacer | intergenic, oligo(N) stretch |
| V1c | 1 | 73173 | DEL | 1 | G | - | *trnP-UGG* - *psaJ* spacer | intergenic, single bp deletion |
| V1c | 1 | 73177 - 73182 | DEL | 6 | TAAGAA | - | *trnP-UGG* - *psaJ* spacer | intergenic, small deletion |
| V1c | 1 | 76536 | SNP | 1 | C | G | *rps12* - *clpP* spacer | intergenic, SNP |
| V1c | 1 | 88454 | DEL | 1 | A | - | *rpl16* - *rps3* | intergenic, oligo(N) stretch |
| V1c | 1 | 98462 - 98485 | DEL | 24 | TAGAA…GGAAG | - | *ycf2* | 8 AS deletion at site 3 of Ycf2 |
| V1c | 1 | 109439 - 109443 | DEL | 5 | ACATA | - | *oriB* | small deletion in *oriB* |
| V1c | 1 | 109449 - 109553 | DEL | 105 | GGATG…GGAGC | - | *oriB* | large deletion in *oriB* |
| V2a | 1 | 52794 | INS | 1 | - | T | *atpH* - *atpF* spacer | intergenic, oligo(N) stretch |
| V2a | 1 | 62054 | SNP | 1 | T | G | *trnQ-UUG* - *accD* spacer | SNP in *accD* promotor/5'-UTR |
| V2a | 1 | 62610 - 62690 | DEL | 66 | GATCC…CTTTA | - | *trnQ-UUG* - *accD* spacer | larger deletion in *accD* promotor/5'-UTR |
| V2a | 1 | 62724 - 62753 | DEL | 30 | CCTTT…CTTTA | - | *trnQ-UUG* - *accD* spacer | deletion in accD promotor/5'-UTR |
| V2a | 1 | 62981 - 63124 | DEL | 144 | TAAGG…GATAA | - | *accD* | 44 AS deletion in AccD |
| V2a | 1 | 95744 - 95767 | DEL | 24 | CCGAA…AAAAC | - | *ycf2* | 8 AS deletion at site 2 of Ycf2 |
| V2a | 1 | 109065 - 109246 | DEL | 182 | TCCCC…GCATG | - | *oriB* | large deletion in *oriB* |
| V2a | 1 | 109449 - 109553 | DEL | 105 | GGATG…GGAGC | - | *oriB* | large deletion in *oriB* |

**Table S1** (continued)

| Variant | Inheritance class[2)] | Alignment position[3)] | Type | size [bp] | Allele in WT I-johSt | Allele in mutant I-johSt variant | Gene/Region | Type/Impact[4)] |
|---|---|---|---|---|---|---|---|---|
| V2b | 3 | 33524 | INS | 1 | - | T | *psbD - trnT-GGU* | intergenic, oligo(N) stretch |
| V2b | 3 | 61914 - 62120 | DEL | 207 | TCGGC…ATGCC | - | *trnQ-UUG - accD* spacer | larger deletion in *accD* promotor/5'-UTR |
| V2b | 3 | 62341 - 62484 | DEL | 144 | AAGAT…AGATC | - | *trnQ-UUG - accD* spacer | larger deletion in *accD* promotor/5'-UTR |
| V2b | 3 | 62658 - 62690 | DEL | 33 | GATCC…CTTTA | - | *trnQ-UUG - accD* spacer | deletion in *accD* promotor/5'-UTR |
| V2b | 3 | 67063 | DEL | 1 | A | - | *ycf4 - cemA* spacer | intergenic, oligo(N) stretch |
| V2b | 3 | 71576 | DEL | 1 | T | - | *psbE - petL* spacer | intergenic, oligo(N) stretch |
| V2b | 3 | 98413 - 98565 | DEL | 153 | GAGGA…CAGAA | - | *ycf2* | 51 AS deletion at site 3 Ycf2 |
| V2g | 1 | 61950 - 61120 | DEL | 171 | TCGGC…ATGCC | - | *trnQ-UUG - accD* spacer | large deletion in *accD* promotor/5'-UTR |
| V2g | 1 | 98486 | SNP | 1 | G | T | *ycf2* | codon 1915, G:GGA to V:GTA |
| V2g | 1 | 98476 - 98478 | DEL | 3 | GAT | - | *ycf2* | 1 AS deletion at site 3 Ycf2 |
| V3a | 3 | 63401 - 63433 | DEL | 33 | TAAGC…GATAA | - | *accD* | 11 AS deletion in AccD |
| V3a | 3 | 95744 - 95767 | DEL | 24 | CCGAA…AAAAC | - | *ycf2* | 8 AS deletion at site 2 of Ycf2 |
| V3a | 3 | 98476 - 98499 | DEL | 24 | GATGA…CAGAA | - | *ycf2* | 8 AS deletion at site 3 Ycf2 |
| V3a | 3 | 98521 - 98544 | DEL | 24 | GATGA…CAGAA | - | *ycf2* | 8 AS deletion at site 3 Ycf2 |
| V3a | 3 | 98566 - 98706 | DEL | 135 | GATGA…CAGAA | - | *ycf2* | 45 AS deletion at site 3 Ycf2 |
| V3a | 3 | 109439 - 109443 | DEL | 5 | ACATA | - | *oriB* | small deletion in *oriB* |
| V3a | 3 | 129507 | INS | 1 | - | T | *rps15 - ycf1* spacer | intergenic, oligo(N) stretch |
| V3b | 1 | 62691 - 62708 | INS | 18 | - | GATCC…CTTTA | *trnQ-UUG - accD* spacer | insertion in *accD* promotor/5'-UTR |
| V3b | 1 | 62754 - 62768 | INS | 15 | - | CCTTT…CTTTA | *trnQ-UUG - accD* spacer | insertion in *accD* promotor/5'-UTR |
| V3b | 1 | 95744 - 95767 | DEL | 24 | CCGAA…AAAAC | - | *ycf2* | 8 AS deletion at site 2 of Ycf2 |
| V3b | 1 | 98476 - 98499 | DEL | 24 | GATGA…CAGAA | - | *ycf2* | 8 AS deletion at site 3 of Ycf2 |
| V3b | 1 | 98521 - 98544 | DEL | 24 | GATGA…CAGAA | - | *ycf2* | 8 AS deletion at site 3 of Ycf2 |
| V3b | 1 | 98566 - 98589 | DEL | 24 | GATGA...CAGAA | - | *ycf2* | 8 AS deletion at site 3 of Ycf2 |
| V3b | 1 | 98614 - 98616 | DEL | 3 | GAT | - | *ycf2* | 1 AS deletion at site 3 Ycf2 |
| V3b | 1 | 98624 | SNP | 1 | G | T | *ycf2* | codon 1960, G:GGA to V:GTA |
| V3c | 2 | 71718 | INS | 1 | - | A | *psbE - petL* spacer | intergenic, oligo(N) stretch |
| V3c | 2 | 76523 | SNP | 1 | T | G | *rps12 - clpP* spacer | intergenic, SNP |
| V3c | 2 | 76536 | SNP | 1 | C | G | *rps12 - clpP* spacer | intergenic, SNP |
| V3c | 2 | 76526 - 76532 | INS | 7 | - | TTAAGAA | *rps12 - clpP* spacer | intergenic, SNP |
| V3c | 2 | 98413 - 98706 | DEL | 288 | GAGGA…CAGAA | - | *ycf2* | 96 AS deletion at site 3 Ycf2 |
| V3c | 2 | 109065 - 109246 | DEL | 182 | TCCCC…GCATG | - | *oriB* | large deletion in *oriB* |
| V3c | 2 | 109406 - 109642 | DEL | 238 | GTTGG…AGCTC | - | *oriB* | large deletion in *oriB* |

**Table S1** (continued)

| Variant | Inheritance class[2)] | Alignment position[3)] | Type | size [bp] | Allele in WT I-johSt | Allele in mutant I-johSt variant | Gene/Region | Type/Impact[4)] |
|---|---|---|---|---|---|---|---|---|
| V3d | 3 | 52315 - 52392 | DEL | 78 | AATCT…TCCAT | - | *atpI* - *atpH* spacer | intergenic, larger deletion |
| V3d | 3 | 62413 - 62484 | DEL | 72 | AAGAT…AGATC | - | *trnQ-UUG* - *accD* spacer | larger deletion in *accD* promotor/5'-UTR |
| V3d | 3 | 63053 - 63124 | DEL | 72 | TAAGG…GATAA | - | *accD* | 24 AS deletion in AccD |
| V3d | 3 | 67063 | DEL | 1 | A | - | *ycf4* - *cemA* spacer | intergenic, oligo(N) stretch |
| V3d | 3 | 76523 | SNP | 1 | T | G | *rps12* - *clpP* spacer | intergenic, SNP |
| V3d | 3 | 76536 | SNP | 1 | C | G | *rps12* - *clpP* spacer | intergenic, SNP |
| V3d | 3 | 76526 - 76532 | INS | 7 | - | TTAAGAA | *rps12* - *clpP* spacer | intergenic, small deletion |
| V3d | 3 | 109065 - 109246 | DEL | 182 | TCCCC…GCATG | - | *oriB* | large deletion in *oriB* |
| V3d | 3 | 109439 - 109443 | DEL | 5 | ACATA | - | *oriB* | small deletion in *oriB* |
| V3d | 3 | 109449 - 109553 | DEL | 105 | GGATG…GGAGC | - | *oriB* | large deletion in *oriB* |
| V3e | 1 | 62413 - 62484 | DEL | 72 | AAGAT…AGATC | - | *trnQ-UUG* - *accD* spacer | larger deletion in *accD* promotor/5'-UTR |
| V3e | 1 | 62643 - 62657 | INS | 15 | - | CCTTT…CTTTA | *trnQ-UUG* - *accD* spacer | insertion in *accD* promotor/5'-UTR |
| V3e | 1 | 62691 - 62708 | INS | 18 | - | GATCC…CTTTA | *trnQ-UUG* - *accD* spacer | insertion in *accD* promotor/5'-UTR |
| V3e | 1 | 62754 - 62786 | INS | 15 | - | CCTTT…CTTTA | *trnQ-UUG* - *accD* spacer | insertion in *accD* promotor/5'-UTR |
| V3e | 1 | 76757 | INS | 1 | - | T | *rps12* - *clpP* spacer | intergenic, single bp insertion |
| V3e | 1 | 95720 - 95767 | DEL | 48 | CCGAA…AAAAC | - | *ycf2* | 16 AS deletion at site 2 of Ycf2 |
| V3e | 1 | 98434 - 98474 | DEL | 42 | GAGGA…CAGAA | - | *ycf2* | 14 AS deletion at site 3 Ycf2 |
| V3e | 1 | 109065 - 109246 | DEL | 182 | TCCCC…GCATG | - | *oriB* | large deletion in *oriB* |
| V3e | 1 | 109439 - 109443 | DEL | 5 | ACATA | - | *oriB* | small deletion in *oriB* |
| V3e | 1 | 109449 - 109553 | DEL | 105 | GGATG…GGAGC | - | *oriB* | large deletion in *oriB* |
| V3f | 2 | 88454 | DEL | 1 | A | - | *rpl16* - *rps4* | intergenic, oligo(N) stretch |
| V3f | 2 | 60935 - 61008 | DEL | 74 | CATAG…GTTGA | - | *trnQ-UUG* - *accD* spacer | intergenic, larger deletion |
| V3f | 2 | 62021 - 62095 | DEL | 75 | TGCCT…GATGA | - | *trnQ-UUG* - *accD* spacer | larger deletion in *accD* promotor/5'-UTR |
| V3f | 2 | 62739 - 62753 | DEL | 15 | CCTTT…CTTTA | - | *trnQ-UUG* - *accD spacer* | deletion in *accD* promotor/5'-UTR |
| V3f | 2 | 62981 - 63124 | DEL | 144 | TAAGG…GATAA | - | *accD* | 48 AS deletion in AccD |
| V3f | 2 | 109065 - 109246 | DEL | 182 | TCCCC…GCATG | - | *oriB* | large deletion in *oriB* |
| V3f | 2 | 134801 - 134821 | DEL | 21 | ACGTT…TTTTG | - | *ycf1* | 7 AS deletion in Ycf1 |
| V3g | 4 | 5720 | INS | 1 | - | T | *rps16* intron | intron, oligo(N) stretch |
| V3g | 4 | 71718 | INS | 1 | - | A | *psbE* - *petL* spacer | intergenic, oligo(N) stretch |
| V3g | 4 | 63434 - 63466 | INS | 33 | - | TAAGC…GATAA | *accD* | 11 AS insertion in AccD |
| V3g | 4 | 92483 - 92714 | DEL | 232 | CAAAA…TAACA | - | *trnI-CAU* - *ycf2* spacer | large deletion in *ycf2* promotor/5'-UTR |
| V3g | 4 | 98434 - 98706 | DEL | 267 | GAGGA…CAGAA | - | *ycf2* | 89 AS deletion at site 3 Ycf2 |

**Table S1** (continued)

| Variant | Inheritance class[2)] | Alignment position[3)] | Type | size [bp] | Allele in WT I-johSt | Allele in mutant I-johSt variant | Gene/Region | Type/Impact[4)] |
|---|---|---|---|---|---|---|---|---|
| V3h | 1 | 16405 | INS | 1 | - | A | *trnF-GGA* - *trnL-UAA* spacer | intergenic, oligo(N) stretch |
| V3h | 1 | 71172 | INS | 1 | - | T | *psbE* - *petL* spacer | intergenic, oligo(N) stretch |
| V3h | 1 | 62724 - 62753 | DEL | 30 | CCTTT…CTTTA | - | *trnQ-UUG* - *accD* spacer | deletion in *accD* promotor/5'-UTR |
| V3h | 1 | 63001 - 63072 | DEL | 72 | CTAAA…AGGGC | - | *accD* | 24 AS deletion in AccD |
| V3h | 1 | 98413 - 98475 | DEL | 63 | GAGGA...CAGAA | - | *ycf2* | 21 AS deletion at site 3 Ycf2 |
| V3h | 1 | 98593 - 98613 | DEL | 21 | GAGGA…CAGAA | - | *ycf2* | 7 AS deletion at site 3 Ycf2 |
| V3h | 1 | 98638 - 98661 | DEL | 24 | GATGA…CAGAA | - | *ycf2* | 8 AS deletion at site 3 Ycf2 |
| V3h | 1 | 98683 - 98685 | INS | 3 | - | GAT | *ycf2* | 1 AS insertion at site 3 Ycf2 |
| V3h | 1 | 98693 | SNP | 1 | T | G | *ycf2* | codon 1982, V:GTA to G:GGA |
| V3h | 1 | 109065 - 109246 | DEL | 182 | TCCCC…GCATG | - | *oriB* | large deletion in *oriB* |
| V3h | 1 | 109393 - 109653 | DEL | 261 | CGACG…TAGGA | - | *oriB* | large deletion in *oriB* |
| V4b | 2 | 76523 | SNP | 1 | T | G | *rps12* - *clpP* spacer | intergenic, SNP |
| V4b | 2 | 76536 | SNP | 1 | C | G | *rps12* - *clpP* spacer | intergenic, SNP |
| V4b | 2 | 60935 - 61008 | DEL | 74 | CATAG…GTTGA | - | *trnQ-UUG* - *accD* spacer | intergenic, larger deletion |
| V4b | 2 | 62485 - 62556 | INS | 72 | - | AAGAT…AGATC | *trnQ-UUG* - *accD* spacer | larger insertion in *accD* promotor/5'-UTR |
| V4b | 2 | 76526 - 76532 | INS | 7 | - | TTAAGAA | *rps12* - *clpP* spacer | Intergenic, small insertion |
| V4b | 2 | 109065 - 109246 | DEL | 182 | TCCCC…GCATG | - | *oriB* | large deletion in *oriB* |
| V4b | 2 | 109439 - 109443 | DEL | 5 | ACATA | - | *oriB* | small deletion in *oriB* |
| V4b | 2 | 109449 - 109553 | DEL | 105 | GGATG…GGAGC | - | *oriB* | large deletion in *oriB* |
| V4c | 1 | 16405 | INS | 1 | - | A | *trnF-GGA* - *trnL-UAA* spacer | intergenic, oligo(N) stretch |
| V4c | 1 | 60935 - 61008 | DEL | 74 | CATAG…GTTGA | - | *trnQ-UUG* - *accD* spacer | intergenic, larger deletion |
| V4c | 1 | 119103 | INS | 1 | - | A | *ndhF* - *rpl32* spacer | intergenic, oligo(N) stretch |
| V4c | 1 | 109065 - 109246 | DEL | 182 | TCCCC…GCATG | - | *oriB* | large deletion in *oriB* |
| V4c | 1 | 109439 - 109443 | DEL | 5 | ACATA | - | *oriB* | small deletion in *oriB* |
| V4c | 1 | 109449 - 109553 | DEL | 105 | GGATG…GGAGC | - | *oriB* | large deletion in *oriB* |
| V7a | 2 | 6005 | SNP | 1 | G | A | *rps16* | intron, SNP |
| V7a | 2 | 66691 | DEL | 1 | T | - | *ycf4* - *cemA* spacer | intergenic, oligo(N) stretch |

**Table S1** (continued)

| Variant | Inheritance class[2)] | Alignment position[3)] | Type | size [bp] | Allele in WT I-johSt | Allele in mutant I-johSt variant | Gene/Region | Type/Impact[4)] |
|---|---|---|---|---|---|---|---|---|
| VC1 | 4 | 15551 | INS | 1 | - | T | *ndhJ - trnF-GGA* spacer | intergenic, oligo(N) stretch |
| VC1 | 4 | 22673 | INS | 1 | - | T | *ycf3 - psaA* spacer | intergenic, oligo(N) stretch |
| VC1 | 4 | 28782 | SNP | 1 | C | T | *trnG-GCC* | tRNA |
| VC1 | 4 | 29904 | INS | 1 | - | T | *trnS-UGA - psbC* | intergenic, oligo(N) stretch |
| VC1 | 4 | 33524 | INS | 1 | - | T | *psbD - trnT-GGU* | intergenic, oligo(N) stretch |
| VC1 | 4 | 34068 | DEL | 1 | A | - | *trnT-GGU - trnE-UUC* | intergenic, oligo(N) stretch |
| VC1 | 4 | 49809 | INS | 1 | - | T | *rpoC2 - rps2* spacer | intergenic, oligo(N) stretch |
| VC1 | 4 | 54037 | INS | 1 | - | A | *atpF* intron | intron, oligo(N) stretch |
| VC1 | 4 | 58868 - 58869 | INS | 2 | - | TT | *psbI - psbK* | intergenic, oligo(N) stretch |
| VC1 | 4 | 59516 - 59517 | INS | 2 | - | TT | *psbK - trnQ-UUG* | intergenic, oligo(N) stretch |
| VC1 | 4 | 62054 | SNP | 1 | T | G | *trnQ-UUG - accD spacer* | SNP in *accD* promotor/5'-UTR |
| VC1 | 4 | 62341 - 62484 | DEL | 144 | AAGAT…AGATC | - | *trnQ-UUG - accD spacer* | large deletion in *accD* promotor/5'-UTR |
| VC1 | 4 | 62610 - 62660 | DEL | 36 | GATCC...TAGAT | - | *trnQ-UUG - accD spacer* | deletion in *accD* promotor/5'-UTR |
| VC1 | 4 | 62754 - 62897 | INS | 144 | - | CCTTT…CTTTA | *trnQ-UUG - accD spacer* | large insertion in *accD* promotor/5'-UTR |
| VC1 | 4 | 62981 - 63124 | DEL | 144 | TAAGG…GATAA | - | *accD* | 48 AS deletion in AccD |
| VC1 | 4 | 63264 - 63293 | INS | 30 | - | TATGA…ATGAG | *accD* | 10 AS insertion in AccD |
| VC1 | 4 | 63434 - 63532 | INS | 99 | - | TAAGC…GATAA | *accD* | 33 AS insertion in AccD |
| VC1 | 4 | 63563 - 63583 | DEL | 21 | AGAAG…AAGGA | - | *accD* | 7 deletion in AccD |
| VC1 | 4 | 67064 | INS | 1 | - | A | *ycf4 - cemA* spacer | intergenic, oligo(N) stretch |
| VC1 | 4 | 71718 | INS | 1 | - | A | *psbE - petL* spacer | intergenic, oligo(N) stretch |
| VC1 | 4 | 73173 | DEL | 1 | G | - | *trnP-UGG - psaJ* spacer | intergenic, single bp deletion |
| VC1 | 4 | 73186 | SNP | 1 | G | C | *trnP-UGG - psaJ* spacer | intergenic, SNP |
| VC1 | 4 | 73177 - 73182 | DEL | 6 | TAAGAA | - | *trnP-UGG - psaJ* spacer | intergenic, small deletion |
| VC1 | 4 | 92483 - 92714 | DEL | 232 | CAAAA…TAACA | - | *trnI-CAU - ycf2* spacer | large deletion in *ycf2* promotor/5'-UTR |
| VC1 | 4 | 98392 - 98474 | DEL | 84 | GAGGA…CAGAA | - | *ycf2* | 28 AS deletion at site 3 Ycf2 |
| VC1 | 4 | 98521 - 98589 | DEL | 69 | GATGA…CAGAA | - | *ycf2* | 23 AS deletion at site 3 Ycf2 |
| VC1 | 4 | 98638 - 98661 | DEL | 24 | GATGA…CAGAA | - | *ycf2* | 8 AS deletion at site 3 Ycf2 |
| VC1 | 4 | 98683 - 98685 | INS | 3 | - | GAT | *ycf2* | 1 AS insertion at site 3 Ycf2 |
| VC1 | 4 | 98707 - 98709 | DEL | 3 | GAT | - | *ycf2* | 1 AS deletion at site 3 Ycf2 |
| VC1 | 4 | 109065 - 109246 | DEL | 182 | TCCCC…GCATG | - | *oriB* | large deletion in *oriB* |
| VC1 | 4 | 109393 - 109653 | DEL | 261 | CGACG…TAGGA | - | *oriB* | large deletion in *oriB* |
| VC1 | 4 | 135265 - 135300 | DEL | 36 | GCTAG…AGCAG | - | *ycf1* | 12 AS deletion in Ycf1 |
| VC1 | 4 | 135756 - 135776 | INS | 21 | - | TTTTT…TATTT | *ycf1* | 7 AS insertion in Ycf1 |

[1)]For details on the WT plastome see Table S7

[2)]For details and definition of these classes in the variants see SI Table 10 and SI Text.

[3)]Alignment provided in Dataset S2

[4)]small insertion/deletion = 5 - 10 bp, insertion/deletion = 10 - 50 bp, larger insertion/deletion = 50 - 100 bp, and large insertion/deletion > 100 bp

**Table S2.** Summary of the mutations found in the very weak variants of III-lamS[1)]

| Variant | Alignment position[2)] | Type | size [bp] | Allele in WT III-lamS | Allele in mutant III-lamS variant | Gene/Region | Type/Impact | Shared with weak I-johSt variants (V3g and VC1)[3)] | Shared with weak plastome IV WTs[4)] |
|---|---|---|---|---|---|---|---|---|---|
| III-V2 | 2240 - 2242 | INS | 3 | - | CTA | *matK* (in *trnK-UUU* intron) | 1 AS insertion (intron) | no | no |
| III-V1 | 4424 | INS | 1 | - | A | *trnK-UUU* - *rps16* spacer | intergenic, oligo(N) stretch | no | yes |
| III-V1 | 6633 | SNP | 1 | C | G | *rps16* - *rbcL* spacer | intergenic, SNP | no | no |
| III-V2 | 13266 | INS | 1 | - | T | *trnV-UAC* - *ndhC* spacer | intergenic, single bp insertion | no | no |
| III-V2 | 16419 | INS | 1 | - | A | *trnF-GAA* - *trnL-UAA* spacer | intergenic, oligo(N) stretch | no | no |
| III-V1 | 17109 | INS | 1 | - | T | *trnL-UAA* | intron, oligo(N) stretch | no | no |
| III-V2 | 33494 | INS | 1 | - | T | *psbD* - *trnT-GGU* | intergenic, oligo(N) stretch | no | no |
| III-V1 | 62409 - 62591 | DEL | 183 | AAGAT…AGATC | - | *trnQ-UUG* - *accD* spacer | large deletions in *accD* promotor/5'-UTR | no | yes |
| III-V1 | 62615 - 62668 | DEL | 54 | CCTTT…TAGAT | - | | | | |
| III-V1 | 66625 | INS | 1 | - | A | *ycf4* - *cemA* spacer | intergenic, oligo(N) stretch | no | no |
| III-V2 | 66625 | INS | 1 | - | A | | | | |
| III-V2 | 71277 | INS | 1 | - | A | *psbE* - *petL* spacer | intergenic, oligo(N) stretch | yes | no |
| III-V1 | 84138 | SNP | 1 | C | A | *rpoA* | codon 43, D:GAT to Y:tAT | no | no |
| III-V1 | 89022 | INS | 1 | - | T | *rpl22* | 5 AS C-terminal extension, oligo(N) stretch | no | yes |
| III-V2 | 89022 | INS | 1 | - | T | | | | |
| III-V1 | 92259 - 92502 | DEL | 244 | ACAAA…ATAAC | - | *trnI-CAU* - *ycf2* spacer | large deletions in ycf2 promotor/5'-UTR | yes | yes |
| III-V2 | 92259 - 92502 | DEL | 244 | ACAAA…ATAAC | - | | | | |
| III-V1 | 98156 - 98329 | DEL | 174 | GGAAG…GAAGA | - | *ycf2* | 116 AS and 146 AS deletion, respectively, at site 3 of Ycf2 | yes | yes |
| III-V1 | 98375 - 98548 | DEL | 174 | GGAAG…GAAGA | - | | | | |
| III-V2 | 98175 - 98175 | DEL | 441 | GAGGA…CAGAA | - | | | | |
| III-V1 | 99078 - 99101 | DEL | 24 | GAAGA…AAGAG | - | *ycf2* | 8 AS deletion | no | no |
| III-V2 | 111272 - 111274 | INS | 2 | - | GT | *trnA-UGC* - *rrn23* | intergenic, 2 bp insertion | no | no |
| III-V1 | 119750 | DEL | 1 | C | - | *trnL-UAG* - *ccsA* | intergenic, oligo(N) stretch | no | no |
| III-V2 | 119750 | DEL | 1 | C | - | | | | |
| III-V2 | 129113 | INS | 1 | - | T | *rps15* - *ycf1* | intergenic, oligo(N) stretch | no | no |
| III-V2 | 134224 - 134244 | DEL | 21 | GTTTA…ATTTC | - | *ycf1* | 7 AS deletion | no | yes |
| III-V1 | 134301 - 134321 | DEL | 21 | TGACT…GCTTT | - | *ycf1* | 7 AS deletion | no | yes |
| III-V2 | 134301 - 134321 | DEL | 21 | TGACT…GCTTT | - | | | | |

[1)]For details on the WT and variants chloroplast genomes, their inheritance class and phenotype in the nuclear background of johansen Standard see Fig S15 and S19, Tables S7, S8 and S11 and SI Text.

[2)]Alignment provided in Dataset S2.

[3)]For details see Table S1and Dataset S2.

[4)]For details see Dataset S2.

**Table S3.** Genotypes used for the individual experimental series of the lipid-level measurements described in this work[1)]

| Variant/mutant/ plastome[2)] | Experiment1 | Experiment 2 | Experiment 3 | # total samples measured | Inheritance class | Phenotype in johansen Standard |
|---|---|---|---|---|---|---|
| I-johSt | x | x | x | 15 | 1 | green |
| I-hookdV | - | x | x | 10 | 1 | green |
| III-lamS | - | x | x | 10 | 1 | *virescent* |
| I-chi | - | x | x | 10 | 1 | white to yellowish |
| V1c | x | - | x | 10 | 1 | green |
| V2a | - | x | x | 10 | 1 | green |
| V2g | x | - | x | 10 | 1 | green |
| V3e | x | - | x | 10 | 1 | green |
| V3c | x | - | x | 10 | 2 | green |
| V3d | - | x | x | 10 | 3 | green |
| VC1 | x | x | x | 15 | 4 | green |
| V3g | x | - | x | 10 | 4 | green |
| IV-atroSt | x | x | x | 15 | 4 | green |
| IV-delta | - | x | x | 10 | 4 | white to yellowish |
| III-V1 | - | x | - | 5 | 5 | *virescent* |
| III-V2 | - | x | - | 5 | 5 | *virescent* |

**x** = included in the experiment  **-** = not included in the experiment

[1)]For details on the chloroplast genomes, their inheritance class, carried mutations and phenotype of the nuclear background of johansen Standard see Fig. 2, Figs. S15, S18 and S19, Tables S1, S2, S6-S11, and SI Text.

**Table S4.** Average weight and corresponding *p*-value of the 102 lipids/molecules analyzed in the LASSO regression model to predict inheritance strengths. The 20 predictive lipid (absolute average weight greater than one standard deviation of all lipids/molecules) are marked in italics

| Lipid/ molecule | Average weight | Adjusted *p*-value[1)] | Lipid/ molecule | Average weight | Adjusted *p*-value[1)] | Lipid/ molecule | Average weight | Adjusted *p*-value[1)] |
|---|---|---|---|---|---|---|---|---|
| *TAG 54.3* | *-2.89* | *$1.27 \times 10^{-30}$* | SQDG 34.3 | -0.04 | $1.27 \times 10^{-1}$ | FA 28.0 | 0.06 | $1.50 \times 10^{-4}$ |
| *PC 36.3* | *-1.93* | *$6.96 \times 10^{-46}$* | TAG 50.2 | -0.03 | 1 | CoQ 9 | 0.07 | $9.11 \times 10^{-6}$ |
| *PC 36.4* | *-1.66* | *$3.05 \times 10^{-27}$* | SQDG 36.2 | -0.03 | $6.92 \times 10^{-2}$ | PC 34.5 | 0.07 | $2.90 \times 10^{-1}$ |
| *DGDG 36.4* | *-1.32* | *$8.84 \times 10^{-31}$* | TAG 54.7 | -0.02 | 1 | MGDG 34.2 | 0.08 | $9.61 \times 10^{-4}$ |
| *DGDG 36.5* | *-1.30* | *$1.01 \times 10^{-33}$* | MGDG 36.6 | -0.02 | 1 | SQDG 36.6 | 0.11 | $6.52 \times 10^{-3}$ |
| *DGDG 34.5* | *-1.29* | *$1.16 \times 10^{-28}$* | TAG 60.3 | -0.02 | 1 | TAG 58.3 | 0.12 | $1.24 \times 10^{-4}$ |
| *TAG 60.6* | *-1.29* | *$2.82 \times 10^{-38}$* | TAG 50.1 | -0.01 | 1 | PC 36.6 | 0.13 | $3.54 \times 10^{-5}$ |
| *TAG 52.3* | *-1.19* | *$5.40 \times 10^{-31}$* | TAG 56.6 | -0.01 | 1 | TAG 56.7 | 0.16 | $6.84 \times 10^{-2}$ |
| *PC 36.2* | *-0.97* | *$1.18 \times 10^{-18}$* | TAG 52.1 | 0.00 | 1 | PE 36.2 | 0.16 | 1 |
| SQDG 36.4 | -0.63 | $2.77 \times 10^{-29}$ | TAG 54.5 | 0.00 | 1 | PC 34.2 | 0.16 | $6.19 \times 10^{-8}$ |
| FA 30.0 | -0.52 | $5.76 \times 10^{-33}$ | DGDG 36.3 | 0.00 | 1 | MGDG 36.2 | 0.17 | $5.08 \times 10^{-6}$ |
| TAG 52.9 | -0.49 | $2.07 \times 10^{-22}$ | TAG 48.0 | 0.00 | 1 | DGDG 32.0 | 0.19 | $4.82 \times 10^{-8}$ |
| TAG 54.8 | -0.43 | $2.80 \times 10^{-13}$ | chlorophyll b | 0.00 | 1 | PE 36.3 | 0.20 | $1.05 \times 10^{-7}$ |
| PC 32.1 | -0.40 | $8.73 \times 10^{-13}$ | DGDG 34.2 | 0.00 | 1 | SQDG 36.3 | 0.22 | $1.60 \times 10^{-7}$ |
| DGDG 34.1 | -0.31 | $3.82 \times 10^{-15}$ | TAG 50.5 | 0.00 | 1 | PE 36.4 | 0.24 | $2.88 \times 10^{-7}$ |
| PE 36.6 | -0.26 | $1.64 \times 10^{-8}$ | TAG 52.7 | 0.00 | 1 | CoQ10 | 0.26 | $1.85 \times 10^{-7}$ |
| MGDG 34.3 | -0.25 | $7.70 \times 10^{-6}$ | TAG 54.4 | 0.00 | 1 | TAG 52.6 | 0.32 | $1.15 \times 10^{-11}$ |
| PC 32.2 | -0.25 | $1.53 \times 10^{-5}$ | TAG 56.3 | 0.00 | 1 | TAG 50.3 | 0.40 | $7.34 \times 10^{-14}$ |
| SQDG 36.5 | -0.20 | $7.40 \times 10^{-9}$ | TAG 56.4 | 0.00 | 1 | pheophytin | 0.49 | $8.42 \times 10^{-39}$ |
| TAG 52.5 | -0.19 | $1.12 \times 10^{-10}$ | TAG 60.2 | 0.00 | 1 | PC 34.3 | 0.60 | $4.03 \times 10^{-19}$ |
| FA 26.0 | -0.16 | $1.15 \times 10^{-11}$ | TAG 60.4 | 0.00 | 1 | TAG 56.2 | 0.61 | $2.89 \times 10^{-15}$ |
| DGDG 36.2 | -0.14 | $5.00 \times 10^{-2}$ | TAG 54.2 | 0.00 | 1 | MGDG 36.3 | 0.63 | $2.21 \times 10^{-46}$ |
| PG 34.2 | -0.13 | $6.70 \times 10^{-3}$ | TAG 50.6 | 0.00 | 1 | PC 32.0 | 0.65 | $3.71 \times 10^{-21}$ |
| PI 34.3 | -0.11 | $1.53 \times 10^{-8}$ | TAG 54.6 | 0.00 | 1 | *TAG 50.7* | *0.80* | *$2.43 \times 10^{-20}$* |
| PE 34.2 | -0.11 | $1.10 \times 10^{-1}$ | TAG 54.1 | 0.01 | 1 | *TAG 56.1* | *0.80* | *$1.14 \times 10^{-34}$* |
| TAG 52.2 | -0.10 | $5.54 \times 10^{-5}$ | MGDG 36.5 | 0.01 | $1.72 \times 10^{-1}$ | *DGDG 34.4* | *0.94* | *$9.52 \times 10^{-29}$* |
| TAG 60.5 | -0.09 | $1.68 \times 10^{-2}$ | TAG 50.4 | 0.01 | 1 | *PC 34.1* | *0.96* | *$3.43 \times 10^{-29}$* |
| TAG 56.5 | -0.09 | $6.92 \times 10^{-2}$ | TAG 48.1 | 0.02 | 1 | *PE 36.5* | *1.01* | *$1.26 \times 10^{-23}$* |
| PI 34.2 | -0.09 | $3.75 \times 10^{-8}$ | TAG 52.4 | 0.02 | $7.39 \times 10^{-1}$ | *MGDG 36.4* | *1.02* | *$2.77 \times 10^{-34}$* |
| DGDG 34.3 | -0.08 | $2.90 \times 10^{-2}$ | chlorophyll a | 0.03 | 1 | *TAG 58.4* | *1.03* | *$7.23 \times 10^{-32}$* |
| PC 34.4 | -0.08 | $1.16 \times 10^{-2}$ | TAG 52.8 | 0.04 | 1 | *PE 34.3* | *1.03* | *$3.58 \times 10^{-23}$* |
| TAG 58.5 | -0.08 | $6.38 \times 10^{-7}$ | TAG 58.1 | 0.05 | $7.72 \times 10^{-6}$ | *PG 34.3* | *1.12* | *$2.13 \times 10^{-32}$* |
| FA 24.0 | -0.06 | $9.17 \times 10^{-8}$ | TAG 60.1 | 0.05 | $2.16 \times 10^{-4}$ | *TAG 54.9* | *2.39* | *$1.30 \times 10^{-55}$* |
| DGDG 36.6 | -0.06 | $5.02 \times 10^{-1}$ | PC 36.5 | 0.06 | $3.40 \times 10^{-13}$ | *TAG 58.2* | *2.68* | *$1.75 \times 10^{-37}$* |

[1)] *p*-values represent significance of deviation of average coefficients (weights) from zero as obtained from the 100 cross-validation runs. Raw *p*-values have been adjusted for multiple testing (102 lipids) using Benjamini and Hochberg procedure.

**Table S5.** Enrichment/depletion of lipid/molecule classes in the set of 20 predictive lipids based on Fisher's exact test *p*-value and odds ratio

| Lipid class | Odds ratio | *p*-value[1)] | # predictive lipids/molecules | # total lipids/ molecules in classes |
|---|---|---|---|---|
| PG | 4.18 | 0.35 | 1 | 2 |
| DGDG | 2.64 | 0.22 | 4 | 11 |
| PC | 2.01 | 0.28 | 4 | 13 |
| PE | 1.70 | 0.62 | 2 | 7 |
| TAG | 0.81 | 0.80 | 8 | 45 |
| MGDG | 0.67 | 1.00 | 1 | 7 |
| FA | 0.00 | 0.58 | 0 | 4 |
| SQDG | 0.00 | 059 | 0 | 6 |
| PI | 0.00 | 1.00 | 0 | 2 |
| chlorophyll | 0.00 | 1.00 | 0 | 2 |
| CoQ | 0.00 | 1.00 | 0 | 2 |
| pheophytin | 0.00 | 1.00 | 0 | 1 |

[1)] *p*-values have not been corrected for multiple testing as none of the raw *p*-values proved below $p < 0.05$. After correction, *p*-values shall be set to 1.

**Table S6.** *Oenothera* strains with native wild type chloroplasts used in this work[1)]

| Species | Strain[1)] | Locality | Collection/ isolation date | Collector/ isolated by | Reference |
|---|---|---|---|---|---|
| *O. elata* ssp. *hookeri* | johansen Standard | USA, CA, Sutter Co., roadside between Nicholas and Yuba City | 1927 | C. B. Wolf | (65) |
| *O. elata* ssp. *hookeri* | hookeri de Vries | USA, CA, Alameda Co., near Berkeley | 1904 | H. de Vries | (66) |
| *O. villosa* ssp. *villosa* | bauri Standard | Poland, Kujawsko-Pomorskie, near Toruń | before 1942 | R. Hölscher | (67) |
| *O. glazioviana* | blandina de Vries[2)] | chromosome translocation mutant of the original lamarckiana de Vries[3)] | isolated in 1908 | H. de Vries | (68, 69) |
| *O. glazioviana* | deserens de Vries[4)] | chromosome translocation mutant of the original lamarckiana de Vries[3)] | isolated in 1913 | H. de Vries | (68, 70, 71) |
| *O. glazioviana* | *r*/*r*-lamarckiana Sweden | Sweden, Skåne Län, garden in Almaröd | 1906 | N. Heribert-Nilsson | (72) |
| *O. biennis* | chicaginensis de Vries | USA, IL, Cook Co., Chicago, near Jackson Park | 1904 | H. de Vries | (66) |
| *O. biennis* | biennis Muenchen | Germany, Bavaria, Munich, Nymphenburg Garden | 1914 | O. Renner | (73) |
| *O. biennis* | rubricaulis Thorn | Poland, Kujawsko-Pomorskie, Vistula River near Toruń | before 1941 | R. Hölscher | (74) |
| *O. biennis* | suaveolens Standard | France, Seine-et-Marne, forest of Fontainebleau | 1912 | L. Blaringhem | (75) |
| *O. biennis* | suaveolens Grado | Italy, Friuli-Venezia Giulia, dune near Grado at the Adriatic sea | before 1950 | H. Zeidler | (76) |
| *O. oakesiana* | ammophila Standard | Germany, Schleswig-Holstein, Helgoland | 1922 | E. Hoeppener | (77) |
| *O. oakesiana* | *r*/*r*-syrticola Ulm | Germany, Baden-Württemberg, Danube River near Ulm | 1917 | O. Renner | (78) |
| *O. parviflora* | rubricuspis Standard | Germany, Hessen, railway embankment between Neu-Isenburg and Luisa near Frankfurt on the Main | 1942/1943 | O. Burck, F. Laibach, and E. Fischer | (79) |
| *O. parviflora* | silesiaca Standard | Poland, Dolnośląskie, bank of Bóbr River near Nowogrodziec | 1937 | O. Renner | (74) |
| *O. parviflora* | atrovirens Standard | USA, NY, Erie Co., Sandy Hill near Lake George[5)] | 1902/1903 | D. T. MacDouglas | (66, 80) |

[1)]For details on *Oenothera* taxonomy and strain designation see (68, 81, 82).

[2)]*Syn*: *O. blandina* de Vries, *O. lamarckiana* de Vries mut. *blandina*, *O. lamarckiana* de Vries mut. *veluntina.*

[3)]The original lamarckiana de Vries was first collected in 1886 at Graveland near Hilversum (The Netherlands, Noord-Holland) by H. de Vries (66, 83).

[4)]*Syn*: *O. deserens* de Vries, *O. lamarckiana* de Vries mut. *deserens.*

[5)]According to Renner (80) this line was originally "received from Amsterdam" by N. v. Gescher in 1907. The material is quite likely identical to that collected by D. T. MacDouglas in 1902/1903 as described in (66). Also see (84).

**Table S7.** Summary on the wild type *Oenothera* chloroplast genomes used in this work[1)]

| Species | Strain[1)] | Abbreviation plastome | Basic plastome[2)] | [%] biparental inheritance (biennis white)[3)] | [%] biparental inheritance (blandina white)[3)] | Inheritance class (Schötz)[4)] | Inheritance class (this work)[5)] | GenBank/EMBL accession number |
|---|---|---|---|---|---|---|---|---|
| *O. elata* ssp. *hookeri* | johansen Standard | I-johSt | I | n/a | n/a | strong | 1 | AJ271079.4 |
| *O. elata* ssp. *hookeri* | hookeri de Vries | I-hookdV | I | 0.0 | 8.2 | strong | 1 | KT881170.1 |
| *O. villosa* ssp. *villosa* | bauri Standard | I-bauriSt | I | 7.4 | 43.8 | strong | 3 | KX687910.1 |
| *O. glazioviana* | blandina de Vries | III-blandV | III | 0.0 | 14.1 | strong | 1 | KT881171.1 |
| *O. glazioviana* | deserens de Vries | III-desdV | III | 1.3 | 21.4 | strong | 1 | KT881172.1 |
| *O. glazioviana* | *r*/*r*-lamarckiana Sweden | III-lamS | III | 3.1 | 26.4 | strong | 1 | EU262890.2 |
| *O. biennis* | chicaginensis de Vries | III-chicdV | III | 5.0 | 24.4 | strong | 1 | KX687913.1 |
| *O. biennis* | biennis Muenchen | II-biM | II | 22.2 | 53.8 | intermediate | 3 | KU521375.1 |
| *O. biennis* | rubricaulis Thorn | II-rcauTh | II | 23.2 | 41.8 | intermediate | 3 | KX687914.1 |
| *O. biennis* | suaveolens Standard | II-suavSt[6)] | II | 26.7 | 49.9 | intermediate | 3 | KX687915.1 |
| *O. biennis* | suaveolens Grado | II-suavG[6)] | II | n/a | n/a | intermediate | 3 | EU262889.2 |
| *O. oakesiana* | ammophila Standard | IV-ammSt | IV | 43.9 | 58.1 | weak | 4 | KT881176.1 |
| *O. oakesiana* | *r*/*r*-syrticola Ulm | IV-syrtU | IV | 48.7 | 63.3 | weak | 4 | KX687918.1 |
| *O. parviflora* | rubricuspis Standard | IV-rcuSt | IV | 52.3 | 68.7 | weak | 4 | KX687916.1 |
| *O. parviflora* | silesiaca Standard | IV-silSt | IV | 53.2 | 67.2 | weak | 4 | KX687917.1 |
| *O. parviflora* | atrovirens Standard | IV-atroSt | IV | 70.1 | 72.6 | weak | 4 | EU262891.2 |

[1)]For details on *Oenothera* taxonomy and strain designation see (68, 81, 82).

[2)]In *Oenothera,* five genetically distinguishable plastome types (I-V) can be recognized based on their compatibility with three nuclear genomes (A, B, C) in either homozygous (AA, BB, CC) or stable heterozygous (AB, AC, BC) states. The basic plastome genotype is accompanied by a given inheritance strength (strong, intermediate and weak). Basic plastome and nuclear genome type are important factors of species definition in *Oenothera*. For details see, e.g. (24, 34, 60, 68, 81, 85, 86).

[3)]Percentage of variegated seedlings, heteroplasmic due to the paternal transmission of the bleached chloroplast mutants “biennis white” or “blandina white” and the maternal transmission of the green wild chloroplast of the seed parent. Data accoding to Schötz (34). For details see therein, reviews in (24, 68) and SI Text.

[4)]Class of inheritance strength as determined by F. Schötz. For reviews see (24, 68, 87).

[5)]For details on the definition of these classes see SI Text.

[6)]The chloroplast genomes of the two suaveolens stains are identical.

**Table S8.** Wild type chloroplast genomes introgressed into the nuclear background of johansen Standard used in this work[1)]

| Plastome[2)] | Basic plastome[3)] | Inheritance class (Schötz)[4)] | Inheritance class (this work)[5)] | Chloroplast donor strain[6)] | Phenotype in the nuclear background of johansen Standard | Produced by | Reference |
|---|---|---|---|---|---|---|---|
| I-johSt | I | strong | 1 | johansen Standard | green | wild type | (60, 65) |
| I-hookdV | I | strong | 1 | hookeri de Vries | green | S. Greiner | (2); this work |
| II-suavG | II | intermediate | 3 | suaveolens Grado | green | W. Stubbe | (60, 85) |
| III-lamS | III | strong | 1 | *r*/*r*-lamarckian Sweden | *viresencet* | W. Stubbe | (3) |
| IV-atroSt | IV | weak | 4 | atrovirens Standard | green | W. Stubbe | (60, 85) |

[1)]Wild type plastids with different transmission efficiencies were introgressed into the constant nuclear background of the johansen Standard strain (*O. elata* ssp. *hookeri*). See Table S6 for details on this line.

[2)]See Table S7 for details.

[3)]In *Oenothera*, five genetically distinguishable plastome types (I-V) can be recognized based on their compatibility with three nuclear genomes (A, B, C) in either homozygous (AA, BB, CC) or stable heterozygous (AB, AC, BC) states. The basic plastome genotype is accompanied by a given inheritance strength (strong, intermediate and weak). Basic plastome and nuclear genome type are important factors of species definition in *Oenothera*. For details see, e.g. (24, 34, 60, 68, 81, 85, 86).

[4)]Classes of inheritance strength as determined by F. Schötz. For reviews see (24, 68, 87).

[5)]For details on the definition of these classes, see SI Text.

[6)]For details on the donor strains, see Table S6.

**Table S9.** Spontaneous mutants in the chloroplast gene *psaA* in the nuclear background of johansen Standard used in this work[1)]

| Mutant/ plastome | Wild type background plastome[2)] | Basic plastome[3)] | Inheritance class (Schötz)[4)] | Inheritance class (this work)[5)] | Phenotype | Mutation | Reference |
|---|---|---|---|---|---|---|---|
| I-chi | I-hookdV | I | strong | 1 | white to yellowish | 5-bp duplication (CCGCT), +1420 to +1424 *psaA*, truncated PsaA | (2) |
| IV-delta | IV-atroSt | IV | weak | 4 | white to yellowish | 5-bp duplication (TTAAC), +667 to +671 *psaA*, truncated PsaA | (2) |

[1)]Plastome mutants with different transmission efficiencies were introgressed into the constant nuclear background of the johansen Standard strain (*O. elata* ssp. *hookeri*) by W. Stubbe. See Table S6 for details on this line.

[2)]See Table S5 for details on the wild type plastomes.

[3)] )In *Oenothera*, five genetically distinguishable plastome types (I-V) can be recognized based on their compatibility with three nuclear genomes (A, B, C) in either homozygous (AA, BB, CC) or stable heterozygous (AB, AC, BC) states. The basic plastome genotype is accompanied by a given inheritance strength (strong, intermediate and weak). Basic plastome and nuclear genome type are important factors of species definition in *Oenothera*. For details see, e.g. (24, 34, 60, 68, 81, 85, 86).

[4)]Classes of inheritance strength as determined by F. Schötz. For reviews see (24, 68, 87).

[5)]For details on the definition of these classes see SI Text.

**Table S10.** Inheritance strength of the wild type chloroplast genomes I-johSt, II-suavG, IV-atroSt[1)] and the green variants of plastomes I-johSt, determined in crosses to the chloroplast mutants I-chi and IV-delta[2)] as pollen donor and seed parents, respectively[3)]

| Variant/ plastome | Generations of *pm* mutagenesis | [%] biparental inheritance (I-chi)[4)] | [%] biparental inheritance (IV-delta)[5)] | Inheritance class (Schötz)[6,7)] | Inheritance class (this work)[7)] | Sequence determined |
|---|---|---|---|---|---|---|
| I-johSt | wild type | 6.0 | 42.1 | strong | 1 | AJ271079.4 |
| II-suavG | wild type | 34.6 | n/a | intermediate | 3 | EU262889.2 |
| IV-atroSt | wild type | 55.4 | n/a | weak | 4 | EU262891.2 |
| V1a | one | 10.7 | 54.6 | strong to intermediate | 2 | yes |
| V1b | one | 7.9 | 31.6 | strong | 1 | yes |
| V1c | one | 5.0 | 40.7 | strong | 1 | yes |
| V2a | one | 13.1 | 36.6 | strong | 1 | yes |
| V2b | one | 14.9 | 28.1 | intermediate | 3 | yes |
| V2e | one | 15.9 | 49.0 | strong to intermediate | 2 | no |
| V2f | one | 7.4 | 41.8 | strong | 1 | no |
| V2g | one | 13.3 | 36.3 | strong | 1 | yes |
| V3a | one | 18.1 | 32.5 | intermediate | 3 | yes |
| V3b | one | 10.0 | 40.3 | strong | 1 | yes |
| V3c | one | 23.0 | 55.6 | strong to intermediate | 2 | yes |
| V3d | one | 14.7 | 35.1 | intermediate | 3 | yes |
| V3e | one | 8.0 | 44.4 | strong | 1 | yes |
| V3f | one | 15.2 | 56.8 | strong to intermediate | 2 | yes |
| V3g | one | 42.0 | 13.4 | weak | 4 | yes |
| V3h | one | 10.7 | 40.5 | strong | 1 | yes |
| V4a | one | 14.2 | 63.0 | strong to intermediate | 2 | no |
| V4b | one | 17.7 | 45.8 | strong to intermediate | 2 | yes |
| V4c | one | 10.7 | 35.3 | strong | 1 | yes |
| V4d | one | 10.0 | 36.7 | strong | 1 | no |
| V7a | one | 10.7 | 49.0 | strong to intermediate | 2 | yes |
| V7c | one | 27.9 | 42.0 | intermediate | 3 | no |
| V7d | one | 24.2 | 30.8 | intermediate | 3 | no |
| VC1 | several | 40.9 | 7.4 | weak | 4 | yes |

[1)]See Table S8 for details on the wild type plastomes.

[2)]See Table S9 for details on the chloroplast mutants.

[3)]All crosses were performed in the nuclear genetic background of *O. elata* ssp. *hookeri* strain johansen Standard. For details on the crosses see Materials and Methods, on the line see Table S6.

[4)]Percentage of variegated seedlings, heteroplasmic due to the paternal transmission of the bleached chloroplast mutant I-chi and the maternal transmission of the green wild type or variant chloroplast of the seed parent. For details see SI Text and Fig. 2

[5)]Percentage of variegated seedlings, heteroplasmic due to the paternal transmission of the green wild type or variant chloroplast and the maternal transmission of the bleached chloroplast mutant IV-delta of the seed parent. For details see SI Text and Fig. 2.

[6)]Classes of inheritance strength as determined by F. Schötz. For reviews see (24, 68, 87).

[7)]For details on the definition of the classes in the variants see SI Text.

**Table S11.** Wild type chloroplast genome III-lamS and its very weak variants[1)] created by the *plastome mutator* in crosses to I-johSt[1)] as pollen donor[2)]

| Variant/ plastome | Generations of *pm* mutagenesis | Cross to I-johSt as pollen donor | | Inheritance class (Schötz)[5,6)] | Inheritance class (this work)[6)] |
|---|---|---|---|---|---|
| | | [%] biparental inheritance[3)] | [%] paternal inheritnace[4)] | | |
| III-lamS | wild type | 6.0 | 0.0 | strong | 1 |
| III-V1 | several | 37.3 | 1.3 | very weak | 5 |
| III-V2 | several | 38.0 | 1.6 | very weak | 5 |

[1)]See Table S8 for details on the wild type plastomes. For sequences of the variants see Dataset S2.

[2)]All crosses were performed in the constant nuclear genetic background of *O. elata* ssp. *hookeri* strain johansen Standard. For details on the crosses see Materials and Methods, on the line see Table S6.

[3)]Percentage of variegated seedlings, heteroplasmic due to the paternal transmission of the green wild type chloroplast I-johSt and the maternal transmission of the *virescent* chloroplast of the seed parent. For details see SI Text and Fig. S19.

[4)]Percentage of homoplasmic green seedlings resulting from paternal inheritance of the green wild type chloroplast I-johSt that fully out-competed the *virescent* chloroplast of the seed parent. For details see SI Text and Fig. S19.

[5)] Classes of inheritance strength as determined by F. Schötz. For reviews see (24, 68, 87).

[6)]For details on the definition of the classes in the variants see SI Text.

**Table S12.** Oligonucleotides employed in the MassARRAY and SNPs distinguishing I-johSt (AJ271079.4) and I-hookdV/I-chi (KT881170.1; I/I assay) or I-johSt and IV-atroSt/IV-delta (EU262891.2; I/IV assay)

| **Primer name** | **Assay** | **Sequence (5' - 3')** | **SNP (position in I-johSt)** | **Function** |
|---|---|---|---|---|
| jh_atpI_W1-1 | I/I | ACGTTGGATGGCCCTTGCTAACCAAGTCAT | A/C (51,812) | PCR |
| jh_atpI_W1-2 | I/I | ACGTTGGATGCGTCCCTGTATACAATCTAC | | PCR |
| jh_atpI_W1UEP | I/I | CAATCTACATGTAGGATACTG | | Extension |
| jh_matK_W1-1 | I/I | ACGTTGGATGATCATTCCGGGTCGGCTTAC | T/C (2,235) | PCR |
| jh_matK_W1-2 | I/I | ACGTTGGATGTGCCTCTGATTGGATCGTTG | | PCR |
| jh_matK_W1UEP | I/I | GCGAAATTTTGTAATGCATTAGG | | Extension |
| jh_rpl32_W1-1 | I/I | ACGTTGGATGCGTATTCGTAAAACTATTTGG | T/G (119,269) | PCR |
| jh_rpl32_W1-2 | I/I | ACGTTGGATGCCAGTAGAAAGAGACTTCGC | | PCR |
| jh_rpl32_W1UEP | I/I | CGCTAACGAAAAAGCTTTCAAGG | | Extension |
| jh_rps16_W1-1 | I/I | ACGTTGGATGAGTTGACAATTTCGGTACTG | G/A (6,919) | PCR |
| jh_rps16_W1-2 | I/I | ACGTTGGATGAGTATGTCAAGTCAACGTCC | | PCR |
| jh_rps16_W1UEP | I/I | CATCCATATTCTAATCTACCCGT | | Extension |
| jh_rps19_W1-1 | I/I | ACGTTGGATGATAGGCAAATGCTCTTTTCC | G/C (89,483) | PCR |
| jh_rps19_W1-2 | I/I | ACGTTGGATGTAAGAACTTGGTCTAGGGCG | | PCR |
| jh_rps19_W1UEP | I/I | CGATTATACTTACAATGATCGG | | Extension |
| jh_trnK_W1-1 | I/I | ACGTTGGATGGCGATTGTATCTACACATAG | C/A (4,124) | PCR |
| jh_trnK_W1-2 | I/I | ACGTTGGATGAGACGCACTTAAAAGCCGAG | | PCR |
| jh_trnK_W1UEP | I/I | TTGAGTTAGCAACCCCCCC | | Extension |
| jh_trnQ_W1-1 | I/I | ACGTTGGATGCATCATCGAAGTCATCATGC | A/G (61,685) | PCR |
| jh_trnQ_W1-2 | I/I | ACGTTGGATGTCGCGAACTTTATACTCCAC | | PCR |
| jh_trnQ_W1UEP | I/I | TGAAGGCATCAGACCAT | | Extension |
| jh_trnS_W1-1 | I/I | ACGTTGGATGCGGATCGTTTGATTCATTTC | T/G (20,001) | PCR |
| jh_trnS_W1-2 | I/I | ACGTTGGATGATTTAGGTGGTTCCCCACCC | | PCR |
| jh_trnS_W1UEP | I/I | CCCACCCTTTTTCTTTC | | Extension |
| jh_ycf1-2_W1-1 | I/I | ACGTTGGATGTCCGAGCTGAGCAAGTTGTG | C/T (135,506) | PCR |
| jh_ycf1-2_W1-2 | I/I | ACGTTGGATGTATGCTTCCTGCATAGCTCG | | PCR |
| jh_ycf1-2_W1UEP | I/I | CGCAGCTTCGATTTCCAT | | Extension |
| jh_ycf1-3_W1-1 | I/I | ACGTTGGATGCTATCGAACGAAACCCAAGC | G/T (135,768) | PCR |
| jh_ycf1-3_W1-2 | I/I | ACGTTGGATGCTCCTTTGCGCTTCGAATTT | | PCR |
| jh_ycf1-3_W1UEP | I/I | CTTTCTTCAAGTTTCTTCTCT | | Extension |
| ja_atpH_W1-1 | I/IV | ACGTTGGATGTTCTTAGCAACGGAAACCCC | A/G (52,971) | PCR |
| ja_atpH_W1-2 | I/IV | ACGTTGGATGTAGAATCTAGGGCGGGTTTC | | PCR |
| ja_atpH_W1UEP | I/IV | CTTGTACCAATCCCTAAAAAAA | | Extension |
| ja_ndhD-1_W1-1 | I/IV | ACGTTGGATGAGCGATACGGGACTTAATGG | G/T (121,802) | PCR |
| ja_ndhD-1_W1-2 | I/IV | ACGTTGGATGCCGGCCAAGAAAAAAAGAGC | | PCR |
| ja_ndhD-1_W1UEP | I/IV | AGAGCCGCCCCAATAAA | | Extension |
| ja_ndhD-2_W1-1 | I/IV | ACGTTGGATGAAACCAATACCCAGAACTCC | A/C (122,300) | PCR |
| ja_ndhD-2_W1-2 | I/IV | ACGTTGGATGCGTCTCTATTCAGCTACAAAG | | PCR |
| ja_ndhD-2_W1UEP | I/IV | AAAGTTCATTTTGTACACGGCGGG | | Extension |

**Table S12.** (continued)

| Primer name | Assay | Sequence (5‘ - 3‘) | SNP (position in I-johSt) | Function |
|---|---|---|---|---|
| ja_ndhH_W1-1 | I/IV | ACGTTGGATGTATTCAGCAGGCTCTGGAAG | C/A (127,723) | PCR |
| ja_ndhH_W1-2 | I/IV | ACGTTGGATGTTCAAAACCGTCCCATTCTG | | PCR |
| ja_ndhH_W1UEP | I/IV | CCATTCTGGATCCTTTTCT | | Extension |
| ja_ndhJ_W1-1 | I/IV | ACGTTGGATGACGTTCCCAATGTGCTTATG | T/C (15,205) | PCR |
| ja_ndhJ_W1-2 | I/IV | ACGTTGGATGGCTTGATCCACACCATACTC | | PCR |
| ja_ndhJ_W1UEP | I/IV | ACACTAGCTAACAGTCC | | Extension |
| ja_petD-1_W1-1 | I/IV | ACGTTGGATGCACACCTACATGAATGAATCC | G/A (82,065) | PCR |
| ja_petD-1_W1-2 | I/IV | ACGTTGGATGAAATTCCTTGCAATGGGTAG | | PCR |
| ja_petD-1_W1UEP | I/IV | AATGGGTAGTTGCAACTGC | | Extension |
| ja_psaA_W1-1 | I/IV | ACGTTGGATGGCCCTGCTAAATGGTGATTC | G/T (23,840) | PCR |
| ja_psaA_W1-2 | I/IV | ACGTTGGATGGCTTTTTGCTGGTTGGTTCC | | PCR |
| ja_psaA_W1UEP | I/IV | CTGGTTGGTTCCATTATCACAAAGC | | Extension |
| ja_psaB_W1-1 | I/IV | ACGTTGGATGAGATGTATTACCGCATCCCG | A/C (26,244) | PCR |
| ja_psaB_W1-2 | I/IV | ACGTTGGATGGGCATAAAGATTCCACTGGC | | PCR |
| ja_psaB_W1UEP | I/IV | TGGCCTGTAAAAAGGGG | | Extension |
| ja_rpl116_W1-1 | I/IV | ACGTTGGATGAACCCCGAATATTGGGTAGC | T/G (86,534) | PCR |
| ja_rpl116_W1-2 | I/IV | ACGTTGGATGTATTTTCTGCGACTCCACCC | | PCR |
| ja_rpl116_W1UEP | I/IV | ACTCCACCCATTTCATAAAG | | Extension |
| ja_rpoB_W1-1 | I/IV | ACGTTGGATGAGCGCGAACTGCTAGTTTTC | T/C (40,519) | PCR |
| ja_rpoB_W1-2 | I/IV | ACGTTGGATGCTTCAGACGTGTCAATTGGG | | PCR |
| ja_rpoB_W1UEP | I/IV | GTCAATTGGGCAAATACGTCCATAGT | | Extension |
| ja_rps4_W1-1 | I/IV | ACGTTGGATGTTTATGTCGCGTTACCGAGG | C/A (18,825) | PCR |
| ja_rps4_W1-2 | I/IV | ACGTTGGATGGGCTCTAGGCCTTTTACTGG | | PCR |
| ja_rps4_W1UEP | I/IV | GCCTTTTACTGGTTAGTCC | | Extension |
| ja_trnW_W1-1 | I/IV | ACGTTGGATGACTGAACTAAGAGCGCTTTC | A/G (72,243) | PCR |
| ja_trnW_W1-2 | I/IV | ACGTTGGATGAGCATACAAGAGGTATTGGG | | PCR |
| ja_trnW_W1UEP | I/IV | TTGGGATTACAAAACAAAAGA | | Extension |
| ja_ycf1_W1-1 | I/IV | ACGTTGGATGGCAATTCTCTACGCCGTTTG | G/A (129,698) | PCR |
| ja_ycf1_W1-2 | I/IV | ACGTTGGATGTATGCCTAAGACCAATGCGG | | PCR |
| ja_ycf1_W1UEP | I/IV | AGTGATTCTGATTTGTTTGTTCC | | Extension |
| ja_ycf2_W1-1 | I/IV | ACGTTGGATGCGAAAGAACGGAACTTAGCC | C/A (94,219; 161,260) | PCR |
| ja_ycf2_W1-2 | I/IV | ACGTTGGATGATTCGAGTGATCGGTCTGAG | | PCR |
| ja_ycf2_W1UEP | I/IV | CGGTCTGAGGTTAGCGAC | | Extension |
| jah_atpF_W1-1 | I/IV | ACGTTGGATGGGGCTTTTTCCAGCTGTTCG | T/C (54,206) | PCR |
| jah_atpF_W1-2 | I/IV | ACGTTGGATGATCGAAAACAGAGGATCTTG | | PCR |
| jah_atpF_W1UEP | I/IV | GAGGATCTTGAATACTATTCG | | Extension |

**Table S13.** Summary of F. Schötz's original transmission frequencies in hybrid populations of *Oenothera*, from which the different classes of inheritance strength were determined (*6*)[1)]

| Species | Strain | Abbreviation plastome | Basic plastome[2)] | [%] biparental inheritance (biennis white)[3)] | [%] biparental inheritance (blandina white)[3)] | Inheritance class (Schötz)[4)] | Inheritance class (this work)[5)] |
|---|---|---|---|---|---|---|---|
| *O. elata* ssp. *hookeri* | hookeri de Vries | I-hookdV | I | 0.0 | 8.2 | strong | 1 |
| *O. elata* ssp. *hookeri* | franciscana de Vries | I-frandV | I | 0.8 | 19.9 | strong | 1 |
| *O. villosa* ssp. *villosa* | cockerelli de Vries | I-cockdV | I | 0.0 | 9.9 | strong | 1 |
| *O. villosa* ssp. *villosa* | mollis Standard | I-molSt | I | 1.1 | 31.5 | strong | 1 |
| *O. villosa* ssp. *villosa* | bauri Standard | I-bauriSt | I | 7.4 | 43.8 | strong | 3 |
| *O. glazioviana* | blandina de Vries | III-blandV | III | 0.0 | 14.1 | strong | 1 |
| *O. glazioviana* | deserens de Vries | III-desdV | III | 1.3 | 21.4 | strong | 1 |
| *O. glazioviana* | *r*/*r*-lamarckiana Sweden | III-lamS | III | 3.1 | 26.4 | strong | 1 |
| *O. glazioviana* | coronifera Standard | II-corSt | II | 38.0 | 57.7 | intermediate | 4 |
| *O. biennis* x *O. glazioviana* | conferta Stanard | II-conSt | II | 16.1 | 34.2 | intermediate | 3 |
| *O. biennis* x *O. villosa* ssp. *villosa* | hoelscheri Standard | II-hoeSt | II | 31.1 | 57.5 | intermediate | 3 |
| *O. biennis* | chicaginensis de Vries | III-chicdV | III | 5.0 | 24.4 | strong | 1 |
| *O. biennis* | nuda Standard | II-nudaSt | II | 12.4 | 54.6 | intermediate | 3 |
| *O. biennis* | purpurata Standard | II-purSt | II | 14.9 | 45.0 | intermediate | 3 |
| *O. biennis* | biennis Muenchen | II-biM | II | 22.2 | 53.8 | intermediate | 3 |
| *O. biennis* | rubricaulis Thorn | II-rcauTh | II | 23.2 | 41.8 | intermediate | 3 |
| *O. biennis* | suaveolens Standard | II-suavSt | II | 26.7 | 49.9 | intermediate | 3 |
| *O. biennis* | grandiflora de Vries | II-gradV | II | 28.3 | 53.5 | intermediate | 3 |
| *O. oakesiana* | ammophila Standard | IV-ammSt | IV | 43.9 | 58.1 | weak | 4 |
| *O. oakesiana* | *r*/*r*-syrticola Ulm | IV-syrtU | IV | 48.7 | 63.3 | weak | 4 |
| *O. oakesiana* | germanica Standard | IV-gerSt | IV | 52.9 | 74.5 | weak | 4 |
| *O. parviflora* | parviflora Waldenburg | IV-parW | IV | 47.6 | 63.2 | weak | 4 |
| *O. parviflora* | rubricuspis Standard | IV-rcuSt | IV | 52.3 | 68.7 | weak | 4 |
| *O. parviflora* | silesiaca Standard | IV-silSt | IV | 53.2 | 67.2 | weak | 4 |
| *O. parviflora* | atrovirens Standard | IV-atroSt | IV | 70.1 | 72.6 | weak | 4 |

[1)]For reviews see (24, 68, 87).

[2)]In *Oenothera*, five genetically distinguishable plastome types (I-V) can be recognized based on their compatibility with three nuclear genomes (A, B, C) in either homozygous (AA, BB, CC) or stable heterozygous (AB, AC, BC) states. The basic plastome genotype is accompanied by a given inheritance strength (strong, intermediate and weak). Basic plastome and nuclear genome type are important factors of species definition in *Oenothera*. For details, see e.g. (24, 34, 60, 68, 81, 85, 86)

[3)]Percentage of variegated seedlings, heteroplasmic due to the paternal transmission of the bleached chloroplast mutants "biennis white" or "blandina white" and the maternal transmission of the green wild chloroplast of the seed parent. Data accoding to Schötz (34). For details see therein, reviews in (24, 68) and SI Text.

[4)]Class of inheritance strength as determined by F. Schötz. For reviews see (24, 68, 87)

[5)]For details on the definition of these classes see SI Text.

**Table S14.** Nucleoids per chloroplasts in plants with the strong plastome I-johSt vs. the weak plastomes VC1, V3g, and IV-atroSt[1)]

| Variant/ plastome[2)] | # cells analyzed | # chloroplasts analyzed | # nucleoids counted | # nucleoids/ chloroplast[3)] | One-way ANOVA *p*-value | Adjusted *p*-value[4)] |
|---|---|---|---|---|---|---|
| I-johSt | 22 | 266 | 4,703 | 17.7 ± 6.8 | 0.53 | n/a |
| VC1 | 22 | 435 | 7,953 | 18.3 ± 5.2 | 0.53 | 0.32 |
| V3g | 22 | 418 | 7,625 | 18.2 ± 4.9 | 0.53 | 0.32 |
| IV-atroSt | 22 | 322 | 5,839 | 18.1 ± 5.7 | 0.53 | 0.38 |

[1)]All analyses were performed in the constant nuclear genetic background of *O. elata* ssp. *hookeri* strain johansen Standard. For details on the lines see Table S6.

[2)]For details on the wild type or variant plastomes see Table S10.

[3)]Means ± standard deviations are given.

[4)]*p*-values obtained with two-tailed homoscedastic *t*-test followed by multiple testing correction according to Benjamini-Hochberg.

**Table S15.** Chloroplasts per cell in plants with the strong plastome I-johSt vs. the weak plastomes VC1, V3g, and IV-atroSt[1)]

| Variant/ plastome[2)] | # cells analyzed | # chloroplasts counted | # chloroplasts/ cell[3)] | One-way ANOVA *p*-value | Adjusted *p*-value[4)] |
|---|---|---|---|---|---|
| I-johSt | 55 | 2,872 | 52.2 ± 10.3 | 0.93 | n/a |
| VC1 | 55 | 2,920 | 53.1 ± 10.2 | 0.93 | 0.66 |
| V3g | 55 | 2,918 | 53.1 ± 9.2 | 0.93 | 0.66 |
| IV-atroSt | 55 | 2,935 | 53.4 ± 7.7 | 0.93 | 0.66 |

[1)]All analyses were performed in the constant nuclear genetic background of *O. elata* ssp. *hookeri* strain johansen Standard. For details on the lines, see Table S6.

[2)]For details on the wild type or variant plastomes see Table S10.

[3)]Means ± standard deviations are given.

[4)]*p*-values obtained with two-tailed homoscedastic *t*-test followed by multiple testing correction according to Benjamini-Hochberg.

**Table S16.** Comparison of mesophyll cell chloroplasts with the strong plastome I-johSt vs. the weak ones VC1, V3g, and IV-atroSt[1)]

| **Variant/ plastome[2)]** | **# chloroplasts analyzed** | **Chloroplast volume index [length x width$^2$][3,5)]** | **One-way ANOVA *p*-value** | **Adjusted *p*-value[4)]** | **Chloroplast length [µm][3)]** | **One-way ANOVA *p*-value** | **Adjusted *p*-value[4)]** | **Chloroplast width [µm][3)]** | **One-way ANOVA *p*-value** | **Adjusted *p*-value[4)]** |
|---|---|---|---|---|---|---|---|---|---|---|
| I-johSt | 60 | 86.8 ± 27.7 | 0.51 | n/a | 5.4 ± 0.6 | 0.66 | n/a | 4.0 ± 0.5 | 0.25 | n/a |
| VC1 | 60 | 78.7 ± 31.6 | 0.51 | 0.42 | 5.3 ± 0.6 | 0.66 | 0.35 | 3.8 ± 0.6 | 0.25 | 0.20 |
| V3g | 60 | 81.2 ± 31.0 | 0.51 | 0.46 | 5.3 ± 0.7 | 0.66 | 0.35 | 3.8 ± 0.5 | 0.25 | 0.41 |
| IV-atroSt | 60 | 85.1 ± 36.6 | 0.51 | 0.78 | 5.3 ± 0.7 | 0.66 | 0.35 | 3.9 ± 0.6 | 0.25 | 0.81 |

[1)]All analyses were performed in the constant nuclear genetic background of *O. elata* ssp. *hookeri* strain johansen Standard. For details on the lines see Table S6.

[2)]For details on the wild type or variant plastomes see Table S10.

[3)]Means ± standard deviations are given.

[4)]*p*-values obtained with two-tailed homoscedastic *t*-test followed by multiple testing correction according to Benjamini-Hochberg.

[5)]Chloroplast volume index calculated according to (20).

**Table S17.** Oligonucleotides used for real-time qPCR.

| Name | Genome | Gene/locus | Sequence (5‘ - 3‘) | Primer efficiency | Amplification product [bp] | GenBank/EMBL accession number template |
|---|---|---|---|---|---|---|
| rbcL-gene-SH_for | plastid | *rbcL* | TGCAGTAGCTGCCGAATCTTCTACTG | 1.97 | 122 | AJ271079.4 |
| OerbcLqPCR_rev2 | plastid | *rbcL* | GATTCTCTTCTCCCGCAACAGGCT | | | EU262891.2 |
| psbB_qPCR_for | plastid | *psbB* | GTGGAGACAAGGTATGTTCGTTATAC | 1.92 | 125 | AJ271079.4 |
| psbB_qPCR_rev2 | plastid | *psbB* | CCACACCTTCGTAACTCCAAA | | | EU262891.2 |
| Oe_ndhI_qPCR-fw | plastid | *ndhI* | CTGACATTGGTAAACGACCCAAAGC | 1.94 | 126 | AJ271079.4 |
| Oe_ndhI_qPCR-rev | plastid | *ndhI* | GTGGTAACTGCGTTGAGTATTGTCC | | | EU262891.2 |
| mtM03_qPCR_for | mitochondrion | mtM03 | TCGCTTTCTTCGTCCACCAG | 1.88 | 184 | n/a |
| mtM03_qPCR_rev | mitochondrion | mtM03 | CGATTCGCTTAGTTCGGTAGGGT | | | |
| mtM04qTAG_for | mitochondrion | mtM04 | TTATTTTTGCAGTGTGGCTCCTAA | 1.87 | 118 | n/a |
| mtM04qTAG_rev | mitochondrion | mtM04 | CCCAGCGAGAACGCTACCT | | | |
| mtM06qTAG_for | mitochondrion | mtM06 | GGAAGTCAAGCTAAAGCAGTCTAAGC | 1.89 | 93 | n/a |
| mtM06qTAG_rev | mitochondrion | mtM06 | CTCTCATTAGCTCTCCTTTTTTTATATGC | | | |
| M02-1_qPCR-Fw | nucleus | M02 | CACAAGCCTCCATCTTTACTCCGT | 1.94 | 177 | EU483117.1 |
| M02-1_qPCR-Rev | nucleus | M02 | GCTACCTTCTCCTTGGTAGGCG | | | |
| M19_qPCR-Fw | nucleus | M19 | GTGAAGCTCTCCCCTGAGGT | 1.92 | 82 | EU447207.1 |
| M19_qPCR-Rev | nucleus | M19 | ATGAGACAGCGGGTTTGGC | | | |
| Mpgi02_qPCR-Fw | nucleus | *pigC* | CAGGTCTGTACTACATGTCGCTCT | 1.97 | 141 | GU176508.1 |
| Mpgi02_qPCR-Rev | nucleus | *pigC* | ACCCATGAACCATTGCGAACT | | | |
